# CRISPR-Cas phage defense systems and prophages in *Candidatus* Accumulibacter

**DOI:** 10.1101/2022.10.12.504627

**Authors:** Xuhan Deng, Jing Yuan, Liping Chen, Hang Chen, Chaohai Wei, Per H. Nielsen, Stefan Wuertz, Guanglei Qiu

**Affiliations:** School of Environment and Energy, South China University of Technology, Guangzhou 510006, China.; Singapore Centre for Environmental Life Sciences Engineering, Nanyang Technological University, Singapore 637551, Singapore.; School of Civil and Environmental Engineering, Nanyang Technological University, Singapore 639798, Singapore; Centre for Microbial Communities, Department of Chemistry and Bioscience, Aalborg University, DK-9220, Aalborg, Denmark.

**Keywords:** *Candidatus* Accumulibacter, EBPR system, CRISPR-Cas systems, prophages, Caudovirales

## Abstract

*Candidatus* Accumulibacter is a key genus of polyphosphate-accumulating organisms (PAOs) found in laboratory- and full-scale wastewater treatment systems, mediating enhanced biological phosphorus removal (EBPR). However, little is known about their ability to resist phage infection. We conducted a systematic analysis of the occurrence and characteristics of clustered regularly interspaced short palindromic repeats and associated proteins (CRISPR-Cas) systems and prophages in diverse *Ca.* Accumulibacter taxa (43 in total, including 10 newly recovered genomes). Fourty complete CRISPR loci were identified in 28 genomes, falling into 6 subtypes. The occurrence and distribution of CRISPR-Cas systems did not follow a vertical evolutionary relationship. Phylogenetic analyses of the *cas* genes and direct repeats (DRs) suggested that the CRISPR-Cas systems were likely acquired via horizontal gene transfer, with acquisition rates higher than those of speciation, rendering different *Ca.* Accumulibacter distinct adaptivity to phage predations. 2448 spacers were identified, 67 of them matched to known phages. Major differences were observed among the numbers of spacers for different *Ca.* Accumulibacter, showing unique phages that could be resisted by different members. A comparison of the spacers in genomes having the same *cas* gene but from distinct geographical locations indicated that habitat isolation may have resulted in the acquisition of different spacers by different *Ca*. Accumulibacter. Metagenomic analysis allowed the identification of 26 viral contigs (18 are Caudovirales members) in 6 metagenomic datasets from three lab-scale enrichment reactors, matching to 73 spacers in 10 *Ca.* Accumulibacter genomes in these reactors, showing the specific immunity of these *Ca.* Accumulibacter. Metatranscriptomic analyses showed the activity of the CRISPR-Cas system under both anaerobic and aerobic conditions. Extra efforts were made to identify prophages in *Ca.* Accumulibacter genomes. In total, 133 prophage regions were identified. Twenty-seven of them were classified as potentially active prophages. Three prophages (all are Caudovirales members, in DS2011, SCELSE-7IIH and SCELSE-5IIH, respectively) are readily activable. Differences in the occurrence of CRISPR-Cas systems and prophages in *Ca.* Accumulibacter will lead to their distinct responses under phage predation. This study represents the first systematic analysis of CRISPR-Cas systems and prophages with combined experimental and bioinformatic methods in the *Ca.* Accumulibacter lineage, providing new perspectives on the potential impacts of phages on *Ca.* Accumulibacter and EBPR systems.

## 1. Introduction

Phosphorus (P) is a pollutant in surface waters and also a non-renewable resource (Oehmen et al., 2007). Enhanced biological phosphorous removal (EBPR) is a widely applied process in municipal wastewater treatment plants (WWTPs) (Chen et al., 2022; Mino et al., 1998); it requires polyphosphate accumulating organisms (PAOs) capable of releasing P under anaerobic, and taking up more P than normally required for metabolism and growth, under aerobic conditions (He and McMahon, 2011; Lu et al., 2021; Zhang et al., 2022). *Candidatus* Accumulibacter represents a key group of PAOs in lab- and full-scale EBPR systems (Martín et al., 2006; Petriglieri et al., 2022; Qiu et al., 2020; Singleton et al., 2022; Srinivasan et al., 2021; Tomás-Martínez et al., 2021). Ensuring the viability of *Ca.* Accumulibacter is crucial for reliable P removal in WWTPs. However, even in well-controlled operating conditions, EBPR systems frequently experience deterioration or failure (Ong et al., 2014; Thomas et al., 2003). The overgrowth of glycogen accumulating organisms (GAOs) was commonly considered as a key cause, which competing with PAOs for carbon sources (such as volatile fatty acids, VFAs) without contributing to P removal (Kolakovic et al., 2021; McIlroy et al., 2014; Wang et al., 2021). However, GAOs might not well-explain EBPR deterioration in a variety of systems(Gu et al., 2008; Law et al., 2016; Nielsen et al., 2019). Even in lab-scale systems, non-GAO induced EBPR failure was commonly observed. In a lab-scale EBPR system operated at elevated temperate, a sudden drop in the EBPR activity (P release and uptake decreased from 40 to 0 mg/L within a day) was observed. The relative abundances of both PAOs and GAOs decreased significantly, suggesting that the affected EBPR activity was not a result of GAO competition (Qiu et al., 2022).

Phages are viruses specialized in infecting bacteria, which play a major role in shaping the microbial communities in natural and engineered systems (Fernández et al., 2018). In WWTPs, phages affect functional microbial communities by infection, gene transfer, and lysogenic conversion (Tamaki et al., 2012; Wu and Liu, 2009). A previous study suggested that phages in activated sludge number on the order of 10^8^-10^9^ ml^-1^ (Chen et al., 2021; Li et al., 2021), that is, their density is 10-1000 times higher than in natural aquatic environments (10^4^-10^8^ ml^-1^) (Wommack and Colwell, 2000). Such high occurrences of phages imply intensive interactions with activated sludge bacteria. Like other bacteria, *Ca.* Accumulibacter are susceptible to phage predation. In a study on a deteriorated laboratory-scale EBPR system, large quantities of lysed *Ca*. Accumulibacter cells were observed, and phages isolated from the supernatant were used to inoculate other sequencing batch reactors (SBRs), resulting in *Ca*. Accumulibacter cell lysis and a significant decline in P removal (Barr et al., 2010).

A key mechanism for bacteria to resist phage predation is the clustered regularly interspaced short palindromic repeats and their associated protein (CRISPR-Cas) system, consisting of a CRISPR array and a few *cas* genes (Barrangou et al., 2007). The immune process of the CRISPR-Cas system works in three stages: adaption, CRISPR RNA (crRNA) biogenesis, and interference (Fig. S1). At the adaption stage, the CRISPR-Cas system captures foreign nucleic acid (such as phage genetic material and plasmids) and integrates a fragment (about 30 base pairs) into the CRISPR array as spacers to provide infection memory. At the crRNA biogenesis stage, CRISPR repeat-spacer arrays are transcribed and processed into crRNAs. Each crRNA contains a conserved direct repeat (DR) and a variable spacer. In the infection stage, the interference machinery is guided by crRNA to identify and cleave foreign nucleic acids (Hille et al., 2018). This immunity mechanism is widely preserved in the bacterial and archaeal kingdoms (Barrangou et al., 2007). Phage predation is a threat to *Ca.* Accumulibacter and EBPR, and CRISPR-Cas systems are essential for *Ca*. Accumulibacter to resist phages (Flowers et al., 2013). However, little is known about this defense system in different lineage members of *Ca.* Accumulibacter.

Apart from lysing the host cells immediately, phages have another infection mode, namely lysogeny. Under such circumstance, temperate phages integrate their genomes into the host bacteria chromosome and become prophages (Feiner et al., 2015), which replicate with their host and stay at an inactive state. They may be activated with or without external stimulations (such as UV light, chemicals, or temperature) (Barnhart et al., 1976; Shan et al., 2014) to enter the lytic cycle and kill their host. Prophages in *Ca.* Accumulibacter genomes are timebombs which might be triggered at times, threatening the reliability of EBPR. Limited information is available on prophages in existing *Ca.* Accumulibacter genomes.

In this study, CRISPR-Cas systems and prophages in different *Ca.* Accumulibacter lineage members were systematically investigated to understand their capability in resisting phage predations. The objectives were to (i) understand the adaptivity of different lineage members of *Ca.* Accumulibacter to phage predictions; (ii) analyze the temporal and spatial variation of their ability in resisting phages; (iii) identify and characterize phages and prophages which are highly threatening to different *Ca.* Accumulibacter; and (iv) characterize the activity of CRISPR-Cas systems *Ca.* Accumulibacter in EBPR cycles. This work represents the first detailed study of the CRISPR-Cas systems and prophages in different *Ca.* Accumulibacter lineage members. The results are expected to lay a fundamental base for the study of interactions between *Ca.* Accumulibacter and their targeting phages, benefiting in understanding phage-mediated inter-species interactions among different *Ca.* Accumulibacter lineage members, and the potential threats of phages to *Ca.* Accumulibacter and their mediated EBPR for the development of countermeasures.

## 2. Material and Methods

### 2.1. SBR reactor setup and operation

Two SBRs (effective volume of 1.59 L) were operated in parallel for the enrichment of *Ca.* Accumulibacter at 30 °C (named NTU30) and 35 °C (named NTU35), respectively (Qiu et al., 2022). These two reactors were inoculated with activated sludge from a WWTP in Singapore (details on operational conditions are available in the *Supplementary Materials*). Activated sludge samples (12 in total) were collected from two SBRs on Days 14, 56, 91, 201, 280, and 301 for metagenomic analysis. Another 4.5-L SBR (named SCUT) was used for the enrichment of *Ca.* Accumulibacter at 25 °C and inoculated with activated sludge from a WWTP in Guangzhou, China. An activated sludge sample was collected on Day783 for metagenomic analysis. An anaerobic-aerobic full cycle study was performed to investigate the gene transcription activity of CRISPR-Cas systems in the SCUT reactor. Six activated sludge samples were collected during this full cycle study for metatranscriptomic analysis (more details on reactor operation, sample collection, and metatranscriptomic sequencing are found in the *Supplementary Materials*).

### 2.2. Metagenomic and metatranscriptomic analysis

After DNA and RNA extraction, metagenomic and metatranscriptomic sequencing were performed (detailed in the *Supplementary Materials*). For metagenomic analysis, low-quality reads and adapter sequences were removed using fastp (0.20.1) (Chen et al., 2018). High-quality reads were assembled to contigs using metaSPAdes (v.3.13.0) (Nurk et al., 2017). The contigs were binned into metagenome-assembled genomes (MAGs) using MetaBAT2(v1.7) and CONCOCT (v1.1) (Alneberg et al., 2014; Kang et al., 2019). Completeness and contamination of MAGs were evaluated by checkM (v1.0.18) (Parks et al., 2015). The taxonomic classifications of obtained MAGs were performed using the Genome Taxonomy Database Toolkit (GTDB-Tk v. 1.1.0) (Chaumeil et al., 2019). MAGs with completeness below 95% and contamination over 6% were discarded. The relative abundance of each *Ca.* Accumulibacter were calculated using BBMap (v38.96) (Bushnell et al., 2017). All draft genomes were annotated using RAST (V2.0) (Brettin et al., 2015). Except for quality control by fastp, annotation by RAST and relative abundance calculation by BBMap, all analyses were processed on the KBase platform (Arkin et al., 2018).

For metatranscriptomics analysis, fastp and SortMeRNA were employed to remove adapter sequences and rRNA (Chen et al., 2018; Kopylova et al., 2012). Reads that passed filtering were mapped to the respective *Ca.* Accumulibacter draft genome (retrieved via metagenomic sequencing) using BBMap (v38.96) (Bushnell et al., 2017). The gene transcriptions were then calculated using HTseq (v2.0.1) and normalized to read per kilo base per million (RPKM) (Mortazavi et al., 2008).

Raw reads and draft genomes obtained for the NTU30 and NTU35 bioreactors were submitted to NCBI under the BioProject No. PRJNA807832. Those obtained for the SCUT bioreactor were submitted under the BioProject No. PRJNA771771.

### 2.3. GenBank Data acquisition and evaluation

Seventy-one *Ca*. Accumulibacter genomes (representing all available *Ca*. Accumulibacter genomes in the GenBank database) were retrieved from the NCBI database. The completeness and contamination of the genomes were evaluated using CheckM (Parks et al., 2015). Genomes with completeness of less than 95% and contamination exceeding 6% were discarded. The remaining *Ca.* Accumulibacter genomes were then used for subsequent analysis.

### 2.4. CRISPR-Cas system identification in *Ca.* Accumulibacter

For each selected genome (43 in total, including 33 genomes retrieved from GenBank database, 10 newly recovered from our lab-scale reactor), CRISPRCasFinder was used to identify the CRISPR locus (Couvin et al., 2018). Only those with evidence levels of four and above were considered in this study (Jiang et al., 2020). Orphan CRISPR arrays (without *cas* genes) were ignored due to their inability to silence foreign genetic materials (Jiang et al., 2020). To ensure the accuracy of *cas* gene identification, RAST was employed for genome annotation (Brettin et al., 2015). The identified CRISPR loci were further checked and confirmed manually. The types of CRISPR system were determined based on the features of *cas* genes (Makarova et al., 2020). Spacers and DRs in each CRISPR array were extracted using CRISPRDetect for comparative analysis (Biswas et al., 2016).

### 2.5. Comparative genomics analysis of the CRISRPR-Cas system

The species-level phylogenetic tree was built using SpeciesTreeBuilder (v.0.1.3). The tree was visualized and annotated using iTOL V6 (Letunic and Bork, 2021). Average nucleotide identity (ANI) was calculated using pyani with the ANIb model (Pritchard et al., 2016).

Phylogenetic analysis of marker genes was employed to study the evolution of the CRISPR-Cas system in *Ca.* Accumulibacter by MEGA X (Stecher et al., 2020). For type I CRISPR-Cas systems, the *cas1* gene is highly conserved, making it a good marker (Makarova et al., 2015). Thus, *cas1* genes were used to establish the gene tree to analyze the phylogenetic relationships of different type I CRISPR-Cas systems. For type III CRISPR-Cas systems, the *cas1* gene is dispensable. *cas10* genes were used as a marker gene to establish the gene tree to understand the phylogenetic relationships among different type III systems.

DRs are conserved sequences occurring repeatedly in the CRISPR locus between adjacent spacers. They were used as markers to further understand the origination of the CRISPR-Cas systems in *Ca*. Accumulibacter. DR in each CRISPR locus were compared to the CRISPRCasdb database (a recently established database for the CRISPR-Cas system) (Couvin et al., 2018; Pourcel et al., 2019). Multiple sequence alignment and visualization of DRs in each *cas* gene clade were done using WebLogo (Crooks et al., 2004).

### 2.6. Spacer analysis in *Ca.* Accumulibacter

Spacers record the history of phage infections and show the identity of phages unable to infect the host bacteria. Spacer comparison and visualization were done using CRISPRStudio (Dion et al., 2018). They were compared to the IMG/VR database using the online comparison function provided by the database for the identification of their corresponding phages. IMG/VR database provided the largest collection of viral sequences covering both cultivated viruses and uncultured ones identified from metagenomes (Roux et al., 2020).

To analyze the relationships between *Ca.* Accumulibacter and phages in our SBR reactors, VirSorter (v2.2.3) (Guo et al., 2021) was used to identify viral contigs from the assembled metagenomic data which contained *Ca.* Accumulibacter. These viral contigs were used to build a phage genome dataset. The spacers in *Ca.* Accumulibacter genomes (retrieved from our SBRs) were compared to the phage genome dataset by BLASTn using the short-BLAST parameter (Altschul et al., 1997). The phage contigs which completely matched the spacers were compared to the IMG/VR database for taxonomic identification (Roux et al., 2020).

### 2.7. Prophage identification and analysis

PHASTER was used to identify the prophage-like elements in *Ca*. Accumulibacter genomes (Arndt et al., 2016). They were classified into intact (score>90), questionable (score 70-90) and incomplete prophages (the evaluation standards are detailed at https://phaster.ca/). Incomplete prophages typically lack essential functional genes, such as those related to entering the lytic cycle (e.g., holins and lysins). To further analyze the complete (including intact and questionable) prophages, all genes in the prophage elements were compared to the NR databases using NCBI BLASTn (Altschul et al., 1997). The PHASTER results and NR database annotation results were combined to tentatively subdivide prophage genes into functional modules (i.e., lysogeny, DNA replication, DNA packaging, tail module and lysis module) (Botstein, 1980) to understand their potential threats to the host *Ca.* Accumulibacter. To determine the identity of these prophages in *Ca.* Accumulibacter genomes, BLAST was employed to compare the prophage sequences with the IMG/VR database (Roux et al., 2020). The similarity between two prophages was analyzed using pyani to calculate the average nucleotide identity (ANI) (Pritchard et al., 2016).

## 3. Results and Discussion

### 3.1. Metagenome assembly

Fourty-four *Ca*. Accumulibacter genomes were recovered from three lab-scale SBRs (NTU35 and NTU30 (Qiu et al., 2022), and SCUT (Tian et al., 2022)). Ten high-quality (completeness >95%, contamination <6%) *Ca*. Accumulibacter genomes (Table S1) were selected for further analysis. All 10 genomes belong to clade II (3 clade IIC members, 5 clade IIF members, and 2 clade IIH members). This is the first study to report successful recovery of clade IIH member genomes.

### 3.2. GenBank genomes evaluation

Seventy-one *Ca.* Accumulibacter genomes were retrieved from the GenBank database and their qualities evaluated using checkM. Thirty-three had a completeness > 95% and contamination < 6% (Table S2), and they were selected for subsequent analysis. These genomes are distributed in 8 different clades (6 clade IA members, 3 clade IC members, 4 clade IIA members, 2 clade IIB members, 6 clade IIC members, 2 clade IID members, 5 clade IIF members, and 5 genomes with unknown clade identity). In total, 43 *Ca.* Accumulibacter genomes were used for CRISPR-Cas system and prophage identification and analysis (i.e., 10 were retrieved from our SBR reactors and 33 from the GenBank database). ANI results for pairs of *Ca.* Accumulibacter genomes are presented in Fig. 1A.

**Fig. 1.**
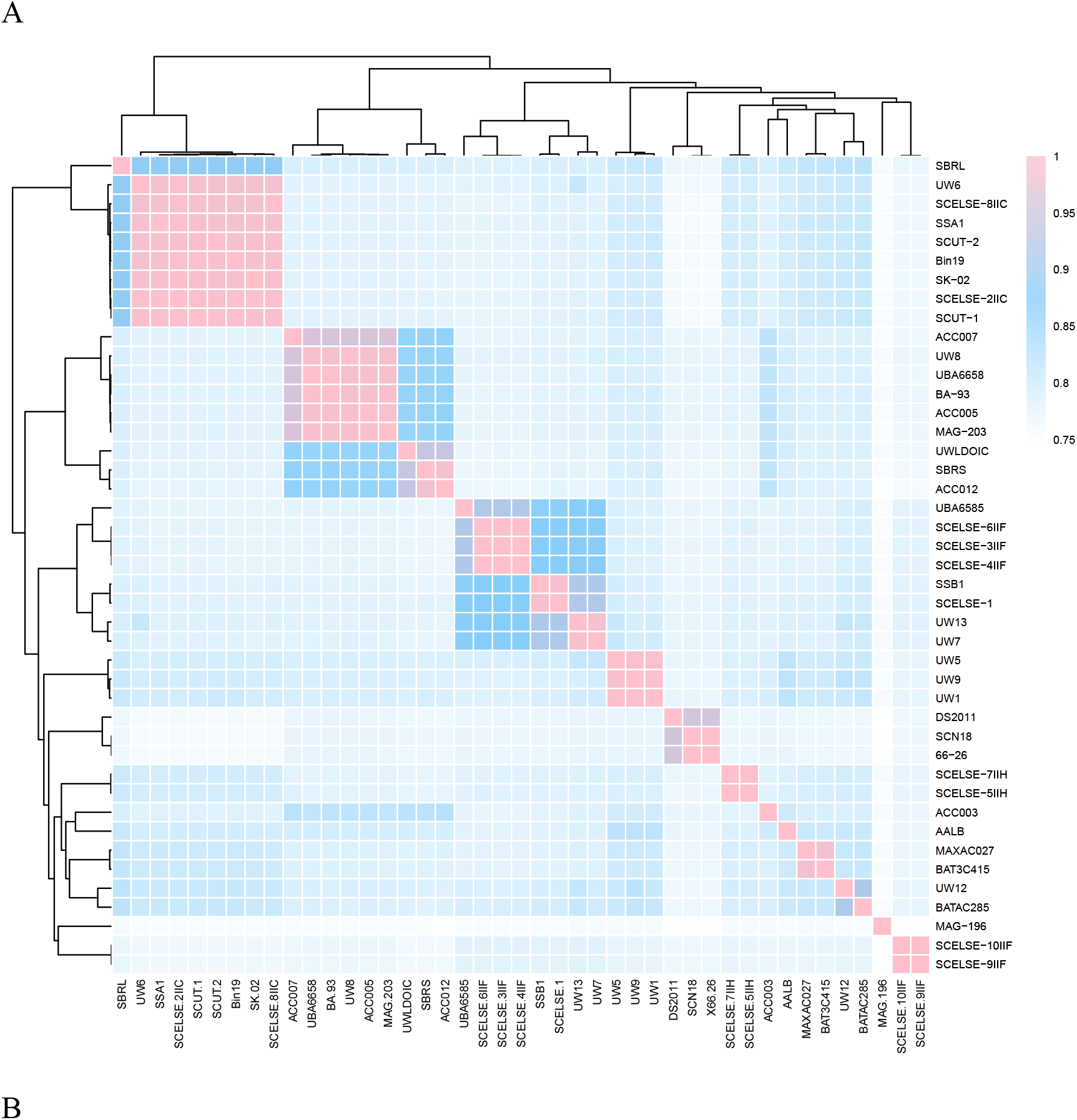
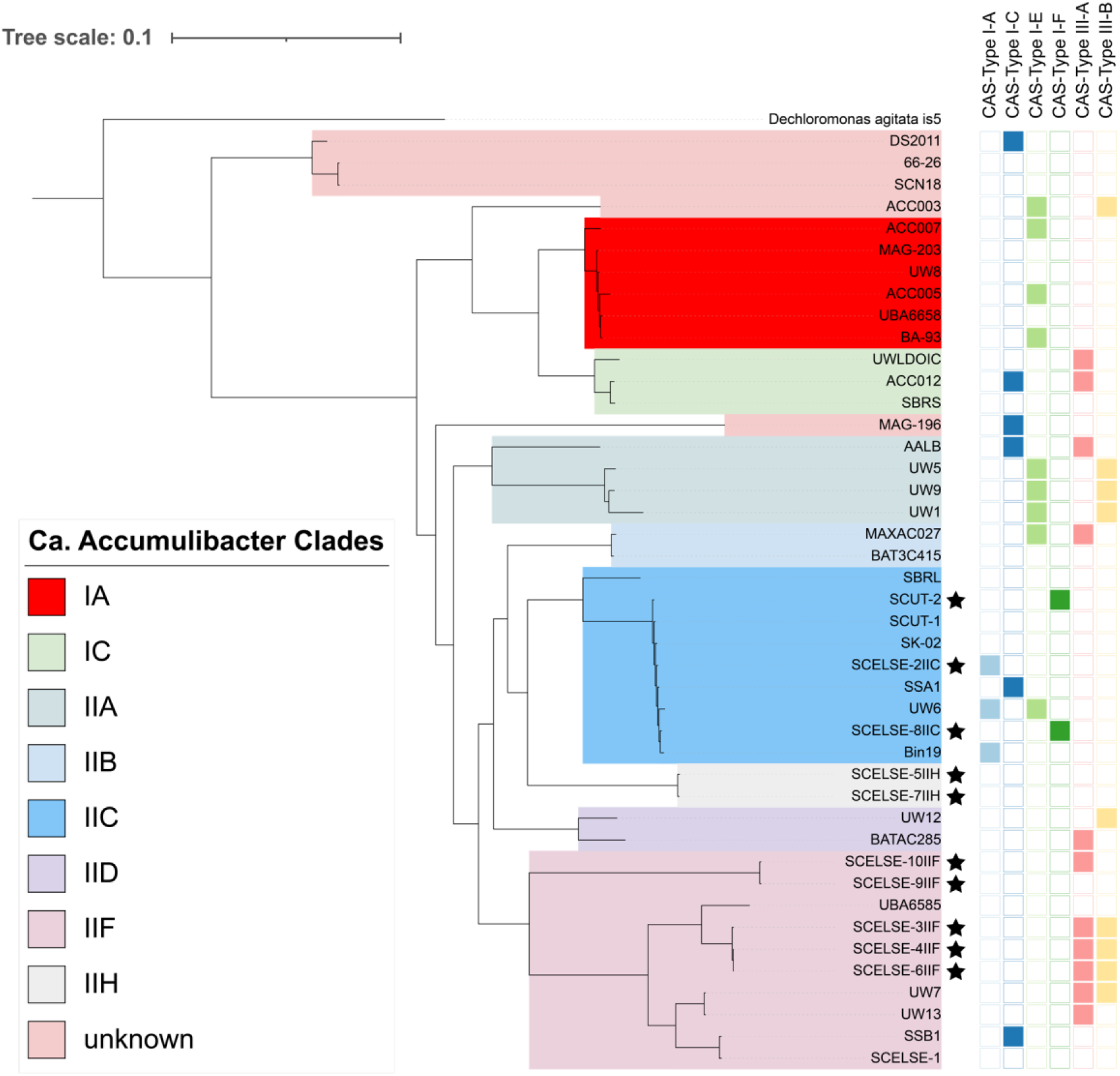
(A) Average nucleotide identity (ANI) analysis of *Ca.* Accumulibacter genomes. (B) Species tree of *Ca.* Accumulibacter and the occurrence and distribution CRISPR-Cas systems. *Dechloromonas agitate is5* is employed as an outgroup. The squares on the right of the tree represent different types of CRISPR-Cas systems. Solid squares indicate presence. The genomes with star marks were retrieved in this study.

### 3.3. High Occurrence of CRISPR-Cas system in *Ca*. Accumulibacter

Fourty CRISPR-Cas loci were found in 28 *Ca.* Accumulibacter genomes (Fig. 1B and Table S3). Fifteen genomes were not found to have a complete CRISPR-Cas system. These genomes are distributed in different clades (e.g., IA member UW8, IIC members SK-02 and SCUT-1, IIF members UBA6585 and SCELSE-1 (Qiu et al., 2020), and IIH member SCELSE-7). The overall occurrence of CRISPR-Cas systems in these *Ca.* Accumulibacter genomes was 65.1%, which is similar to the discovery rate in our newly recovered genomes (70%, 7 out of 10 had CRISPR-Cas systems). The rate is higher than the estimated average occurrence of CRISPR-Cas systems in bacteria (42.3%) (Makarova et al., 2020). There is consensus that CRISPR-Cas systems provide vital immunity to prokaryotes (Hille and Charpentier, 2016). The prevalence of CRISPR-Cas in *Ca.* Accumulibacter may indicate an overall higher capability to resist phage infections compared to other bacteria. Under pressure of phage-mediated natural selection, *Ca.* Accumulibacter taxa harboring CRISPR-Cas systems tend to have a higher chance to survive if most other *Ca.* Accumulibacter cells are killed. A few of the survivors obtain new spacers matching the genome of the invader. Should the same phage type return, these *Ca.* Accumulibacter could effectively resist. On the contrary, *Ca.* Accumulibacter without CRISPR-Cas systems may be more susceptible to phage invasion, affecting the reliability of the EBPR system.

Known CRISPR-Cas systems to date are classified into 2 classes, 6 types and 33 subtypes (Makarova et al., 2020). All CRISPR-Cas systems in *Ca.* Accumulibacter fall into Class 1, including 2 types (type I and type III) and 6 subtypes (3 type I-A, 6 type I-C, 9 type I-E, 2 type I-F, 11 type III-A and 9 type III-B, Fig. 1B). Type I system primarily targets double-stranded DNA (Lin et al., 2020). Type III system has both RNA and single-stranded DNA cleavage activities, which works against both the transcripts and genomes of invaders (Jiang et al., 2016). Previous studies hypothesized that type III systems could effectively counteract propagation of single-stranded RNA phage in *E. coli* (Tamulaitis et al., 2014). Overall, RNA phages are rare in the prokaryotic world (<10%) and spacers in type III system that match RNA viruses are yet to be identified (Koonin et al., 2015; Krupovic et al., 2011). Additionally, the mutation rates of RNA viruses are higher than those of DNA viruses (by three orders of magnitude), and maintaining long-term immunity against RNA viruses would require extremely rapid spacer renewal (Duffy, 2018). Of 28 *Ca*. Accumulibacter genomes with CRISPR-Cas systems, 12 harbored two CRISPR-Cas loci (e.g., clade IIA members UW1 and AALB, and clade IIF member SCELSE-6), and 16 had only one CRISPR-Cas locus (e.g., IIC members SCELSE-8, SCUT-2 and SSA1, IIF member SSB1, and IA member BA-93) (Fig.1B). In principle, *Ca.* Accumulibacter genomes with multiple CRISPR-Cas systems (i.e., UW1, AALB, SCELSE-6IIF) would have a higher capability of handling threats from different kinds of phages. However, the maintenance of multiple CRISPR-Cas systems may also exert extra energy and resource investment (Westra et al., 2015), which would only be advantageous in a highly stressed and dynamic environment.

Of the 10 newly recovered genomes, three do not harbor any CRISPR-Cas systems (Fig. 2A). The dynamics in their relative abundance in the lab-scale reactors was analyzed, showing that *Ca*. Accumulibacter with CRISPR-Cas systems typically exhibited high relative abundances (e.g., 37.1% for SCUT-2). *Ca.* Accumulibacter without CRISPR-Cas system are more likely to be washed out during long-term operation owing to their lower ability to resist phage predation. Inoculating an EBPR system with *Ca.* Accumulibacter strains harboring CRISPR-Cas systems could potentially enhance the system’s ability to resist phages and benefit stable EBPR.

**Fig. 2.**
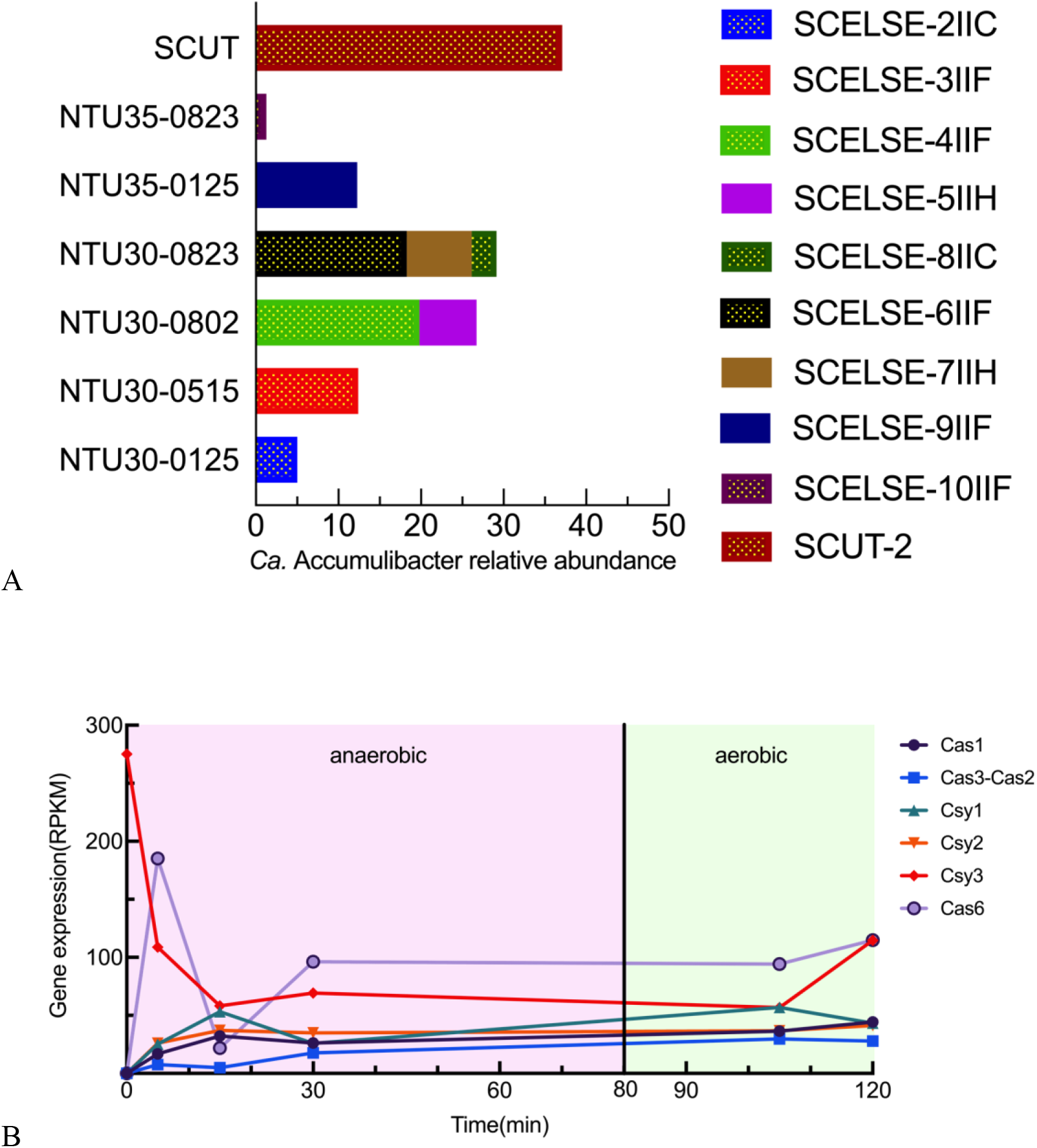
(A) The relative abundance of different *Ca*. Accumulibacter members retrieved from the lab-scale reactors. Bars with yellow dots indicate *Ca*. Accumulibacter genomes with CRISPR-Cas systems. The y-axis indicates the metagenomic dataset where the *Ca.* Accumulibacter genomes were recovered. (B) Transcription of CRISPR-associate genes in *Ca*. Accumulibacter SCUT-2, which harbors a type I-F CRISPR-Cas system. Gene transcription were normalized to read per kilo base per million (RPKM).

### 3.4. Activity of CRISPR-Cas systems

To investigate the activity of CRISPR-Cas systems in *Ca*. Accumulibacter, metatranscriptomic analyses were employed to analyze the transcription of CRISPR-Cas system in an anaerobic-aerobic full cycle (in the SCUT SBR, from which the draft genome of *Ca.* Accumulibacter SCUT-2 with a Type I-F CRISPR locus was recovered).

Results show that before the SBR cycle starts (0 min, Fig. 2B), the transcription of all *cas* genes except the *csy3* gene were low, suggesting that after going through sedimentation, discharge, and idle stages, the CRISPR-Cas system was largely not activity. The silence of *cas* genes saved energy. The high transcription of *csy3* (275.1 RPKM, Fig. 2B) may be a result of the need to form backbones for CRISPR-Cas effector complexes, which contain six Csy3 proteins in each complex (Majumdar et al., 2015; Xue and Sashital, 2019).

After the addition of the carbon source, all *cas* genes started transcription (5 min, Fig. 2B). Cas1 and Cas3-Cas2 proteins form an adaption complex, which capture and trim foreign DNA sequence and integrate a piece of them into the CRISPR array as an immune memory. The transcription of the *cas1* gene remained relatively consistent (32.2-44.3 RPKM) during the cycle, while the transcription of *cas3-cas2* increased 6.1 times after 30 min. Cas3-Cas2 serves as a structural scaffold for the adaption complex and a nuclease/helicase for the degradation of target dsDNA (Xue and Sashital, 2019). Unchanged transcription of the *cas1* gene suggested that Cas3-Cas2 proteins likely acted as a nuclease/helicase.

Cas6 is necessary in the generation of a mature crRNA (Makarova et al., 2020). Csy1, Csy2, Csy3, Cas3-Cas2 and Cas6 proteins form an effector complex, which guides crRNA for invader nucleic acid recognition and destruction in the interference stage (Barrangou et al., 2007; Plagens et al., 2014; Xue and Sashital, 2019). The transcription of these genes suggested the interference activities of the CRISPR-Cas system.

Overall, all necessary genes in the CRISPR-Cas systems were transcribed in both the aerobic and anaerobic stages, suggesting an active CRISPR-Cas system in *Ca.* Accumulibacter SCUT2 in the EBPR cycle (Fig. 2B).

### 3.5. Phylogenetic analysis of the CRISPR-Cas system

Phylogenetic analysis was performed based on the species tree of *Ca.* Accumulibacter (Fig. 1B). The distribution of CRISPR-Cas systems was not in line with the vertical evolutionary relationships among different *Ca.* Accumulibacter members. CRISPR loci from different subtypes scattered across the species tree (Fig. 1B). Even for closely related species, for instance, Clade IIC members SSA1 (obtained from an EBPR reactor in Singapore, Arumugam et al., 2019), SK-02 (obtained in an SBR in Queensland (Skennerton et al., 2015)) and UW6 (from an EBPR reactor in the University of Wisconsin, McDaniel et al., 2021) are closely related (ANI>99.1%) (Fig.1A). However, the occurrence and identity of the CRISPR-Cas systems in these strains showed significant differences. SSA1 harbors a type I-C CRISPR-Cas locus, and UW6 contains type I-A and type I-E CRISPR-Cas loci. No CRISPR-Cas system was found for SK-02. Similar cases were found for Clade IIF members SSB1 and SCELSE-1, clade IIA members UW1 and AALB, and Clade IA members BA93, UBA6658 and UW8. Overall, different members of the same clades were shown to possess different type CRISPR-Cas systems (Fig. 1B).

To further understand the origin and distribution of CRISPR-Cas systems in *Ca.* Accumulibacter, phylogenetic analysis of the marker genes (*cas1* gene for the type I CRISPR-Cas system and *cas10* gene for the type III system) was performed. *cas1* and *cas10* genes in *Ca.* Accumulibacter genomes were classified into 5 and 6 clusters, respectively (Fig. 3A and 3B). Even for the same CRISPR-Cas subtype, the phylogenetic relationship of the *cas* genes was distant. For example, *cas1* genes in type I-E systems were classified into two clusters (Fig. 3A), suggesting that the type I-E CRISPR-Cas system in *Ca.* Accumulibacter came from different sources (e.g., those in UW1 and UW6 are non-homologous, although both were recovered in UW). Similar results were found for subtype I-C (in AALB and SSA1), III-A (in SCELSE-10IIF, AALB and UW7) and III-B (in UW1 and UW7) systems. Overall, the distribution of CRISPR-Cas systems in the *Ca.* Accumulibacter lineage and the *cas* gene phylogenetic analyses strongly suggests that CRISPR-Cas systems in *Ca.* Accumulibacter genomes were not inherited from a common ancestor but obtained via horizontal gene transfer (HGT) from multiple sources. However, *cas1* genes in subtype I-A (e.g., in IIC members UW6, Bin19 and SCELSE-2) and in subtype I-F are highly conserved (e.g., in IIC members SCUT-2 and SCELSE-8). *Ca.* Accumulibacter genomes harboring the same subtype CRISPR-Cas systems (subtype I-A or I-F) are also closely related (Fig.1A and 1B), implying that type I-A and type I-F CRISPR-Cas systems were probably obtained before speciation of these clade members.

**Fig. 3.**
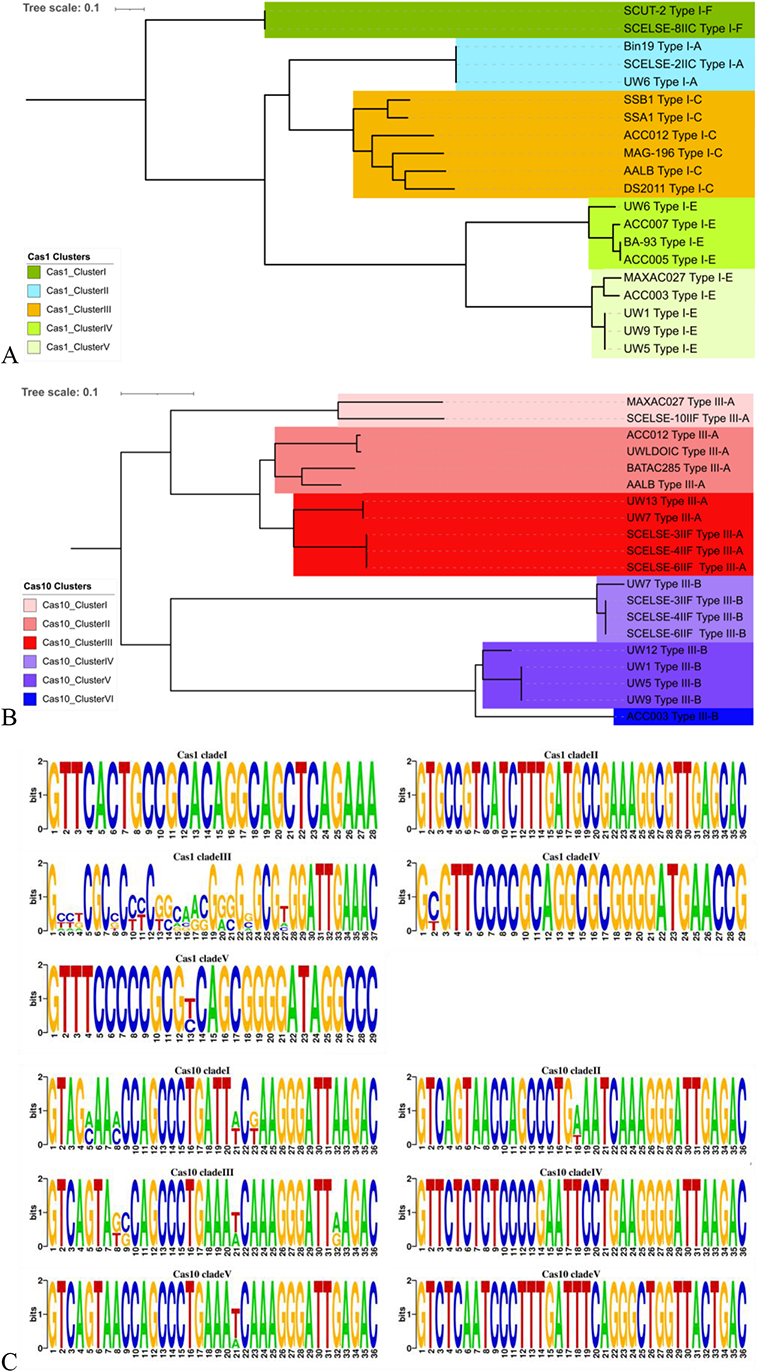
(A) A gene tree of type I CRISPR-Cas systems in different *Ca.* Accumulibacter genomes. *cas1* gene were employed as a marker gene. The tree was grouped into five clusters denoted as I to V. (B) A gene tree of type III CRISPR-Cas systems in different *Ca.* Accumulibacter genomes. *cas10* gene were employed as a marker gene. The tree was grouped into six clusters denoted as I to VI. (C) Multiple sequence alignment and visualization of DRs in *Ca.* Accumulibacter by Weblogo (Crooks et al., 2004). The size of the base letter indicates the frequency of occurrence of the base at a given position.

Apart from *cas* genes, DRs serve as intervals between adjacent spacers, were also gene markers for the CRISPR-Cas system (Chakraborty et al., 2010). Multiple sequence alignment of DRs in each *cas* gene cluster were performed by using WebLogo (Fig. 3C) (Crooks et al., 2004). Results showed that in each *cas* gene cluster, DRs are relatively conservative (e.g., in *cas1*-cluster IV, the sequence identity between each two DRs was > 90%) (Fig. 3B and Fig. S2). DRs from different *cas* gene clusters are largely different (e.g., the sequence identity between DRs in *cas1*-cluster IV and those in *cas1*-cluster V < 75%) (Fig. 3B and Fig. S2). These results indicate a strong relationship between *cas* genes (*cas1* and *cas10*) and DRs. Previous research suggested that *cas* genes and DRs usually move together between different bacteria via HGT (Delaney et al., 2012; Makarova et al., 2015). This seems to be true for the different types of CRISPR-Cas systems in *Ca.* Accumulibacter as showed in this study.

Bringing together the species phylogenetic tree, *cas* gene tree and DRs analysis, all results indicated that CRISPR-Cas systems in different *Ca.* Accumulibacter members was not inherited from their last common ancestors, but were probably obtained at different nodes by different lineage members via HGT, suggesting that phage predation was a very strong selection pressure in the evolution of the *Ca.* Accumulibacter lineage members. The distribution of the CRISPR-Cas systems among different *Ca.* Accumulibacter showed that they were independently acquired for at least 11 times. The major differences in their CRISPR-Cas systems gave rise to distinct abilities of different *Ca.* Accumulibacter to resist predations from different phages. Previous study suggested high frequency of HGT in *Ca*. Accumulibacter by analyzing the genes in 10 *Ca.* Accumulibacter genomes via a pan-genome analysis (Oyserman et al., 2016). HGT of CRISPR-Cas loci was also found to be common in the kingdom of bacteria (Chakraborty et al., 2010; Godde and Bickerton, 2006; Horvath et al., 2009; Westra et al., 2012). Wastewater treatment system is a highly dynamic environment with high occurrence of phages (10^8^-10^9^ ml^-1^) in the activated sludge (Chen et al., 2021). Arm races between phages and their host *Ca.* Accumulibacter are inevitable. CRISPR-Cas systems may have rendered *Ca.* Accumulibacter increased abilities to cope with diverse phage invasions, conferring their broader ecological niche and high occurrence in EBPR systems. The occurrence and distribution of CRISPR-Cas systems is not closely related to the clade classification of *Ca.* Accumulibacter, suggesting that the rate of dynamics of the CRISPR-Cas system in the *Ca*. Accumulibacter lineage was higher than those of speciation.

DRs were compared to the CRISPRCasdb database to tentatively explore the source of CRISPR-Cas systems via HGT. 6 out of 40 DRs matched completely to the CRISPRCasdb database (Table S4). A high number of non-matches may be explained by the fact that a majority of prokaryotes remain un-sequenced (98.9% globally, (Zhang et al., 2020)). All matched DRs had multiple hits. For example, The DRs in Type I-F CRISPR-Cas system of the clade IIC member SCELSE-8, are identical to these in *Desulfurivibrio alkaliphilus* AHT 2(GCA_000092205.1), *Halomonas sp.* Y2R2 (GCA_008329985.2), *Pseudoalteromonas tunicata* D2 (GCA_002310815.1) and *Shewanella inventionis* D1489 (GCA_019931735.1), indicating the possible origination of each CRISPR-Cas system in representative *Ca.* Accumulibacter genomes.

### 3.6. *cas* gene characteristics in *Ca.* Accumulibacter

Through CRISPRCasFinder identify and annotation via RAST, *cas* gene arrangements were analyzed (Brettin et al., 2015). Similar *cas* gene arrangements, sizes, and directions was observed in the same CRISPR-Cas subtypes showed in the different *Ca.* Accumulibacter genomes (Fig. 4).

**Fig. 4.**
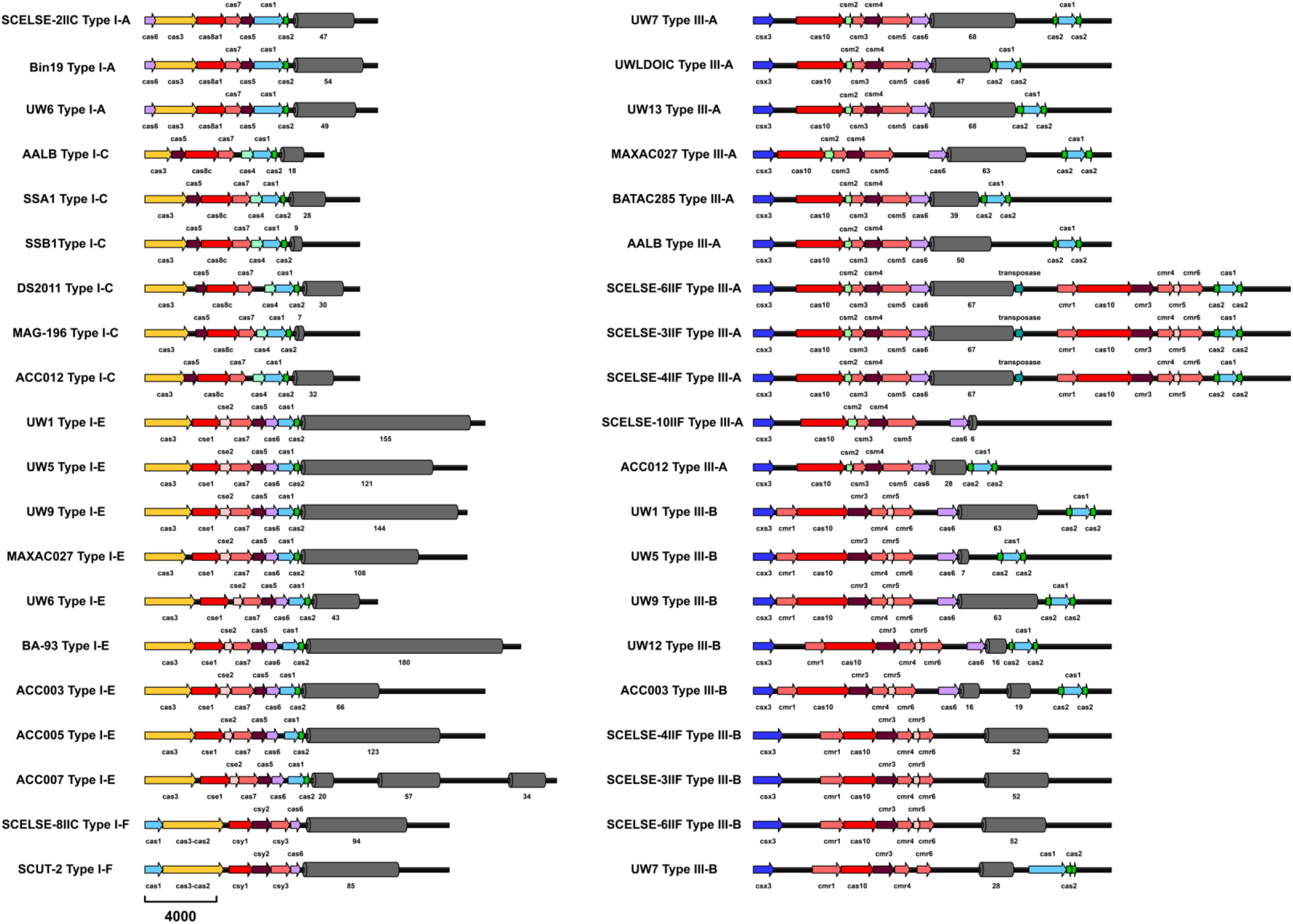
Schematics of CRISPR-Cas systems in *Ca*. Accumulibacter genomes. Arrows in the same color represent homologous *cas* genes. The length of the *cas* gene is indicated by the length of the arrow. Gray cylinders represent the CRISPR array. The numbers below gray cylinders represent the numbers of spacers. Dark green arrows in CRISPR loci SCELSE-6IIF Type III-A, SCELSE-3IIF Type III-A, and SCELSE-4IIF Type III-A indicate the transposase.

*cas* gene arrangements of type I-C, I-E and I-F CRISPR-Cas systems are identical to these in Makarova’s latest classification (Makarova et al., 2020), preserving all necessary immune-related components. *cas* gene arrangements of type I-A, III-A and III-B systems are not consistent with the above-mentioned classification (Fig. 4). For subtype I-A systems, all CRISPR loci (i.e., in clade IIC members SCELSE-2, UW6, and Bin19) lack *cas4* and *cas11*, *cas4* gene encode an endonuclease, which is necessary for efficient pre-spacer processing by forming a Cas4-Cas1-Cas2 complex (Lee et al., 2019). The lack of *cas4* gene may affect an effective acquisition of functional spacers by these clade IIC members. The protein encoded by *cas11* does not directly interact with crRNA, but it is considered as a bridge to connect the head and tail of the CRISPR-Cas effector complexes, which can increase the stability of combination between crRNA and target phage DNA (Xue and Sashital, 2019; Zheng et al., 2020). The lack of *cas11* may affect the interference stage of immunity.

For subtype III-A, all CRISPR-Cas loci lack *csm6* genes. A previous study showed that Csm6 is related to degradation of phage transcripts (Jiang et al., 2016). For subtype III-B, 4 CRISPR-Cas loci (in clade IIF members SCELSE-3, SCELSE-4, SCELSE-6 and UW7) lack *cas6* genes. Cas6 is not an effector complex, but an enzyme responsible for pre-CRISPRRNA (pre-crRNA) processing into mature crRNAs in the interference stage (Makarova et al., 2020). The lack of *cas6* gene may lead to increased inability of CRISPR-Cas systems in these Clade IIF *Ca.* Accumulibacter members to generate crRNAs for invader phage sequences identification.

An interesting observation was that in some type III-A CRISPR loci (in clade IIF members SCELSE-3, SCELSE-4, and SCELSE-6), the transposases genes were found at the downstream of the spacers (Fig. 4) and the *cas* genes located at the downstream of the transposase genes. Transposase are proteins mediating the transposition process (Vigil-Stenman et al., 2017). For prokaryotes, transposons can jump between chromosomal DNA and plasmid, increasing the rate of HGT (Hidalgo-Cantabrana et al., 2017). The presence of transposase in type III-A CRISPR-Cas system suggested that the *cas* genes at the downstream was probably obtained via HGT. The acquisition of these genetic architectures is an evolutionary advantage in a complex environment like in the activated sludge.

### 3.7. Spacers in different *Ca.* Accumulibacter

Spacers record the history of phage infections and show the kind of phages which could be resisted by the host bacteria. Upon phage invasion, spacers are responsible for invader recognition and Cas protein guidance to cleave phage genetic materials. A detailed intercomparison of spacers in different CRISPR-Cas loci was performed (Fig. 5). Major differences were observed in the number of spacers for different CRISPR-Cas systems in different *Ca.* Accumulibacter members (Fig. 5A). Subtype I-C systems were found to contain the lowest number of spacers, ranging from 7 to 32. Subtype I-E systems had the greatest number of spacers, ranging from 43 to 180. The lowest number (6) of spacers was found in *Ca.* Accumulibacter SCELSE-10IIF (Subtype III-A) (recovered from NTU35), suggesting small numbers of phages to which this *Ca.* Accumulibacter is immune. *Ca.* Accumulibacter BA-93 (Subtype I-E) encoded the largest number (180) of spacers (recovered from an SBR with the seed sludge from Thorneside WWTP, Australia, (Skennerton et al., 2015)), implying its potential in readily resisting diverse phages. Although maintaining high numbers of spacers may also be costly, a previous study suggested that the optimal number of spacers is 10 to 100 (Bradde et al., 2020). Most CRISPR-Cas systems in *Ca.* Accumulibacter fall within this range.

**Fig. 5.**
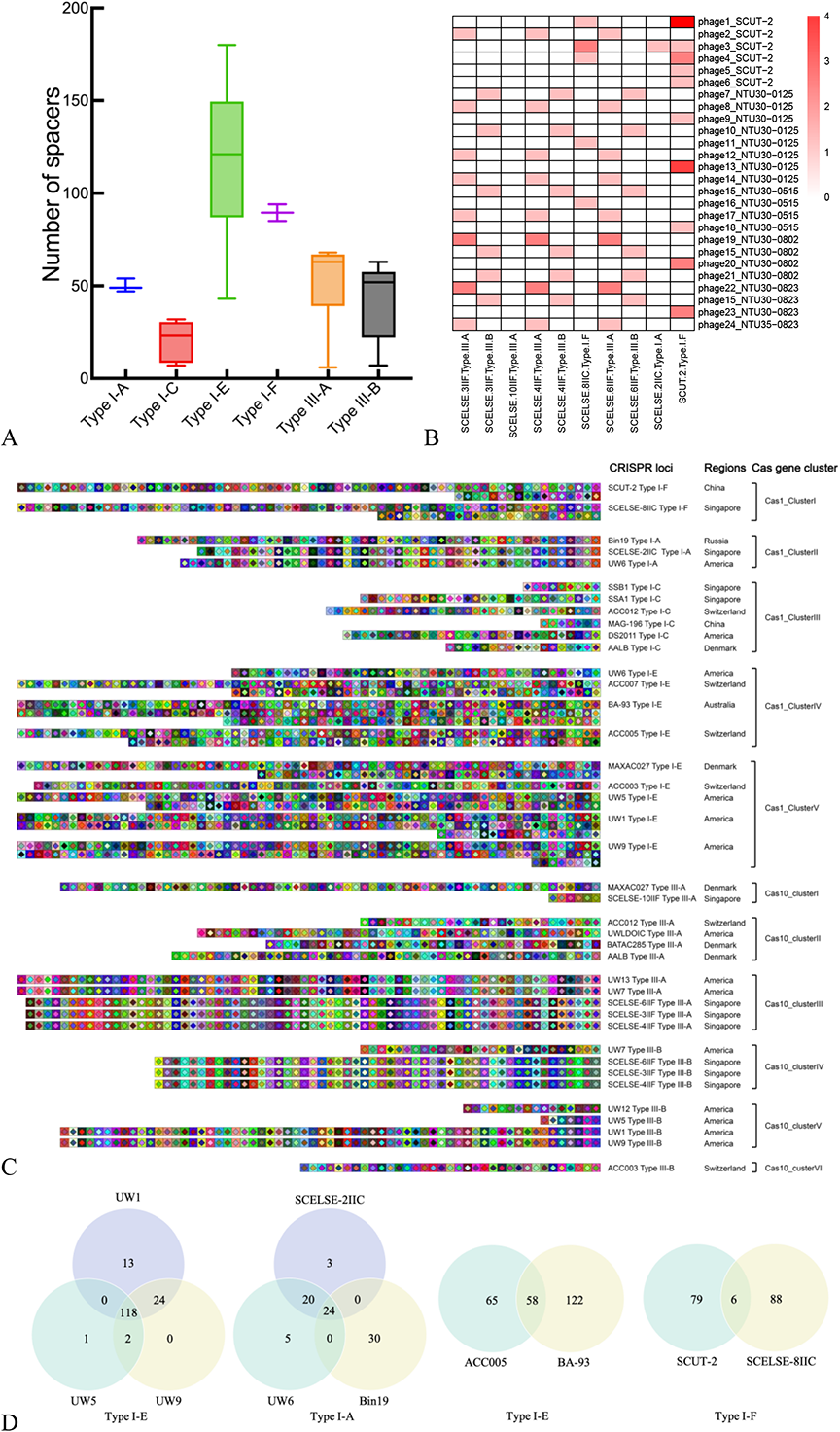
Comparisons of spacers in different CRISPR loci. (A) A boxplot of the number of spacers in different types of CRISPR-Cas systems. (B) Heatmap of spacers in CRISPR loci and phages identified by VirSorter (Guo et al., 2021). The color scale indicates the number of spacers in a CRISPR locus which matched to a specific phage genome. (Phage15_NTU30-0515, phage15_NTU30-0802 and phage15_NTU30-0823 are identical phage contigs retrieved from different metagenomic data). (C) A schematic representation of the CRISPR spacer array in different *Ca.* Accumulibacter. Each square represents a spacer, common spacers are in the same color. (D) Venn diagrams of spacers in different CRISPR loci. The numbers represent the numbers of spacers. The overlapping parts represent the shared numbers of spacers.

Intercomparison of spacers among CRISPR-Cas systems in *Ca.* Accumulibacter was done using CRISPRStudio (Fig. 5C) (Dion et al., 2018). Within each CRISPR-Cas cluster, for genomes having the same *cas* gene (sequence identity=100%), the numbers and composition of the spacers in these CRISPR-Cas loci were analyzed (i.e., Type III-A and Type III-B, respectively, in SCELSE-4IIF, SCELSE-3IIF and SCELSE-6IIF; Type I-E in UW1, UW5 and UW9; Type I-F in SCUT-2 and SCELSE-8IIC; Type I-E in ACC005 and BA-93; and Type I-A in UW6, Bin19 and SCELSE-2IIC). The same *cas* genes indicated the homology of respective CRISPR-Cas systems (obtained from the same ancestor or horizontal transferred from homologic sources). Analysis of spacers in the CRISPR-Cas systems with the same *cas* gene allowed the elucidation of spacer acquisition and deletion in different *Ca.* Accumulibacter.

For Type III-A and Type III-B systems in SCELSE-3IIF, SCELSE-4IIF and SCELSE-6IIF (ANI between each two genomes >99.5%, Fig.1A) no difference was observed in the kind and number of spacers (Fig. 5C), suggesting that no spacer acquisition or deletion occurred during the relatively short period of reactor operation (79 and 21 days apart).

For *Ca*. Accumulibacter members having the same marker *cas* genes, and whose genomes were obtained from the same geographic location, i.e., UW1, UW5 and UW9 (all of them have a Type I-E CRISPR locus), 118 spacers are shared among three genomes. UW1 and UW9 share an additional 24 spacers, and UW5 and UW9 share two more spacers. UW1 and UW5 have 13 and one specific spacers, respectively (Fig. 5C and Fig. 5D). These three *Ca.* Accumulibacter were obtained from enrichment cultures inoculated with seed sludge from the same WWTP (Nine Springs, Madison, USA), but were recovered approximately 11 and 12LJyears apart (McDaniel et al., 2021). ANI analysis showed that UW1, UW5 and UW9 were closely related (>99.5%, Fig. 1A). The differences in their spacers may be a result of spacer acquisition and deletion over this time period. When comparing the CRISPR loci in UW1 and UW5, three spacers were newly acquired, and 37 spacers were lost. Previous research also observed spacer acquisition and deletion in *Streptococcus* in laboratory experiments and natural conditions (Deveau et al., 2008; Lopez-Sanchez et al., 2012). This phenomenon may be explained by the fact that bacteria tend to acquire new spacers and delete less valuable spacers (since phages tend to escape immunity through point mutations) to adapt to changing environments (Horvath et al., 2008). A decrease in the number of spacers is not necessarily unfavorable. As suggested by Martynov et al. (Martynov et al., 2017), the acquisition and loss rate of spacers determines the steady state number of spacers, which changes along with the environment. The changes in the number of spacers in closely related species in the same geographic location suggested that the CRISPR-Cas systems in these *Ca.* Accumulibacter members are active and evolving.

For genomes with the same *cas* gene but obtained from different geographic locations, for instance, both clade IIC members SCUT-2 (retrieved from our lab-scale reactor in the South China University of Technology, China) and SCELSE-8 (retrieved from our lab-scale reactor in the Nanyang Technological University, Singapore) have a Type I-F CRISPR-Cas locus. SCUT-2 and SCELSE-8IIC harbor 85 and 94 spacers in their CRISPR-Cas loci, respectively (Fig. 5C and Fig. 5D). Only six spacers were shared between these two genomes. Similarly, both clade IA members ACC005 and BA-93 have a Type I-E CRISPR-Cas locus. ACC005 was retrieved from a SBR with seed sludge from the ARA Thunersee WWTP, Switzerland (Adler and Holliger, 2020). BA-93 was retrieved from a SBR with the seed sludge from Thorneside WWTP, Australia, (Skennerton et al., 2015). Fifty-eight spacers are shared between these two genomes, and 122 and 65 spacers are specific for BA-93 and ACC005, respectively (Fig. 5C and Fig. 5D). Once again, clade IIC members UW6 (retrieved from an enrichment culture inoculated with seed sludge from the Nine Springs WWTP, Madison, WI, USA (McDaniel et al., 2021), Bin19 (assembled from a SBR in the Russian Academy of Sciences (Kotlyarov et al., 2019) and SCELSE-2IIC (obtained in this study from NTU30), all have a Type I-A CRISPR-Cas system. Twenty-four identical spacers were shared among three genomes. UW6 and SCELSE-2IIC shared additional 20 identical spacers. UW6, SCELSE-2IIC and Bin19 had 5, 3 and 30 specific spacers, respectively (Fig. 5C and 5D). For genomes having the same *cas* marker gene but from different regions, their CRISPR-Cas systems may have been obtained before they were dispersed to different geographic locations. The shared spacers reflected the history of phage infections before these *Ca.* Accumulibacter were geographically separated. The specific spacers indicated that they had experienced distinct phage predation events after having been exposed to different locations and environment.

Spacers record the infection history of different *Ca.* Accumulibacter members by different phages. By using spacers as a marker, the identity of phages which infected different *Ca.* Accumulibacter members may be revealed (Díez-Villaseñor and Rodriguez-Valera, 2019). The CRISPR-Cas system can only provide resistance if a specific spacer identically matches the genomic sequence of a phage (Deveau et al., 2008). Thus, only spacers completely mapped to phage sequences were considered in this study. In total, 2448 spacers were identified in all CRISPR-Cas systems, 67 of them completely matched to the IMG/VR database, of which 52 are classified as Caudovirales, 3 as Tubulavirales, and 12 as unclassified phages. Among these phages, 63 were discovered from lab-scale EBPR bioreactors or full-scale WWTPs, and four were obtained from natural water bodies (Table S3). For example, a spacer (GCTGGCGTAACCGGCCTTGCGCCTGGAAAGGA) in clade IIA members UW1, UW5 and UW9 completely matched a phage affiliated to a Siphoviridae family member under the order of Caudovirales, which occurred in an EBPR reactor in the University of Queensland, Australia. Further, a spacer (GCAGGGCAGCAGCGGGCCGGGCGACTTGAACT) in UW7 completely matched a *Rauchvirus* genus phage member (also belonging to the order Caudovirales), which occurred in the same EBPR reactor. Collectively, these results suggested that Caudovirales are a key group of known phages that infect *Ca.* Accumulibacter.

To further analyze the relationship between phages and spacers in *Ca*. Accumulibacter, viral data mining was performed in the metagenomic data obtained from our lab-scale SBRs. In total, 24193 (out of 618997) contigs were identified as viral contigs by VirSorter (Guo et al., 2021), accounting for 3.59-4.34% of total contigs in 6 metagenomic datasets (Fig. S4). A viral contig dataset was constructed, and the spacers in *Ca.* Accumulibacter genomes (retrieved from these lab-scale reactor) were compared to the viral genome dataset. Within 10 CRISPR-Cas loci in 7 *Ca.* Accumulibacter genomes retrieved from our lab-scale SBR, 41 (out of 589) spacers matched to 26 viral contigs (including three identical viral contigs retrieved from three different metagenomic datasets) (Fig. 5B). For instance, 14 spacers in SCUT-2 match 10 viral contigs in the viral datasets. The matches between the spacers in the CRISPR-Cas loci and the occurring phages in the SBR suggest that the CRISPR-Cas loci were likely active in resisting these phages in the reactors. We also found that multiple spacers can match to viral contigs in different regions (Fig. 5B and Table S5). For example, in SCUT-2, four spacers matched to the same viral contig (Caudovirales IMGVR_UVIG3300013800_000258), and two spacers matched to another viral contig (Caudovirales IMGVR_UVIG3300028804_000035). This phenomenon suggests that these phages are under extremely stringent selection pressure and are unlike to evade the *Ca.* Accumulibacter SCUT-2’s CRISPR immune system via single-nucleotide point mutations (Nasko et al., 2019). The results also suggest that *Ca.* Accumulibacter SCUT-2 and these two phages may have experienced long-term co-evolution.

Furthermore, the viral contigs which matched to spacers were compared to the IMG/VR database. 18 of them were identified as Caudovirales order members (including 1 Podoviridae family members, 2 Siphoviridae family members, Table S6). These results again suggested that Caudovirales is a key group of phages targeting *Ca.* Accumulibacter.

### 3.8. Prophages in *Ca*. Accumulibacter genomes

Prophages in the *Ca.* Accumulibacter genome represent a threat to the cell and may be activated with or without external stimuli (Cu^2+^, KCN or ciprofloxacin) (Motlagh et al., 2015). All 43 *Ca*. Accumulibacter genomes were found to contain prophages; a total of 133 prophage-like regions were identified (Fig. 6A and Table S7). Each *Ca.* Accumulibacter genome contained at least one prophage-like region, for example, *Ca.* Accumulibacter SCELSE-8IIC (retrieved from our lab-scale reactor NTU30) harbors 5 prophages, and *Ca.* Accumulibacter clade IIA members AALB (obtained from a enrichment reactor in Aalborg, Denmark (Albertsen et al., 2016) and UW1 (obtained from a enrichment reactor in UW) both harbor two prophage regions. Clade IIC member SSA1 and clade IIF member SSB1 both have 5 prophages. The size of prophage-like regions accounted for 0.15% (MAG-196) to 2.97% (ACC005, Table S7) of *Ca*. Accumulibacter genomes, at an average of 1.06%. 6.77% (9) and 13.53% (18) of the prophage-like regions were classified as potentially inducible (intact and questionable) prophages, respectively, ranging from 6.8kb to 49.2kb (26.3kb in average). The rest are incomplete prophages, ranging from 4.8kb to 35.3kb (13.04kb in average). 19 genomes were found to have intact or questionable prophages (e.g., including UW13, SSA1, SK-02 and SCELSE-5IIH, Fig. 6A and Table S7). No correlation was observed between the size of the *Ca.* Accumulibacter genome and the accumulative size of prophage-like regions (R^2^=0.2, Figure S5). In bacterial genomes, the frequent occurrence of incomplete prophages has been reported, owing to pervasive domestication of defective prophages (Bobay et al., 2014). According to genomic streamlining theory (Lawrence et al., 2001), incomplete prophages typically occur with neutral or beneficial genes being retained and the genes which are essential for prophage activation being removed. The high numbers of putatively defective prophage sequences observed in these *Ca.* Accumulibacter genomes indicate rapid inactivation of prophages by *Ca.* Accumulibacter together with a slow decay of prophage genes (Brüssow et al., 2004). Incomplete prophages typically lack the ability to entry the lytic cycle and kill the host. Prophages in *Ca.* Accumulibacter genomes may also benefit cells, e.g., via the superinfection exclusion mechanism, where prophages tend to prevent subsequent phage infections of their host (Bondy-Denomy and Davidson, 2014), like those in *Escherichia coli* and *Salmonella* spp. (Labrie et al., 2010). The capability of *Ca.* Accumulibacter to resist lytic phages may also be enhanced due to the expression of genes that increase fitness of the host cell by a process known as lysogenic conversion. However, since previous research showed that prophages in *Ca.* Accumulibacter could be induced with external stimuli (Motlagh et al., 2015), the presence of intact and questionable prophages in *Ca.* Accumulibacter represents a threat to them and hence EBPR stability.

**Fig. 6.**
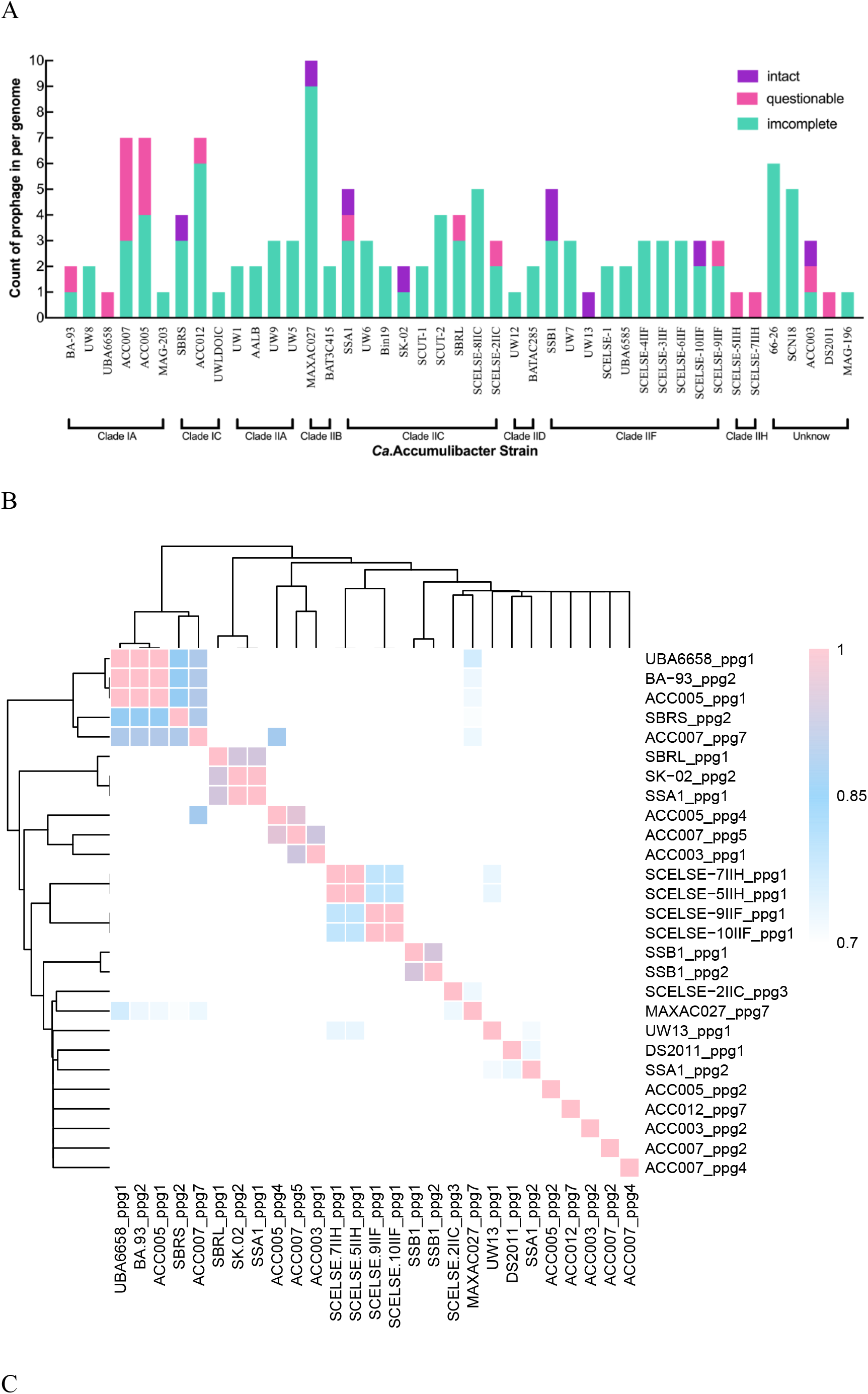
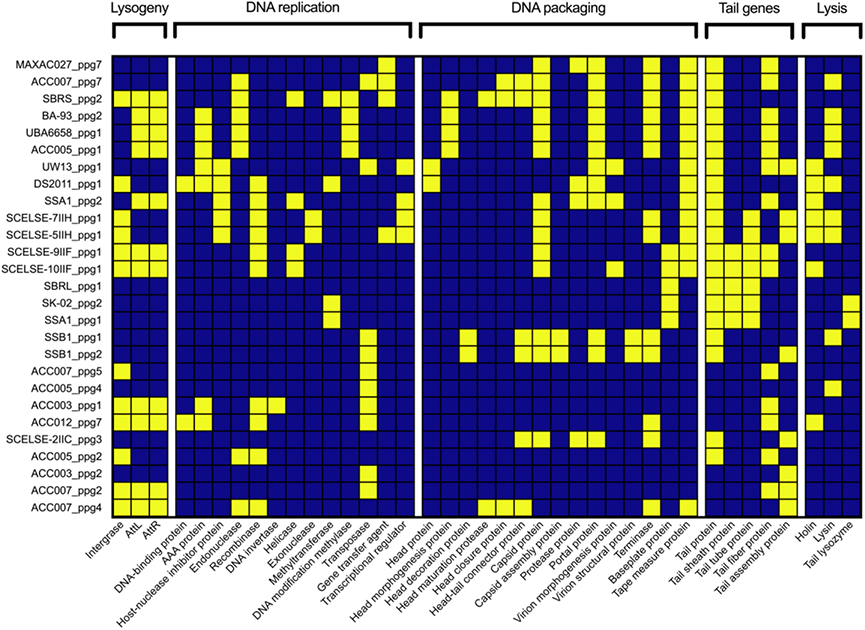
(A) Occurrence and distribution of prophage-like elements in 43 *Ca.* Accumulibacter genomes. (B) Average nucleotide identity (ANI) analysis of different prophages in *Ca.* Accumulibacter genomes. (C) Prophage gene identification based on prophage gene modules. The prophage genes were divided into five functional modules. The name of each functional module was given on the top of the heatmap. Yellow square represents the occurrence of a respective gene.

ANI analysis showed that prophages in *Ca.* Accumulibacter are highly diverse (ANI<70%, Fig. 6B). There are also closely related prophages (e.g., BA-93_ppg2, UBA6658_ppg1 and ACC005_ppg1, ANI=100%) whose host *Ca.* Accumulibacter genomes also shared close phylogenetic relationships (ANI > 98.9%, Fig. 6B), implying that these prophages might have been inherited from their latest ancestor.

### 3.9. Prophage gene integrity

For intact and questionable prophages, their genes were classified into five modules as lysogeny, DNA replication, DNA packaging, tail morphogenesis, and host lysis (Fig. 6B). The tail morphogenesis module was well-preserved in intact and questionable prophages (26 out of 27 intact and questionable prophages have the tail gene module). On the contrary, the genes classified into the lysis module showed the lowest occurrence. It was identified only in 15 prophages, three of which harbored both the lysin and holin genes. Holins and lysins are viral enzymes that lead to host cell destruction for phage lease (Wang et al., 2000). The presence of these genes confers higher lysogeny (Feiner et al., 2015). Three of these prophages (i.e., SCELSE-5IIH_ppg1, SCELSE-7IIH_ppg1, and DS2011_ppg1, encoding both holin and lysin genes) are readily inducible. The induction of prophages may lead to the extinction of host *Ca*. Accumulibacter cells and adversely affect the EBPR performance. Prophages which lack of holin and/or lysin genes are not necessarily harmless. For example, the *Dickeya* spp. specific phage vB_DsoM_LIMEstone1 and the *Pectobacterium carotovorum* specific phage PP1 lack the genes encoding holin and lysin, respectively, but both were shown to be highly infectious (Adriaenssens et al., 2012; Lee et al., 2012). It is possible that some hypothetic protein genes may have encoded the holin and lysin.

All intact and questionable prophages were compared against the IMG/VR database. Except for 8 prophages, which were unclassified, all prophages were identified as Caudovirales order members, (including 6 Siphoviridae family members, Table S8). These results are consistent with the previous finding that the induced prophages in *Ca.* Accumulibacter in a full-scale WWTP were classified as Caudovirales members (Motlagh et al., 2015), and confirms that Caudovirales is a major group of phages interacting with *Ca.* Accumulibacter.

Among all genomes, three (MAXAC027, BATAC285, and BAT3C415) were retrieved from full-scale WWTPs in Danmark (Singleton et al., 2021). Two of them had CRISPR-Cas systems. MAXAC027 harbored Type I-E (108 spacers) and III-A (63 spacers) CRISPR-Cas systems together with 10 prophages in its genome (1 intact and 9 questionable). BATAC285 encoded a Type III-A (39 spacers) CRISPR-Cas system and 2 incomplete prophages. The overall occurrence of the CRISPR-Cas system in these genomes (2/3) was similar to that in all analyzed genomes (65.1%). Comparing to other *Ca.* Accumulibacter, numbers of spacers and prophages in MAXAC027 were in a high rang (87.4 spacers and 3.16 prophage in average), suggesting that MAXAC027 had experienced extensive phage selection, which may endorse it an elevated level of robustness upon phage predation.

## 4. CONCLUSIONS

- Highly diverse CRISPR-Cas systems present in different *Ca.* Accumulibacter, classifying into 6 subtypes, including a subtype I-F CRISPR-Cas system, which was only found in the newly recovered *Ca.* Accumulibacter genomes (two clade IIC members SCELSE-8 and SCUT-2).
- Polygenetic analysis of *Ca.* Accumulibacter species tree and the *cas* gene, and multiple sequence alignments of DRs, suggest that the CRISPR-Cas systems in *Ca.* Accumulibacter are not commonly inherited from their ancestors but obtained at different nodes by different lineage members via HGT, implying high phage selection pressure during the evolution and speciation of the *Ca*. Accumulibacter lineage members.
- The diverse and distinct CRISPR-Cas systems impart specific immunity to phage predation to different *Ca.* Accumulibacter.
- In long-term bioreactor operation, *Ca.* Accumulibacter taxa with CRISPR-Cas systems exhibited high relative abundance, indicating that *Ca.* Accumulibacter with a CRISPR-Cas system are more likely to survive and prosper under phage-mediated selection. Metatranscriptomic analysis showed transcription of CRISPR-associated genes in both aerobic and anaerobic stages, demonstrating active CRISPR-Cas systems in *Ca.* Accumulibacter throughout the SBR cycle.
- Different *Ca.* Accumulibacter species have distinct phage predation histories, showing great spatial heterogeneity in closely related *Ca.* Accumulibacter species from different geographic locations and long-term temporal dynamics in those from the same geographic location. Caudovirales is a key group of phages threatening *Ca.* Accumulibacter.
- 133 prophage-like regions were identified. All *Ca.* Accumulibacter harbored prophages. 15 prophages were highly threatening to their host *Ca*. Accumulibacter strains. Three of them (in SCELSE-5IIH, SCELSE-7IIH and DS2011) are readily inducible.

## Acknowledgements

This research was partially supported by the National Natural Science Foundation of China (51808297), the Natural Science Foundation of Guangdong Province (2021A1515010494), the Guangzhou Science and Technology Planning Program (202002030340), the Pearl River Talent Recruitment Program (2019QN01L125), and the Program for Science and Technology of Guangdong Province, China (No. 2018A050506009). Additional support came from the Singapore National Research Foundation and the Ministry of Education under the Research Centre of Excellence Programme, and through a research grant from the National Research Foundation under its Environment and Water Industry Programme (project number 1102–IRIS– 10–02), administered by PUB-Singapore’s national water agency.

## Data available

All data generated or analyzed during this study are included in this published article. (Metagenomic raw reads and draft genomes were submitted to NCBI under the BioProject No. PRJNA807832 and No. PRJNA771771. Metatranscriptomic data were submitted to NCBI under the submitted No. PRJNA807832. Other data were showed in the *Supplementary Materials*.)

## CRediT authorship contribution statement

**Xuhan Deng:** Conceptualization, Methodology, Software, Formal analysis, Investigation, Data Curation, Writing – Original Draft, Visualization.

**Jing Yuan:** Data Curation, Resources, Visualization.

**Hang Chen:** Investigation, Resources, Data Curation.

**Liping Chen:** Investigation, Resources, Data Curation.

**Chaohai Wei:** Writing – review & editing, Supervision.

**Per H. Nielsen:** Writing – review & editing, Supervision.

**Stefan Wuertz:** Conceptualization, Supervision, Writing – review & editing, Project administration, Funding acquisition.

**Guanglei Qiu:** Conceptualization, Methodology, Investigation, Supervision, Writing – review & editing, Validation, Project administration, Funding acquisition.

