## Supplementary file for "CRISPR-Cas phage defense systems and prophages in *Candidatus* Accumulibacter"

### S1 Sequencing batch reactor (SBRs) operation

***S1.1 Operation of the SBRs NTU30 and NTU35***

Two SBRs (operated in parallel at 30 ℃ (NTU30) and 35 ℃ (NTU35)) with a working volume of 1.59 L each were inoculated with activated sludge from a wastewater treatment plant (WWTP) in Singapore. The SBRs were operated with 6 h cycles, including a 60 min feeding, a 20 min anaerobic, a 180 min aerobic, and a 100 min settling/decant stage. In each cycle, 0.74 L of synthetic wastewater (containing 193.8 mg/L acetate, 17.2 mg/L propionate and 8.8 mg/L PO4^3-^-P) was introduced into the reactor, with a resultant TOC/P molar ratio of 25:1. Temperature control was achieved using a proportional-integral-derivative temperature controller connected to heating jackets wrapping the reactors. The HRT and SRT in both reactors were 12.9 h and 25 d, respectively. The pH was automatically controlled at 6.80–7.50 by using a M200 transmitter connected to an acid/base (0.5 M HCl/NaOH) dosing system. The DO was maintained at 0.8–1.2 mg/L during the aerobic phase by using the same transmitter connected to a solenoid valve in the aeration system. During Days 175–220, sodium acetate and propionic acid concentrations in the feed for NTU35 were increased 1.6 times (resultant TOC/P molar ratio of 40:1), which was reduced to 1.2 times (TOC/P molar ratio of 30:1) from Day 220 onwards until the end of the experiment.

***S1.2 Operation of the SBR SCUT***

The SBR SCUT with a working volume of 4.5 L was inoculated with activated sludge collected from a WWTP in Guangzhou, China. The SBR was operated with 6 h cycles, including a slow-feeding phase (60 min), an anaerobic phase (20 min), an aerobic phase (180 min), and a settling/decant phase (100 min). Acetate was used as sole carbon source for the reactor. In each cycle, 2.5 L of synthetic wastewater (containing 100 mg/L acetate, and 20 mg/L PO_4_^3-^-P) was fed into the reactor, with a resultant TOC/P molar ratio of 15:1. The HRT and SRT in the reactor were 12 h and 15 d, respectively. The pH was automatically controlled at 7.0-7.5 by using a M200 transmitter (Mettler-Toledo, Switzerland) connected to an acid/base (0.5 M HCl/NaOH) dosing system. The DO was maintained at 0.8-1.2 mg/L during the aerobic phase by using the same transmitter connected to a solenoid valve in the aeration system. Temperature was controlled at 25℃, using a temperature controller connected to thermostatic water bath.

***S1.3 DNA extraction and metagenomic analysis***

For SBRs NTU30 and NTU35, activated sludge samples were collected on Day 14, Day 56, Day 91, Day201, Day 280 and Day 301 from each SBR. Genomic DNA were extracted using the Fast DNATM 2 mL SPIN Kit for Soil samples (MP Biomedicals, CA, USA) following the manufacturer’s instructions, and stored at -80 °C prior to metagenomic analysis. Sequencing library preparation was performed using a modified version of the Illumina TruSeq DNA Sample Preparation protocol: 1 μg DNA was sheared on a Covaris S220 to approximately 300 bp, following the manufacturer’s recommendation. Size selection was performed on a Sage Science Pippin Prep instrument, using a 2% EtBr agarose cassette and selecting for a tight peak around 400 bp. Each library was tagged with a TruSeq LT DNA barcode (Illumina, CA, USA) to allow for library pooling prior to sequencing. Library quantitation was performed using the Picogreen assay (Invitrogen, CA, USA) and the average library size was determined by running the libraries on a Bioanalyzer DNA 7500 chip (Agilent, CA, USA). Library concentrations were normalized to 4 nM and validated by qPCR on a ViiA-7 realtime thermocycler (Applied Biosystems, CA, US), using qPCR primers recommended by Illumina in their qPCR protocol, and the Illumina PhiX control library was used as a standard. Libraries were then combined in one pool, which was sequenced across two lanes of an Illumina HiSeq2500 sequencing run at a read-length of 250 bp paired-end. Raw reads have been submitted to NCBI and are accessible under the BioProject No. PRJNA807832.

For SBR SCUT, activated sludge was collected on Day 783. Genomic DNA were extracted using the OMEGA Soil DNA Kit (D5625-01) (Omega Bio-Tek, GA, USA), following the manufacturer’s instructions, and stored at -80 °C prior to further analysis. The quantity and quality of the extracted DNA were measured using a Qubit™ 4 Fluorometer (with WiFi: Q33238; Qubit™ Assay Tubes: Q32856; Qubit™ 1X dsDNA HS Assay Kit: Q33231) and 1% agarose gel electrophoresis, respectively. Genomic library was constructed following the Illumina TruSeq DNA Sample Preparation Guide. Illumina NovaSeq sequencing run at a read-length of 150 bp paired-end. Raw reads have been submitted to NCBI and are accessible under the BioProject No. PRJNA771771.

**S2 Metatranscriptomic analysis**

To further confirm the activity of CRISPR-Cas systems, *Ca.* Accumulibacter-PAO-enriched activated sludge (MLSS= 3.8 g/L; relative abundance of 37.09 %) from the SCUT SBR (on Day 783) was collected for metagenomic analysis. Activated sludge samples were collected just before the start of the SBR cycle (0 min), at 5 min (anaerobic phase), 30 min (anaerobic phase), 105min (anaerobic phase), 120 min (aerobic phase), snap-frozen in liquid N_2_, and stored at -80 ^o^C before RNA extraction for metatranscriptomic analysis.

Total RNA was extracted using the RNA PowerSoil® Total RNA Isolation Kit (Omega Bio-Tek, GA, USA). The quality and quantity of the extracted RNA were measured using 1.5% agarose gel electrophoresis and UV spectrophotometer, respectively. cDNA library was constructed using a TruSeq Standard mRNA LT Sample Prep Kit (Illumina, CA, USA). The quality and quantity of the library were measured using Agilent Bioanalyzer and Promega QuantiFluor, respectively. Illumina NovaSeq sequencing run at a read-length of 150 bp paired-end. Raw reads have been submitted to NCBI and are accessible under the BioProject No. PRJNA771771.

**Table S1.** SBRs recovered *Ca*. Accumulibacter genomes information and genomes evaluation results

| **Number** | **Strain** | **Species name** | **Clades** | **GenBank No.** | **Date of sampling** | **Source** | **Relative abundance (%)** | **Completeness (%)** | **Contamination (%)** |
| --- | --- | --- | --- | --- | --- | --- | --- | --- | --- |
| 1 | SCELSE-2 | *Ca.* Accumulibacter cognatus | IIC | GCA_023806705.1 | 2018.1.25 | NTU30 | 5.02 | 95.4 | 1.11 |
| 2 | SCELSE-3 | *--* | IIF | GCA_023806635.1 | 2018.5.15 | NTU30 | 12.38 | 98.57 | 0.27 |
| 3 | SCELSE-4 | *--* | IIF | GCA_023806545.1 | 2018.8.2 | NTU30 | 19.82 | 96.22 | 0.0 |
| 4 | SCELSE-5 | -- | IIH | GCA_023806585.1 | 2018.8.2 | NTU30 | 6.9 | 97.2 | 0.61 |
| 5 | SCELSE-6 | *--* | IIF | GCA_023806505.1 | 2018.8.23 | NTU30 | 18.28 | 96.22 | 0.48 |
| 6 | SCELSE-7 | *--* | IIH | GCA_023806565.1 | 2018.8.23 | NTU30 | 7.85 | 98.1 | 0.61 |
| 7 | SCELSE-8 | *Ca.* Accumulibacter cognatus | IIC | GCA_023806525.1 | 2018.8.23 | NTU30 | 3.03 | 95.47 | 2.06 |
| 8 | SCELSE-9 | *--* | IIF | GCA_023806595.1 | 2018.1.25 | NTU35 | 12.28 | 97.83 | 0.19 |
| 9 | SCELSE-10 | -- | IIF | GCA_023806625.1 | 2018.8.23 | NTU35 | 1.27 | 98.78 | 0.21 |
| 10 | SCUT-2 | *Ca.* Accumulibacter cognatus | IIC | GCA_024380085.1 | 2021.11.17 | SCUT | 37.09 | 98.1 | 1.11 |

**Table S2.** GenBank *Ca.* Accumulibacter genomes information and genomes evaluation results

| **Number** | **Genomes** | **Code** | **Species name** | **Clades** | **GenBank assembly accession** | **Completeness** | **Contaminess** | **Citation** |
| --- | --- | --- | --- | --- | --- | --- | --- | --- |
| 1 | BA-93 | BA-93 | *Ca*. Accumulibacter regalis | IA | GCA_000585075.1 | 100 | 0.27 | A lab-scale SBR seeded with activated sludge from Thorneside Wastewater Treatment Plant, Queensland, Australia (Skennerton et al., 2015) |
| 2 | UW1 | UW1 | *Ca*. Accumulibacter phosphatis | IIA | GCA_000024165.1 | 100 | 0.24 | A lab-scale SBR inoculated with activated sludge from the Madison, WI, USA Nine Springs Wastewater Treatment Plant (Martín et al., 2006) |
| 3 | UW8-POB | UW8 | *Ca*. Accumulibacter regalis | IA | GCA_017302345.1 | 99.84 | 1.69 | A lab-scale SBR seeded with activated sludge from wastewater treatment systems (McDaniel et al., 2021b) |
| 4 | 66-26 | 66-26 | -- | unknown | GCA_001897745.1 | 99.76 | 0.03 | A lab-scale reactor inoculated with activated sludge from a demonstration-scale bioreactor treating SCN-containing mining effluent(Kantor et al., 2015) |
| 5 | SCN18_14_9_16_R1_B_66_13 | SCN18 | *--* | unknown | GCA_017307795.1 | 99.76 | 1.5 | Biofilm fraction from a lab-scale bioreactor (Huddy et al., 2021) |
| 6 | aalborg | AALB | *Ca*. Accumulibacter aalborgensis | IIA | GCA_900089955.1 | 99.52 | 0.06 | A lab-scale SBR seeded with activated sludge from the full-scale EBPR plant operating in Aalborg West, Denmark (Albertsen et al., 2016) |
| 7 | UW9-POB | UW9 | *Ca*. Accumulibacter phosphatis | IIA | GCA_017302455.1 | 99.37 | 4.05 | A lab-scale SBR seeded with activated sludge from wastewater treatment systems (McDaniel et al., 2021b) |
| 8 | UBA6658 | UBA6658 | *--* | IA | GCA_002455435.1 | 99.08 | 0.24 | A lab-scale EBPR bioreactor sample (Parks et al., 2017) |
| 9 | SSA1 | SSA1 | *Ca.* Accumulibacter cognatus | IIC | GCA_013414765.1 | 99.05 | 1.11 | A lab-scale SBR inoculated with activated sludge from an EBPR mother reactor (Arumugam et al., 2019) |
| 10 | SSB1 | SSB1 | *Ca*. Accumulibacter similis | IIF | GCA_013347225.1 | 99.05 | 0.03 | A lab-scale SBR inoculated with activated sludge from an EBPR mother reactor (Arumugam et al., 2019) |
| 11 | UW5 | UW5 | *Ca*. Accumulibacter phosphatis | IIA | GCA_017592745.1 | 98.99 | 5.24 | A lab-scale SBR seeded with activated sludge from the Nine Springs Wastewater Treatment Plant (McDaniel et al., 2021a) |
| 12 | UW7 | UW7 | *Ca*. Accumulibacter conexus | IIF | GCA_017592775.1 | 98.97 | 3.04 | A lab-scale SBR seeded with activated sludge from the Nine Springs Wastewater Treatment Plant (McDaniel et al., 2021a) |
| 13 | UW6 | UW6 | *Ca*. Accumulibacter cognatus | IIC | GCA_017592725.1 | 98.57 | 2.46 | A lab-scale SBR seeded with activated sludge from the Nine Springs Wastewater Treatment Plant (McDaniel et al., 2021a) |
| 14 | ACC003_d322 | ACC003 | *--* | unknown | GCA_907163085.1 | 98.57 | 0.35 | A lab-scale aerobic granular sludge (AGS) reactor (Adler et al., 2022) |
| 15 | Bin19 | Bin19 | *Ca.* Accumulibacter cognatus | IIC | GCA_005889575.1 | 98.51 | 1.94 | A lab-scale SBR reactor with seed activated sludge obtained from a pilot wastewater treatment system (Kotlyarov et al., 2019) |
| 16 | DS2.011 | DS2011 | *--* | unknown | GCA_013823155.1 | 98.37 | 0 | A tap of groundwater-sourced drinking water system (Zhang et al., 2017) |
| 17 | UW12-POB | UW12 | *Ca.* Accumulibacter necessarius | IID | GCA_017302435.1 | 98.1 | 0.98 | A lab-scale SBR seeded with activated sludge from wastewater treatment systems (McDaniel et al., 2021b) |
| 18 | ACC007_d740 | ACC007 | *--* | IA | GCA_907163205.1 | 98.1 | 4.39 | A lab-scale aerobic granular sludge (AGS) reactor (Adler et al., 2022) |
| 19 | ACC005_d427 | ACC005 | *--* | IA | GCA_907163215.1 | 97.85 | 2.17 | A lab-scale aerobic granular sludge (AGS) reactor (Adler et al., 2022) |
| 20 | Fred_18-Q3-R57-64_MAXAC.027 | MAXAC027 | *Ca*. Accumulibacter propinquus | IIB | GCA_016712935.1 | 97.67 | 4 | Fresh activated sludge samples collected from Danish WWTPs in August and September 2018 (Singleton et al., 2021) |
| 21 | SK-02 | SK-02 | *Ca.* Accumulibacter cognatus | IIC | GCA_000584975.2 | 97.62 | 1.59 | A lab-scale SBR seeded with activated sludge from Thorneside Wastewater Treatment Plant, Queensland, Australia (Skennerton et al., 2015) |
| 22 | SBR_S | SBRS | *Ca*. Accumulibacter delftensis | IC | GCA_012939955.1 | 97.34 | 0.66 | Unknow |
| 23 | UW13-POB | UW13 | *Ca*. Accumulibacter conexus | IIF | GCA_017302415.1 | 97.14 | 4.39 | A lab-scale SBR seeded with activated sludge from wastewater treatment systems (McDaniel et al., 2021b) |
| 24 | SCUT-1 | SCUT-1 | *--* | IIC | GCA_020709745.1 | 96.98 | 1.59 | A lab-scale SBR inoculated with activated sludge collected from a local WWTP in Guangzhou, China (Tian et al., 2022) |
| 25 | SBR_L | SBRL | *Ca.* Accumulibacter contiguus | IIC | GCA_012940005.1 | 96.85 | 0.98 | Unknow |
| 26 | EsbW_18-Q3-R4-48_BATAC.285 | BATAC285 | *Ca.* Accumulibacter proximus | IID | GCA_016709675.1 | 96.67 | 3.63 | Fresh activated sludge samples collected from Danish WWTPs in August and September 2018 (Singleton et al., 2021) |
| 27 | UBA6585 | UBA6585 | *--* | IIF | GCA_002433845.1 | 96.51 | 0.15 | A lab-scale EBPR bioreactor sample (Parks et al., 2017) |
| 28 | ACC012_d740 | ACC012 | *--* | IC | GCA_907163155.1 | 95.87 | 4.63 | A lab-scale aerobic granular sludge (AGS) reactor (Adler et al., 2022) |
| 29 | new MAG-196 | MAG-196 | *--* | unknown | GCA_016789495.1 | 95.52 | 0.48 | A lab-scale SBR seeded with activated sludge obtained from a full-scale WWTP in Danyang, China (Lin et al., 2021) |
| 30 | new MAG-203 | MAG-203 | *--* | IA | GCA_016790605.1 | 95.27 | 1.01 | A lab-scale SBR seeded with activated sludge obtained from a full-scale WWTP in Danyang, China (Lin et al., 2021) |
| 31 | OdNE_18-Q3-R46-58_BAT3C.415 | BAT3C415 | *Ca.* Accumulibacter propinquus | IIB | GCA_016714935.1 | 95.26 | 0.98 | Fresh activated sludge samples collected from Danish WWTPs in August and September 2018 (Singleton et al., 2021) |
| 32 | UWLDOIC | UWLDOIC | *Ca*. Accumulibacter meliphilus | IC | GCA_003332265.1 | 95.24 | 1.1 | A lab-scale SBR was used in this study. The reactor was originally inoculated with activated sludge obtained from the Nine Springs wastewater treatment plant in Madison (Camejo et al., 2019) |
| 33 | SCELSE-1 | SCELES-1 | *Ca.* Accumulibacter similis | IIF | GCA_005524045.1 | 95.24 | 0.6 | A lab-scale SBR inoculated with activated sludge from a full-scale WWTP in Singapore (Qiu et al., 2020) |

**Table S3.** CRISPR loci in *Ca.* Accumulibacter and spacers comparison against the IMG/VR database.

| **No** | **Strain** | **CRISPR Loci** | **CRRISPR-Cas Subtype** | **Total Spacers** | **Matched spacers** | **Spacer Squence** | **IMG/VR_UViGs** | **Lineage** | **Genome Name** | **E-value** | **Identities** |
| --- | --- | --- | --- | --- | --- | --- | --- | --- | --- | --- | --- |
| 1 | Bin19 | Bin19 Type I-A | I-A | 54 | 0 |  |  |  |  |  |  |
| 2 | UW6 | UW6 Type I-A | I-A | 49 | 0 |  |  |  |  |  |  |
| 3 | SCELSE-2IIC | SCELSE-2IIC Type I-A | I-A | 47 | 1 | TGGTAGTCGATCACCCAGGTCTTGCCGCCTTTGCCG | [IMGVR_UViG_3300028804_000035](https://img.jgi.doe.gov/cgi-bin/vr/main.cgi?section=ViralBrowse&page=uviginfo&uvig_id=IMGVR_UViG_3300028804_000035) | Duplodnaviria; Heunggongvirae; Uroviricota; Caudoviricetes; Caudovirales; ; ; ; | Activated sludge microbial communities from WWTP in Nijmegen, Gelderland, Netherland - WWTP Weurt | 2.00E-09 | 36/36 100% |
| 4 | SSA1 | SSA1 Type I-C | I-C | 28 | 3 | ACGATTGAGGAGATTTGCGCCATGCCGGTAGCT | [IMGVR_UViG_3300000730_000048](https://img.jgi.doe.gov/cgi-bin/vr/main.cgi?section=ViralBrowse&page=uviginfo&uvig_id=IMGVR_UViG_3300000730_000048) | Duplodnaviria; Heunggongvirae; Uroviricota; Caudoviricetes; Caudovirales; ; ; ; | Activated sludge microbial communities from San Jose/Santa Clara Water Pollution Control Plant, California, USA | 6.00E-08 | 33/33 100% |
|  |  |  | I-C |  |  | GGCATTGTCGATGCGGCTTTCGGCAGCGGCGCGAA | [IMGVR_UViG_3300006092_000164](https://img.jgi.doe.gov/cgi-bin/vr/main.cgi?section=ViralBrowse&page=uviginfo&uvig_id=IMGVR_UViG_3300006092_000164) | Duplodnaviria; Heunggongvirae; Uroviricota; Caudoviricetes; Caudovirales; ; ; ; | Activated sludge microbial communities from wastewater treatment plant in Ulu Pandan, Singapore | 5.00E-09 | 35/35 100% |
|  |  |  | I-C |  |  | GTTGTTATCGGACGATTTGTTGCTGCGCCCTTTCAG | [IMGVR_UViG_3300028804_000035](https://img.jgi.doe.gov/cgi-bin/vr/main.cgi?section=ViralBrowse&page=uviginfo&uvig_id=IMGVR_UViG_3300028804_000035) | Duplodnaviria; Heunggongvirae; Uroviricota; Caudoviricetes; Caudovirales; ; ; ; | Activated sludge microbial communities from WWTP in Nijmegen, Gelderland, Netherland - WWTP Weurt | 2.00E-09 | 36/36 100% |
| 5 | DS2011 | DS2011 Type I-C | I-C | 30 | 0 |  |  |  |  |  |  |
| 6 | SSB1 | SSB1 Type I-C | I-C | 9 | 1 | ACTTCAATCTTGGAATTGAATCGGAACGGCAAAT | [IMGVR_UViG_3300005676_000005](https://img.jgi.doe.gov/cgi-bin/vr/main.cgi?section=ViralBrowse&page=uviginfo&uvig_id=IMGVR_UViG_3300005676_000005) | Duplodnaviria; Heunggongvirae; Uroviricota; Caudoviricetes; Caudovirales; ; ; ; | Enhanced biological phosphorus removal bioreactor viral communities from the University of Queensland, Australia - SBR4-V90903 Phage Sequencing | 2.00E-08 | 34/34 100% |
| 7 | AALB | AALB Type I-C | I-C | 18 | 0 |  |  |  |  |  |  |
| 8 | ACC012 | ACC012 Type I-C | I-C | 32 | 0 |  |  |  |  |  |  |
| 9 | MAG-196 | MAG-196 Type I-C | I-C | 7 | 0 |  |  |  |  |  |  |
| 10 | UW6 | UW6 Type I-E | I-E | 49 | 1 | GTTCAAACTGACATGGAAGCAGCAGGCTACGC | [IMGVR_UViG_3300035661_000141](https://img.jgi.doe.gov/cgi-bin/vr/main.cgi?section=ViralBrowse&page=uviginfo&uvig_id=IMGVR_UViG_3300035661_000141) | Duplodnaviria; Heunggongvirae; Uroviricota; Caudoviricetes; Caudovirales; Podoviridae; ; Rauchvirus; | Filamentous streamer microbial communities from Yellowstone Lake, YNP, Wyoming, USA - RO1DNA_8/2/18_2/3_streamer | 2.00E-07 | 32/32 100% |
| 11 | UW9 | UW9 Type I-E | I-E | 144 | 4 | CCATCGCTCCAGGCGACACGCCGCAAGCGGCG | [IMGVR_UViG_3300025896_000038](https://img.jgi.doe.gov/cgi-bin/vr/main.cgi?section=ViralBrowse&page=uviginfo&uvig_id=IMGVR_UViG_3300025896_000038) | Duplodnaviria; Heunggongvirae; Uroviricota; Caudoviricetes; Caudovirales; ; ; ; | Aqueous microbial communities from the Delaware River and Bay under freshwater to marine salinity gradient to study organic matter cycling in a time-series - DEBay_Spr_0.19_<0.8_DNA (SPAdes) | 2.00E-07 | 32/32 100% |
|  |  |  |  |  |  | AACGGGGACCGGGCTGGCTACGCGTCAAGCTG | [IMGVR_UViG_3300005479_000008](https://img.jgi.doe.gov/cgi-bin/vr/main.cgi?section=ViralBrowse&page=uviginfo&uvig_id=IMGVR_UViG_3300005479_000008) | Monodnaviria; Loebvirae; Hofneiviricota; Faserviricetes; Tubulavirales; Inoviridae; ; ; | Activated sludge microbial communities from EBPR reactors in the US and Australia - sample from Queensland, Australia | 2.00E-07 | 32/32 100% |
|  |  |  |  |  |  | GCTGGCGTAACCGGCCTTGCGCCTGGAAAGGA | [IMGVR_UViG_3300005678_000068](https://img.jgi.doe.gov/cgi-bin/vr/main.cgi?section=ViralBrowse&page=uviginfo&uvig_id=IMGVR_UViG_3300005678_000068) | Duplodnaviria; Heunggongvirae; Uroviricota; Caudoviricetes; Caudovirales; Siphoviridae; ; ; | Enhanced biological phosphorus removal bioreactor viral communities from the University of Queensland, Australia - SBR4-V91307 Phage Sequencing | 2.00E-07 | 32/32 100% |
|  |  |  |  |  |  | GAACGCATCCGGCGCGCGCACTACGTGCGAGA | [IMGVR_UViG_3300025877_000066](https://img.jgi.doe.gov/cgi-bin/vr/main.cgi?section=ViralBrowse&page=uviginfo&uvig_id=IMGVR_UViG_3300025877_000066) | Duplodnaviria; Heunggongvirae; Uroviricota; Caudoviricetes; Caudovirales; ; ; ; | Active sludge microbial communities of municipal wastewater-treating anaerobic digesters from USA - AD_STIC10_MetaG (SPAdes) | 2.00E-07 | 32/32 100% |
| 12 | UW5 | UW5 Type I-E | I-E | 121 | 4 | CCATCGCTCCAGGCGACACGCCGCAAGCGGCG | [IMGVR_UViG_3300025896_000038](https://img.jgi.doe.gov/cgi-bin/vr/main.cgi?section=ViralBrowse&page=uviginfo&uvig_id=IMGVR_UViG_3300025896_000038) | Duplodnaviria; Heunggongvirae; Uroviricota; Caudoviricetes; Caudovirales; ; ; ; | Aqueous microbial communities from the Delaware River and Bay under freshwater to marine salinity gradient to study organic matter cycling in a time-series - DEBay_Spr_0.19_<0.8_DNA (SPAdes) | 2.00E-07 | 32/32 100% |
|  |  |  |  |  |  | GGCGGACGCGAAGCGCTGCACCACTATTTGGT | [IMGVR_UViG_3300025877_000066](https://img.jgi.doe.gov/cgi-bin/vr/main.cgi?section=ViralBrowse&page=uviginfo&uvig_id=IMGVR_UViG_3300025877_000066) | Duplodnaviria; Heunggongvirae; Uroviricota; Caudoviricetes; Caudovirales; ; ; ; | Active sludge microbial communities of municipal wastewater-treating anaerobic digesters from USA - AD_STIC10_MetaG (SPAdes) | 2.00E-07 | 32/32 100% |
|  |  |  |  |  |  | GCTGGCGTAACCGGCCTTGCGCCTGGAAAGGA | [IMGVR_UViG_3300005678_000068](https://img.jgi.doe.gov/cgi-bin/vr/main.cgi?section=ViralBrowse&page=uviginfo&uvig_id=IMGVR_UViG_3300005678_000068) | Duplodnaviria; Heunggongvirae; Uroviricota; Caudoviricetes; Caudovirales; Siphoviridae; ; ; | Enhanced biological phosphorus removal bioreactor viral communities from the University of Queensland, Australia - SBR4-V91307 Phage Sequencing | 2.00E-07 | 32/32 100% |
|  |  |  |  |  |  | AACGGGGACCGGGCTGGCTACGCGTCAAGCTG | [IMGVR_UViG_3300005479_000008](https://img.jgi.doe.gov/cgi-bin/vr/main.cgi?section=ViralBrowse&page=uviginfo&uvig_id=IMGVR_UViG_3300005479_000008) | Monodnaviria; Loebvirae; Hofneiviricota; Faserviricetes; Tubulavirales; Inoviridae; ; ; | Activated sludge microbial communities from EBPR reactors in the US and Australia - sample from Queensland, Australia | 2.00E-07 | 32/32 100% |
| 13 | MAXAC027 | MAXAC027 Type I-E | I-E | 108 | 0 |  |  |  |  |  |  |
| 14 | ACC003 | ACC003 Type I-E | I-E | 66 | 1 | GCAATGTCAAGCGCTGCGTGTGCGTCGAGCAG | [IMGVR_UViG_3300027781_000088](https://img.jgi.doe.gov/cgi-bin/vr/main.cgi?section=ViralBrowse&page=uviginfo&uvig_id=IMGVR_UViG_3300027781_000088) | Duplodnaviria; Heunggongvirae; Uroviricota; Caudoviricetes; Caudovirales; ; ; ; | Wastewater effluent complex algal communities from Wisconsin, to seasonally profile nutrient transformation and Carbon sequestration - JI 9/18/14 C2 DNA (SPAdes) | 2.00E-07 | 32/32 100% |
| 15 | ACC005 | ACC005 Type I-E | I-E | 123 | 7 | GCCCGTCAGTGATGTACGACTACTGGATGACC | [IMGVR_UViG_3300005625_000006](https://img.jgi.doe.gov/cgi-bin/vr/main.cgi?section=ViralBrowse&page=uviginfo&uvig_id=IMGVR_UViG_3300005625_000006) | Duplodnaviria; Heunggongvirae; Uroviricota; Caudoviricetes; Caudovirales; ; ; ; | Activated sludge viral communities from EBPR reactors in Madison, Wisconsin, USA v1 | 2.00E-07 | 32/32 100% |
|  |  |  |  |  |  | CGGCATTCAGGAGCGCGAGCAAAAGCGCCAGG | [IMGVR_UViG_3300005625_000006](https://img.jgi.doe.gov/cgi-bin/vr/main.cgi?section=ViralBrowse&page=uviginfo&uvig_id=IMGVR_UViG_3300005625_000006) | Duplodnaviria; Heunggongvirae; Uroviricota; Caudoviricetes; Caudovirales; ; ; ; | Activated sludge viral communities from EBPR reactors in Madison, Wisconsin, USA v1 | 2.00E-07 | 32/32 100% |
|  |  |  |  |  |  | CGAACAGGAGGAAATGCACCGCATGTTGACGG | [IMGVR_UViG_3300003782_000016](https://img.jgi.doe.gov/cgi-bin/vr/main.cgi?section=ViralBrowse&page=uviginfo&uvig_id=IMGVR_UViG_3300003782_000016) | Duplodnaviria; Heunggongvirae; Uroviricota; Caudoviricetes; Caudovirales; ; ; ; | Wastewater treatment Type I Accumulibacter community from EBPR Bioreactor in Madison, WI, USA - Reactor 1_4/24/2008_ DNA | 2.00E-07 | 32/32 100% |
|  |  |  |  |  |  | CAGCGCAGCAAGGGCGGGCGGAGAATGAAGCC | [IMGVR_UViG_3300005686_000119](https://img.jgi.doe.gov/cgi-bin/vr/main.cgi?section=ViralBrowse&page=uviginfo&uvig_id=IMGVR_UViG_3300005686_000119) | Duplodnaviria; Heunggongvirae; Uroviricota; Caudoviricetes; Caudovirales; ; ; ; | Enhanced biological phosphorus removal bioreactor viral communities from the University of Queensland, Australia - SBR4-V92206 Phage Sequencing | 2.00E-07 | 32/32 100% |
|  |  |  |  |  |  | CCAGCATCTCTGCTGGCTCTCTTTGGGCGCAC | [IMGVR_UViG_3300029936_000040](https://img.jgi.doe.gov/cgi-bin/vr/main.cgi?section=ViralBrowse&page=uviginfo&uvig_id=IMGVR_UViG_3300029936_000040) | Duplodnaviria; Heunggongvirae; Uroviricota; Caudoviricetes; Caudovirales; ; ; ; | Activated sludge bacterial and viral communities from EBPR bioreactors in Queensland, Australia - SBR4-V92206 | 2.00E-07 | 32/32 100% |
|  |  |  |  |  |  | ACGATTTCCAGAAGCTCGCCGGCAATCACTGG | [IMGVR_UViG_3300005625_000006](https://img.jgi.doe.gov/cgi-bin/vr/main.cgi?section=ViralBrowse&page=uviginfo&uvig_id=IMGVR_UViG_3300005625_000006) | Duplodnaviria; Heunggongvirae; Uroviricota; Caudoviricetes; Caudovirales; ; ; ; | Activated sludge viral communities from EBPR reactors in Madison, Wisconsin, USA v1 | 2.00E-07 | 32/32 100% |
|  |  |  |  |  |  | ACCCGAACTTGCTTACCGCTTCTGCAATGATC | [IMGVR_UViG_3300026284_000009](https://img.jgi.doe.gov/cgi-bin/vr/main.cgi?section=ViralBrowse&page=uviginfo&uvig_id=IMGVR_UViG_3300026284_000009) |  | Wastewater treatment Type I Accumulibacter community from EBPR Bioreactor in Madison, WI, USA - Reactor 1_6/14/2005_ DNA (SPAdes) | 2.00E-07 | 32/32 100% |
| 16 | ACC007 | ACC007 Type I-E | I-E | 111 | 1 | GTCTGCAGACGTCGCGCAGAGATTGGTGCCGC | [IMGVR_UViG_3300026299_000007](https://img.jgi.doe.gov/cgi-bin/vr/main.cgi?section=ViralBrowse&page=uviginfo&uvig_id=IMGVR_UViG_3300026299_000007) | Duplodnaviria; Heunggongvirae; Uroviricota; Caudoviricetes; Caudovirales; ; ; ; | Wastewater treatment Type I Accumulibacter community from EBPR Bioreactor in Madison, WI, USA - Reactor 1_9/17/2007_ DNA (SPAdes) | 2.00E-07 | 32/32 100% |
| 17 | UW1 | UW1 Type I-E | I-E | 155 | 4 | CCATCGCTCCAGGCGACACGCCGCAAGCGGCG | [IMGVR_UViG_3300025896_000038](https://img.jgi.doe.gov/cgi-bin/vr/main.cgi?section=ViralBrowse&page=uviginfo&uvig_id=IMGVR_UViG_3300025896_000038) | Duplodnaviria; Heunggongvirae; Uroviricota; Caudoviricetes; Caudovirales; ; ; ; | Aqueous microbial communities from the Delaware River and Bay under freshwater to marine salinity gradient to study organic matter cycling in a time-series - DEBay_Spr_0.19_<0.8_DNA (SPAdes) | 2.00E-07 | 32/32 100% |
|  |  |  |  |  |  | AACGGGGACCGGGCTGGCTACGCGTCAAGCTG | [IMGVR_UViG_3300005479_000008](https://img.jgi.doe.gov/cgi-bin/vr/main.cgi?section=ViralBrowse&page=uviginfo&uvig_id=IMGVR_UViG_3300005479_000008) | Monodnaviria; Loebvirae; Hofneiviricota; Faserviricetes; Tubulavirales; Inoviridae; ; ; | Activated sludge microbial communities from EBPR reactors in the US and Australia - sample from Queensland, Australia | 2.00E-07 | 32/32 100% |
|  |  |  |  |  |  | GCTGGCGTAACCGGCCTTGCGCCTGGAAAGGA | [IMGVR_UViG_3300005678_000068](https://img.jgi.doe.gov/cgi-bin/vr/main.cgi?section=ViralBrowse&page=uviginfo&uvig_id=IMGVR_UViG_3300005678_000068) | Duplodnaviria; Heunggongvirae; Uroviricota; Caudoviricetes; Caudovirales; Siphoviridae; ; ; | Enhanced biological phosphorus removal bioreactor viral communities from the University of Queensland, Australia - SBR4-V91307 Phage Sequencing | 2.00E-07 | 32/32 100% |
|  |  |  |  |  |  | GAACGCATCCGGCGCGCGCACTACGTGCGAGA | [IMGVR_UViG_3300025877_000066](https://img.jgi.doe.gov/cgi-bin/vr/main.cgi?section=ViralBrowse&page=uviginfo&uvig_id=IMGVR_UViG_3300025877_000066) | Duplodnaviria; Heunggongvirae; Uroviricota; Caudoviricetes; Caudovirales; ; ; ; | Active sludge microbial communities of municipal wastewater-treating anaerobic digesters from USA - AD_STIC10_MetaG (SPAdes) | 2.00E-07 | 32/32 100% |
| 18 | BA-93 | BA-93 Type I-E | I-E | 180 | 10 | GCCCGTCAGTGATGTACGACTACTGGATGACC | [IMGVR_UViG_3300005625_000006](https://img.jgi.doe.gov/cgi-bin/vr/main.cgi?section=ViralBrowse&page=uviginfo&uvig_id=IMGVR_UViG_3300005625_000006) | Duplodnaviria; Heunggongvirae; Uroviricota; Caudoviricetes; Caudovirales; ; ; ; | Activated sludge viral communities from EBPR reactors in Madison, Wisconsin, USA | 2.00E-07 | 32/32 100% |
|  |  |  |  |  |  | TACGCGGCCGGGGCGACGGACGCGGAAAAGGC | [IMGVR_UViG_3300005982_000112](https://img.jgi.doe.gov/cgi-bin/vr/main.cgi?section=ViralBrowse&page=uviginfo&uvig_id=IMGVR_UViG_3300005982_000112) | Duplodnaviria; Heunggongvirae; Uroviricota; Caudoviricetes; Caudovirales; ; ; ; | Wastewater effluent complex algal communities from Wisconsin, to seasonally profile nutrient transformation and Carbon sequestration - JI 8/11/14 A brown DNA | 2.00E-07 | 32/32 100% |
|  |  |  |  |  |  | AAATCGCGGAGCGCATCAAGCGCACGATCCCG | [IMGVR_UViG_3300005625_000006](https://img.jgi.doe.gov/cgi-bin/vr/main.cgi?section=ViralBrowse&page=uviginfo&uvig_id=IMGVR_UViG_3300005625_000006) | Duplodnaviria; Heunggongvirae; Uroviricota; Caudoviricetes; Caudovirales; ; ; ; | Activated sludge viral communities from EBPR reactors in Madison, Wisconsin, USA | 2.00E-07 | 32/32 100% |
|  |  |  |  |  |  | CGCACGATCCCGCCCGAATTGCTCGGCGATGA | [IMGVR_UViG_3300005625_000006](https://img.jgi.doe.gov/cgi-bin/vr/main.cgi?section=ViralBrowse&page=uviginfo&uvig_id=IMGVR_UViG_3300005625_000006) | Duplodnaviria; Heunggongvirae; Uroviricota; Caudoviricetes; Caudovirales; ; ; ; | Activated sludge viral communities from EBPR reactors in Madison, Wisconsin, USA | 2.00E-07 | 32/32 100% |
|  |  |  |  |  |  | TCAGTAATCTCCGTATTTCTTCTACGGAAAGT | [IMGVR_UViG_3300029931_000024](https://img.jgi.doe.gov/cgi-bin/vr/main.cgi?section=ViralBrowse&page=uviginfo&uvig_id=IMGVR_UViG_3300029931_000024) |  | Activated sludge bacterial and viral communities from EBPR bioreactors in Queensland, Australia - SBR4-V90308 | 2.00E-07 | 32/32 100% |
|  |  |  |  |  |  | CTGTCGCCGCTGTTCAATATTCCGGCCTCCTGG | [IMGVR_UViG_3300001035_000001](https://img.jgi.doe.gov/cgi-bin/vr/main.cgi?section=ViralBrowse&page=uviginfo&uvig_id=IMGVR_UViG_3300001035_000001) | Duplodnaviria; Heunggongvirae; Uroviricota; Caudoviricetes; Caudovirales; ; ; ; | Wastewater treatment Type I Accumulibacter community from EBPR Bioreactor in Madison, WI - Accumulibacter clade IA genome-pooled fosmids | 6.00E-08 | 33/33 100% |
|  |  |  |  |  |  | CTGCTCGGGCCAGCGCTGGCCGCGACGGGTTC | [IMGVR_UViG_3300005686_000119](https://img.jgi.doe.gov/cgi-bin/vr/main.cgi?section=ViralBrowse&page=uviginfo&uvig_id=IMGVR_UViG_3300005686_000119) | Duplodnaviria; Heunggongvirae; Uroviricota; Caudoviricetes; Caudovirales; ; ; ; | Enhanced biological phosphorus removal bioreactor viral communities from the University of Queensland, Australia - SBR4-V92206 Phage Sequencing | 2.00E-07 | 32/32 100% |
|  |  |  |  |  |  | AGCGAGACATCACCGGCGAGTCGGAGCGCGTT | [IMGVR_UViG_3300029936_000040](https://img.jgi.doe.gov/cgi-bin/vr/main.cgi?section=ViralBrowse&page=uviginfo&uvig_id=IMGVR_UViG_3300029936_000040) | Duplodnaviria; Heunggongvirae; Uroviricota; Caudoviricetes; Caudovirales; ; ; ; | Activated sludge bacterial and viral communities from EBPR bioreactors in Queensland, Australia - SBR4-V92206 | 2.00E-07 | 32/32 100% |
|  |  |  |  |  |  | GTTGTACTGGTCGCGCTGCCCGGATAGCGCGC | [IMGVR_UViG_3300005625_000006](https://img.jgi.doe.gov/cgi-bin/vr/main.cgi?section=ViralBrowse&page=uviginfo&uvig_id=IMGVR_UViG_3300005625_000006) | Duplodnaviria; Heunggongvirae; Uroviricota; Caudoviricetes; Caudovirales; ; ; ; | Activated sludge viral communities from EBPR reactors in Madison, Wisconsin, USA | 2.00E-07 | 32/32 100% |
|  |  |  |  |  |  | AGACGATTGGCTATTCCGAGCTTCTGCCAGAG | [IMGVR_UViG_3300026284_000009](https://img.jgi.doe.gov/cgi-bin/vr/main.cgi?section=ViralBrowse&page=uviginfo&uvig_id=IMGVR_UViG_3300026284_000009) |  | Wastewater treatment Type I Accumulibacter community from EBPR Bioreactor in Madison, WI, USA - Reactor 1_6/14/2005_ DNA (SPAdes) | 2.00E-07 | 32/32 100% |
| 19 | SCELSE-8IIC | SCELSE-8IIC Type I-F | I-F | 94 | 1 | TCTGCAGCAACGACGAAACCCCCTGCGCCCTC | [IMGVR_UViG_3300028804_000035](https://img.jgi.doe.gov/cgi-bin/vr/main.cgi?section=ViralBrowse&page=uviginfo&uvig_id=IMGVR_UViG_3300028804_000035) | Duplodnaviria; Heunggongvirae; Uroviricota; Caudoviricetes; Caudovirales; ; ; ; | Activated sludge microbial communities from WWTP in Nijmegen, Gelderland, Netherland - WWTP Weurt | 2.00E-07 | 32/32 100% |
| 20 | SCUT-2 | SCUT-2 Type I-F | I-F | 85 | 3 | ATCTCGCGCACCCGTGCCCGCTGTTCTTCAAT | [IMGVR_UViG_3300009655_000158](https://img.jgi.doe.gov/cgi-bin/vr/main.cgi?section=ViralBrowse&page=uviginfo&uvig_id=IMGVR_UViG_3300009655_000158) |  | Active sludge microbial communities of municipal wastewater-treating anaerobic digesters from Japan - AD_JPNTR4_MetaG | 2.00E-07 | 32/32 100% |
|  |  |  |  |  |  | TCCTTGTCCCGCAGTCGGTCCCGGTGCAGTTT | [IMGVR_UViG_3300013800_000258](https://img.jgi.doe.gov/cgi-bin/vr/main.cgi?section=ViralBrowse&page=uviginfo&uvig_id=IMGVR_UViG_3300013800_000258) | Duplodnaviria; Heunggongvirae; Uroviricota; Caudoviricetes; Caudovirales; ; ; ; | Wastewater microbial communities from municipal sewage treatment plant in Nanjing, China - ZZ_EW_meta | 2.00E-07 | 32/32 100% |
|  |  |  |  |  |  | GCGGATTCAAGGGCTTCTATCAGTTCGTTCTT | [IMGVR_UViG_3300028804_000035](https://img.jgi.doe.gov/cgi-bin/vr/main.cgi?section=ViralBrowse&page=uviginfo&uvig_id=IMGVR_UViG_3300028804_000035) | Duplodnaviria; Heunggongvirae; Uroviricota; Caudoviricetes; Caudovirales; ; ; ; | Activated sludge microbial communities from WWTP in Nijmegen, Gelderland, Netherland - WWTP Weurt | 2.00E-07 | 32/32 100% |
| 21 | UW7 | UW7 Type III-A | III-A | 68 | 0 |  |  |  |  |  |  |
| 22 | UW13 | UW13 Type III-A | III-A | 68 | 0 |  |  |  |  |  |  |
| 23 | UWLDOIC | UWLDOIC Type III-A | III-A | 47 | 0 |  |  |  |  |  |  |
| 24 | MAXAC027 | MAXAC027 Type III-A | III-A | 63 | 0 |  |  |  |  |  |  |
| 25 | BATAC285 | BATAC285 Type III-A | III-A | 39 | 0 |  |  |  |  |  |  |
| 26 | SCELSE-3IIF | SCELSE-3IIF Type III-A | III-A | 67 | 3 | CCGCCAAGTACGACGGCCTCAACCCCGGCCAGAAG | [IMGVR_UViG_3300028804_000099](https://img.jgi.doe.gov/cgi-bin/vr/main.cgi?section=ViralBrowse&page=uviginfo&uvig_id=IMGVR_UViG_3300028804_000099) |  | Activated sludge microbial communities from WWTP in Nijmegen, Gelderland, Netherland - WWTP Weurt | 5.00E-09 | 35/35 100% |
|  |  |  |  |  |  | CGCGAACTGCAACCGGAGTGGAACAGGGCTTTTGG | [IMGVR_UViG_3300013771_000004](https://img.jgi.doe.gov/cgi-bin/vr/main.cgi?section=ViralBrowse&page=uviginfo&uvig_id=IMGVR_UViG_3300013771_000004) | Duplodnaviria; Heunggongvirae; Uroviricota; Caudoviricetes; Caudovirales; ; ; ; | Activated sludge bacterial and viral communities from EBPR bioreactors in Brisbane, Australia - M92206 | 5.00E-09 | 35/35 100% |
|  |  |  |  |  |  | CCAGATGAGCGTCGCAACCGCGGTGCGCATTTT | [IMGVR_UViG_3300005675_000059](https://img.jgi.doe.gov/cgi-bin/vr/main.cgi?section=ViralBrowse&page=uviginfo&uvig_id=IMGVR_UViG_3300005675_000059) | Duplodnaviria; Heunggongvirae; Uroviricota; Caudoviricetes; Caudovirales; Podoviridae; ; ; | Enhanced biological phosphorus removal bioreactor viral communities from the University of Queensland, Australia - SBR4-V90806 Phage Sequencing | 6.00E-08 | 33/33 100% |
| 27 | SCELSE-4IIF | SCELSE-4IIF Type IIIA | III-A | 67 | 3 | AAAATGCGCACCGCGGTTGCGACGCTCATCTGG | [IMGVR_UViG_3300005675_000059](https://img.jgi.doe.gov/cgi-bin/vr/main.cgi?section=ViralBrowse&page=uviginfo&uvig_id=IMGVR_UViG_3300005675_000059) | Duplodnaviria; Heunggongvirae; Uroviricota; Caudoviricetes; Caudovirales; Podoviridae; ; ; | Enhanced biological phosphorus removal bioreactor viral communities from the University of Queensland, Australia - SBR4-V90806 Phage Sequencing | 6.00E-08 | 33/33 100% |
|  |  |  |  |  |  | CCAAAAGCCCTGTTCCACTCCGGTTGCAGTTCGCG | [IMGVR_UViG_3300013771_000004](https://img.jgi.doe.gov/cgi-bin/vr/main.cgi?section=ViralBrowse&page=uviginfo&uvig_id=IMGVR_UViG_3300013771_000004) | Duplodnaviria; Heunggongvirae; Uroviricota; Caudoviricetes; Caudovirales; ; ; ; | Activated sludge bacterial and viral communities from EBPR bioreactors in Brisbane, Australia - M92206 | 5.00E-09 | 35/35 100% |
|  |  |  |  |  |  | CTTCTGGCCGGGGTTGAGGCCGTCGTACTTGGCGG | [IMGVR_UViG_3300028804_000099](https://img.jgi.doe.gov/cgi-bin/vr/main.cgi?section=ViralBrowse&page=uviginfo&uvig_id=IMGVR_UViG_3300028804_000099) |  | Activated sludge microbial communities from WWTP in Nijmegen, Gelderland, Netherland - WWTP Weurt | 5.00E-09 | 35/35 100% |
| 28 | SCELSE-6IIF | SCELSE-6IIF Type III-A | III-A | 67 | 3 | AAAATGCGCACCGCGGTTGCGACGCTCATCTGG | [IMGVR_UViG_3300005675_000059](https://img.jgi.doe.gov/cgi-bin/vr/main.cgi?section=ViralBrowse&page=uviginfo&uvig_id=IMGVR_UViG_3300005675_000059) | Duplodnaviria; Heunggongvirae; Uroviricota; Caudoviricetes; Caudovirales; Podoviridae; ; ; | Enhanced biological phosphorus removal bioreactor viral communities from the University of Queensland, Australia - SBR4-V90806 Phage Sequencing | 6.00E-08 | 33/33 100% |
|  |  |  |  |  |  | CCAAAAGCCCTGTTCCACTCCGGTTGCAGTTCGCG | [IMGVR_UViG_3300013771_000004](https://img.jgi.doe.gov/cgi-bin/vr/main.cgi?section=ViralBrowse&page=uviginfo&uvig_id=IMGVR_UViG_3300013771_000004) | Duplodnaviria; Heunggongvirae; Uroviricota; Caudoviricetes; Caudovirales; ; ; ; | Activated sludge bacterial and viral communities from EBPR bioreactors in Brisbane, Australia - M92206 | 5.00E-09 | 35/35 100% |
|  |  |  |  |  |  | CTTCTGGCCGGGGTTGAGGCCGTCGTACTTGGCGG | [IMGVR_UViG_3300028804_000099](https://img.jgi.doe.gov/cgi-bin/vr/main.cgi?section=ViralBrowse&page=uviginfo&uvig_id=IMGVR_UViG_3300028804_000099) |  | Activated sludge microbial communities from WWTP in Nijmegen, Gelderland, Netherland - WWTP Weurt | 5.00E-09 | 35/35 100% |
| 29 | SCELSE-10IIF | SCELSE-10IIF Type III-A | III-A | 6 | 0 |  |  |  |  |  |  |
| 30 | AALB | AALB Type III-A | III-A | 50 | 0 |  |  |  |  |  |  |
| 31 | ACC012 | ACC012 Type III-A | III-A | 28 | 0 |  |  |  |  |  |  |
| 32 | UW1 | UW1 Type III-B | III-B | 63 | 1 | GTGTACGACGACATCCAAGGCGGCCCGGAAGGCGT | [IMGVR_UViG_3300005684_000006](https://img.jgi.doe.gov/cgi-bin/vr/main.cgi?section=ViralBrowse&page=uviginfo&uvig_id=IMGVR_UViG_3300005684_000006) | Duplodnaviria; Heunggongvirae; Uroviricota; Caudoviricetes; Caudovirales; ; ; ; | Enhanced biological phosphorus removal bioreactor viral communities from the University of Queensland, Australia - SBR4-V91805 Phage Sequencing | 5.00E-09 | 35/35 100% |
| 33 | UW5 | UW5 Type III-B | III-B | 7 | 0 |  |  |  |  |  |  |
| 34 | UW7 | UW7 Type III-B | III-B | 28 | 6 | CGGGGACTGCTGCTTCAGCACGAACGCGCTGA | [IMGVR_UViG_3300005675_000026](https://img.jgi.doe.gov/cgi-bin/vr/main.cgi?section=ViralBrowse&page=uviginfo&uvig_id=IMGVR_UViG_3300005675_000026) | Duplodnaviria; Heunggongvirae; Uroviricota; Caudoviricetes; Caudovirales; Podoviridae; ; Rauchvirus; | Enhanced biological phosphorus removal bioreactor viral communities from the University of Queensland, Australia - SBR4-V90806 Phage Sequencing | 2.00E-07 | 32/32 100% |
|  |  |  |  |  |  | TCCGATCGTCCGCTGCAAGGACTTGGCTCGTGG | [IMGVR_UViG_3300013771_000004](https://img.jgi.doe.gov/cgi-bin/vr/main.cgi?section=ViralBrowse&page=uviginfo&uvig_id=IMGVR_UViG_3300013771_000004) | Duplodnaviria; Heunggongvirae; Uroviricota;Caudoviricetes; Caudovirales; ; ; ; | Activated sludge bacterial and viral communities from EBPR bioreactors in Brisbane, Australia - M92206 | 6.00E-08 | 33/33 100% |
|  |  |  |  |  |  | GCTGTACCGGATTTACCGCTACTGGGGCAACA | [IMGVR_UViG_3300005684_000015](https://img.jgi.doe.gov/cgi-bin/vr/main.cgi?section=ViralBrowse&page=uviginfo&uvig_id=IMGVR_UViG_3300005684_000015) | Duplodnaviria; Heunggongvirae; Uroviricota; Caudoviricetes; Caudovirales; Podoviridae; ; Rauchvirus; | Enhanced biological phosphorus removal bioreactor viral communities from the University of Queensland, Australia - SBR4-V91805 Phage Sequencing | 2.00E-07 | 32/32 100% |
|  |  |  |  |  |  | GCAGGGCAGCAGCGGGCCGGGCGACTTGAACT | [IMGVR_UViG_3300013281_000030](https://img.jgi.doe.gov/cgi-bin/vr/main.cgi?section=ViralBrowse&page=uviginfo&uvig_id=IMGVR_UViG_3300013281_000030) | Duplodnaviria; Heunggongvirae; Uroviricota; Caudoviricetes; Caudovirales; Podoviridae; ; Rauchvirus; | Activated sludge bacterial and viral communities from EBPR bioreactors in Queensland, Australia - SBR4-V91307 | 2.00E-07 | 32/32 100% |
|  |  |  |  |  |  | ACCTTTCTGCCGTAGGAGAGCAGCATCCTATT | [IMGVR_UViG_3300014273_000005](https://img.jgi.doe.gov/cgi-bin/vr/main.cgi?section=ViralBrowse&page=uviginfo&uvig_id=IMGVR_UViG_3300014273_000005) |  | Activated sludge bacterial and viral communities from EBPR bioreactors in Queensland, Australia - SBR4-V91801 | 2.00E-07 | 32/32 100% |
|  |  |  |  |  |  | GGTTGCCGCCAGGATCAAGGCGGAGATTTCGC | [IMGVR_UViG_3300014059_000504](https://img.jgi.doe.gov/cgi-bin/vr/main.cgi?section=ViralBrowse&page=uviginfo&uvig_id=IMGVR_UViG_3300014059_000504) |  | Activated sludge microbial communities from Shanghai, China - membrane bioreactor - Membrane foulants | 2.00E-07 | 32/32 100% |
| 35 | UW9 | UW9 Type III-B | III-B | 63 | 1 | GTGTACGACGACATCCAAGGCGGCCCGGAAGGCGT | [IMGVR_UViG_3300005684_000006](https://img.jgi.doe.gov/cgi-bin/vr/main.cgi?section=ViralBrowse&page=uviginfo&uvig_id=IMGVR_UViG_3300005684_000006) | Duplodnaviria; Heunggongvirae; Uroviricota; Caudoviricetes; Caudovirales; ; ; ; | Enhanced biological phosphorus removal bioreactor viral communities from the University of Queensland, Australia - SBR4-V91805 Phage Sequencing | 3.00E-06 | 30/30 100% |
| 36 | UW12 | UW12 Type III-B | III-B | 16 | 0 |  |  |  |  |  |  |
| 37 | SCELSE-6IIF | SCELSE-6IIF Type III-B | III-B | 52 | 3 | GAAGATGGATTTTCGTTCGTCGCCGTCGTTCC | [IMGVR_UViG_3300005675_000013](https://img.jgi.doe.gov/cgi-bin/vr/main.cgi?section=ViralBrowse&page=uviginfo&uvig_id=IMGVR_UViG_3300005675_000013) | Duplodnaviria; Heunggongvirae; Uroviricota; Caudoviricetes; Caudovirales; ; ; ; | Enhanced biological phosphorus removal bioreactor viral communities from the University of Queensland, Australia - SBR4-V90806 Phage Sequencing | 2.00E-07 | 32/32 100% |
|  |  |  |  |  |  | GCCATCAGCGTGCCCTCCTCTTGACGTCGGGG | [IMGVR_UViG_3300005670_000031](https://img.jgi.doe.gov/cgi-bin/vr/main.cgi?section=ViralBrowse&page=uviginfo&uvig_id=IMGVR_UViG_3300005670_000031) | Duplodnaviria; Heunggongvirae; Uroviricota; Caudoviricetes; Caudovirales; ; ; ; | Enhanced biological phosphorus removal bioreactor viral communities from the University of Queensland, Australia - SBR4-V90104 Phage Sequencing | 2.00E-07 | 32/32 100% |
|  |  |  |  |  |  | CGTACAGACCATTCTGCCAGCGAACAGCAACCTT | [IMGVR_UViG_3300013281_000019](https://img.jgi.doe.gov/cgi-bin/vr/main.cgi?section=ViralBrowse&page=uviginfo&uvig_id=IMGVR_UViG_3300013281_000019) |  | Activated sludge bacterial and viral communities from EBPR bioreactors in Queensland, Australia - SBR4-V91307 | 2.00E-08 | 34/34 100% |
| 38 | SCELSE-4IIF | SCELSE-4IIF Type III-B | III-B | 52 | 3 | GAAGATGGATTTTCGTTCGTCGCCGTCGTTCC | [IMGVR_UViG_3300005675_000013](https://img.jgi.doe.gov/cgi-bin/vr/main.cgi?section=ViralBrowse&page=uviginfo&uvig_id=IMGVR_UViG_3300005675_000013) | Duplodnaviria; Heunggongvirae; Uroviricota; Caudoviricetes; Caudovirales; ; ; ; | Enhanced biological phosphorus removal bioreactor viral communities from the University of Queensland, Australia - SBR4-V90806 Phage Sequencing | 2.00E-07 | 32/32 100% |
|  |  |  |  |  |  | GCCATCAGCGTGCCCTCCTCTTGACGTCGGGG | [IMGVR_UViG_3300005670_000031](https://img.jgi.doe.gov/cgi-bin/vr/main.cgi?section=ViralBrowse&page=uviginfo&uvig_id=IMGVR_UViG_3300005670_000031) | Duplodnaviria; Heunggongvirae; Uroviricota; Caudoviricetes; Caudovirales; ; ; ; | Enhanced biological phosphorus removal bioreactor viral communities from the University of Queensland, Australia - SBR4-V90104 Phage Sequencing | 2.00E-07 | 32/32 100% |
|  |  |  |  |  |  | CGTACAGACCATTCTGCCAGCGAACAGCAACCTT | [IMGVR_UViG_3300013281_000019](https://img.jgi.doe.gov/cgi-bin/vr/main.cgi?section=ViralBrowse&page=uviginfo&uvig_id=IMGVR_UViG_3300013281_000019) |  | Activated sludge bacterial and viral communities from EBPR bioreactors in Queensland, Australia - SBR4-V91307 | 2.00E-08 | 34/34 100% |
| 39 | SCELSE-3IIF | SCELSE-3IIF Type III-B | III-B | 52 | 3 | GAAGATGGATTTTCGTTCGTCGCCGTCGTTCC | [IMGVR_UViG_3300005675_000013](https://img.jgi.doe.gov/cgi-bin/vr/main.cgi?section=ViralBrowse&page=uviginfo&uvig_id=IMGVR_UViG_3300005675_000013) | Duplodnaviria; Heunggongvirae; Uroviricota; Caudoviricetes; Caudovirales; ; ; ; | Enhanced biological phosphorus removal bioreactor viral communities from the University of Queensland, Australia - SBR4-V90806 Phage Sequencing | 2.00E-07 | 32/32 100% |
|  |  |  |  |  |  | GCCATCAGCGTGCCCTCCTCTTGACGTCGGGG | [IMGVR_UViG_3300005670_000031](https://img.jgi.doe.gov/cgi-bin/vr/main.cgi?section=ViralBrowse&page=uviginfo&uvig_id=IMGVR_UViG_3300005670_000031) | Duplodnaviria; Heunggongvirae; Uroviricota; Caudoviricetes; Caudovirales; ; ; ; | Enhanced biological phosphorus removal bioreactor viral communities from the University of Queensland, Australia - SBR4-V90104 Phage Sequencing | 2.00E-07 | 32/32 100% |
|  |  |  |  |  |  | CGTACAGACCATTCTGCCAGCGAACAGCAACCTT | [IMGVR_UViG_3300013281_000019](https://img.jgi.doe.gov/cgi-bin/vr/main.cgi?section=ViralBrowse&page=uviginfo&uvig_id=IMGVR_UViG_3300013281_000019) |  | Activated sludge bacterial and viral communities from EBPR bioreactors in Queensland, Australia - SBR4-V91307 | 2.00E-08 | 34/34 100% |
| 40 | ACC003 | ACC003 Type III-B | III-B | 35 | 0 |  |  |  |  |  |  |

**Table S4.** DRs in each CRISPR-Cas locus compared to the CRISPRCasdb database.

| **Number** | **Strains** | **CRISPR loci** | **Maker Cas gene clade** | **Direct repeat** | **Strain(s) having this sequence** | **Strain's sequence(s)** | **Nucleotides match** | **Blast E-value** |
| --- | --- | --- | --- | --- | --- | --- | --- | --- |
| Type I-A | | | | | | | | |
| 1 | UW6 | UW6 Type I-A | Cas1 cladeII | GTGCCGTCATCTTTGATGCCGAAAGGCGTTGAGCAC | Leptospira kmetyi (bacteria) LS 001/16 | CP033614.1 | 26/36 | 2.00E-09 |
| 2 | Bin19 | Bin19 Type I-A | Cas1 cladeII | GTGCCGTCATCTTTGATGCCGAAAGGCGTTGAGCAC | Leptospira kmetyi (bacteria) LS 001/16 | CP033614.1 | 26/36 | 2.00E-09 |
| 3 | SCELSE-2IIC | SCELSE-2IIC Type I-A | Cas1 cladeII | GTGCCGTCATCTTTGATGCCGAAAGGCGTTGAGCAC | Leptospira kmetyi (bacteria) LS 001/16 | [CP033614.1](https://crisprcas.i2bc.paris-saclay.fr/MainDb/StrainList/CP033614.1) | 26/36 | 3.00E-09 |
| Type I-C | | | | | | | | |
| 4 | SSA1 | SSA1 Type I-C | Cas1 cladeIII | GTCGCGCGCTCCTCACGGGGCGCGCGTGGATTGAAAC | Xanthomonas oryzae pv. oryzae (g-proteobacteria) T7133 | CP071891.1 | 18/31 | 2e-04 |
| 5 | SSB1 | SSB1 Type I-C | Cas1 cladeIII | GGTCGCGCGCCCCGCGAGGGGCGCGCGTGGATTGAAAC | Planctomycetes bacterium Enr13 (bacteria) | CP037423.1 | 18/32 | 2e-04 |
| 6 | DS2011 | DS2011 Type I-C | Cas1 cladeIV | GCTTCGCCCTTCGGCAACGAGGGGCGTGGATTGAAAC | Denitratisoma sp. DHT3 (b-proteobacteria) | CP020914.1 | 31/37 | 1.00E-08 |
| 7 | MAG-196 | MAG-196 Type I-C | Cas1 cladeIV | GCTTCGCCCCTCTGCAACGGGGGGCGTGGATTGAAAC | Ferribacterium limneticum (b-proteobacteria) 128 | [CP075191.1](https://crisprcas.i2bc.paris-saclay.fr/MainDb/StrainList/CP075191.1) | 32/37 | 3.00E-09 |
| 8 | AALB | AALB Type I-C | Cas1 cladeIV | GCATCGCCCCTCGGCAACGAGGGGCGCGGATTGAAAC | Ferribacterium limneticum (b-proteobacteria) 128 | [CP075191.1](https://crisprcas.i2bc.paris-saclay.fr/MainDb/StrainList/CP075191.1) | 34/37 | 1.00E-08 |
| 9 | ACC012 | ACC012 Type I-C | Cas1 cladeIV | GACACGCTCCCCGGCGACGGGGAGCGAGGATTGAAAC | Ferribacterium limneticum (b-proteobacteria) 128 | [CP075191.1](https://crisprcas.i2bc.paris-saclay.fr/MainDb/StrainList/CP075191.1) | 28/37 | 7.00E-07 |
| Type I-E | | | | | | | | |
| 10 | UW6 | UW6 Type I-E | Cas1 clade V | GTGTTCCCCGCAGGCGCGGGGATGAACCG | Acidiphilium multivorum (a-proteobacteria) JZ-6 | [CP041716.1](https://crisprcas.i2bc.paris-saclay.fr/MainDb/StrainList/CP041716.1) | 29/29 | 3.00E-11 |
|  |  |  |  |  | Geobacter sulfurreducens (d-proteobacteria) AM-1 | [CP010430.1](https://crisprcas.i2bc.paris-saclay.fr/MainDb/StrainList/CP010430.1) | 29/29 | 3.00E-11 |
|  |  |  |  |  | Pseudomonas balearica DSM 6083 (g-proteobacteria) DSM6083 (=SP1402) | [CP007511.1](https://crisprcas.i2bc.paris-saclay.fr/MainDb/StrainList/CP007511.1) | 29/29 | 3.00E-11 |
|  |  |  |  |  | Roseomonas sp. 1318 (a-proteobacteria) | [CP061096.1](https://crisprcas.i2bc.paris-saclay.fr/MainDb/StrainList/CP061096.1) | 29/29 | 3.00E-11 |
|  |  |  |  |  | Acidiphilium cryptum JF-5 (a-proteobacteria) | [CP000690.1](https://crisprcas.i2bc.paris-saclay.fr/MainDb/StrainList/CP000690.1) | 29/29 | 3.00E-11 |
|  |  |  |  |  | Acidiphilium multivorum (a-proteobacteria) JZ-6 | [CP041719.1](https://crisprcas.i2bc.paris-saclay.fr/MainDb/StrainList/CP041719.1) | 29/29 | 3.00E-11 |
|  |  |  |  |  | Pseudomonas balearica (g-proteobacteria) EC28 | [CP045858.1](https://crisprcas.i2bc.paris-saclay.fr/MainDb/StrainList/CP045858.1) | 29/29 | 3.00E-11 |
|  |  |  |  |  | Pseudomonas balearica (g-proteobacteria) FDAARGOS_1013 | [CP067013.1](https://crisprcas.i2bc.paris-saclay.fr/MainDb/StrainList/CP067013.1) | 29/29 | 3.00E-11 |
| 11 | BA-93 | BA-93 Type I-E | Cas1 clade V | GCGTTCCCCGCAGGCGCGGGGATGAACCG | Acidiphilium cryptum JF-5 (a-proteobacteria) | CP000690.1 | 29/29 | 3.00E-11 |
|  |  |  |  |  | Pseudomonas balearica (g-proteobacteria) EC28 | CP045858.1 | 29/29 | 3.00E-11 |
|  |  |  |  |  | Pseudomonas balearica (g-proteobacteria) FDAARGOS_1013 | CP067013.1 | 29/29 | 3.00E-11 |
|  |  |  |  |  | Kushneria marisflavi (g-proteobacteria) SW32 | CP021358.1 | 29/29 | 3.00E-11 |
|  |  |  |  |  | Pseudomonas balearica DSM 6083 (g-proteobacteria) DSM6083 (=SP1402) | CP007511.1 | 29/29 | 3.00E-11 |
|  |  |  |  |  | Salinisphaera sp. LB1 (g-proteobacteria) | CP029488.1 | 29/29 | 3.00E-11 |
| 12 | ACC005 | ACC005 Type I-E | Cas1 clade V | GCGTTCCCCGCAGGCGCGGGGATGAACCG | Acidiphilium cryptum JF-5 (a-proteobacteria) | [CP000690.1](https://crisprcas.i2bc.paris-saclay.fr/MainDb/StrainList/CP000690.1) | 29/29 | 3.00E-11 |
|  |  |  |  |  | Pseudomonas balearica (g-proteobacteria) EC28 | [CP045858.1](https://crisprcas.i2bc.paris-saclay.fr/MainDb/StrainList/CP045858.1) | 29/29 | 3.00E-11 |
|  |  |  |  |  | Pseudomonas balearica (g-proteobacteria) FDAARGOS_1013 | [CP067013.1](https://crisprcas.i2bc.paris-saclay.fr/MainDb/StrainList/CP067013.1) | 29/29 | 3.00E-11 |
|  |  |  |  |  | Kushneria marisflavi (g-proteobacteria) SW32 | [CP021358.1](https://crisprcas.i2bc.paris-saclay.fr/MainDb/StrainList/CP021358.1) | 29/29 | 3.00E-11 |
|  |  |  |  |  | Pseudomonas balearica DSM 6083 (g-proteobacteria) DSM6083 (=SP1402) | [CP007511.1](https://crisprcas.i2bc.paris-saclay.fr/MainDb/StrainList/CP007511.1) | 29/29 | 3.00E-11 |
|  |  |  |  |  | Salinisphaera sp. LB1 (g-proteobacteria) | [CP029488.1](https://crisprcas.i2bc.paris-saclay.fr/MainDb/StrainList/CP029488.1) | 29/29 | 3.00E-11 |
| 13 | ACC007 | ACC007 Type I-E | Cas1 clade V | GCGTTCCCCGCAGGCGCGGGGATGAACCG | Acidiphilium cryptum JF-5 (a-proteobacteria) | [CP000690.1](https://crisprcas.i2bc.paris-saclay.fr/MainDb/StrainList/CP000690.1) | 29/29 | 3.00E-11 |
|  |  |  |  |  | Pseudomonas balearica (g-proteobacteria) EC28 | [CP045858.1](https://crisprcas.i2bc.paris-saclay.fr/MainDb/StrainList/CP045858.1) | 29/29 | 3.00E-11 |
|  |  |  |  |  | Pseudomonas balearica (g-proteobacteria) FDAARGOS_1013 | [CP067013.1](https://crisprcas.i2bc.paris-saclay.fr/MainDb/StrainList/CP067013.1) | 29/29 | 3.00E-11 |
|  |  |  |  |  | Kushneria marisflavi (g-proteobacteria) SW32 | [CP021358.1](https://crisprcas.i2bc.paris-saclay.fr/MainDb/StrainList/CP021358.1) | 29/29 | 3.00E-11 |
|  |  |  |  |  | Pseudomonas balearica DSM 6083 (g-proteobacteria) DSM6083 (=SP1402) | [CP007511.1](https://crisprcas.i2bc.paris-saclay.fr/MainDb/StrainList/CP007511.1) | 29/29 | 3.00E-11 |
|  |  |  |  |  | Salinisphaera sp. LB1 (g-proteobacteria) | [CP029488.1](https://crisprcas.i2bc.paris-saclay.fr/MainDb/StrainList/CP029488.1) | 29/29 | 3.00E-11 |
| 14 | UW1 | UW1 Type I-E | Cas1 clade VI | GTTTCCCCCGCGTCAGCGGGGATAGGCCC | Aromatoleum bremense (b-proteobacteria) PbN1 | [CP059467.1](https://crisprcas.i2bc.paris-saclay.fr/MainDb/StrainList/CP059467.1) | 28/29 | 8.00E-09 |
| 15 | UW5 | UW5 Type I-E | Cas1 clade VI | GTTTCCCCCGCGTCAGCGGGGATAGGCCC | Aromatoleum bremense (b-proteobacteria) PbN1 | [CP059467.1](https://crisprcas.i2bc.paris-saclay.fr/MainDb/StrainList/CP059467.1) | 28/29 | 8.00E-09 |
| 16 | UW9 | UW9 Type I-E | Cas1 clade VI | GTTTCCCCCGCGTCAGCGGGGATAGGCCC | Aromatoleum bremense (b-proteobacteria) PbN1 | [CP059467.1](https://crisprcas.i2bc.paris-saclay.fr/MainDb/StrainList/CP059467.1) | 28/29 | 8.00E-09 |
| 17 | MAXAC027 | MAXAC027 Type I-E | Cas1 clade VI | GTTTCCCCCGCGCCAGCGGGGATAGGCCC | Nitrosospira lacus (b-proteobacteria) APG3 | [CP021106.3](https://crisprcas.i2bc.paris-saclay.fr/MainDb/StrainList/CP021106.3) | 27/28 | 3.00E-08 |
| 18 | ACC003 | ACC003 Type I-E | Cas1 clade VI | GTTTCCCCCGCGCCAGCGGGGATAGGCCC | Nitrosospira lacus (b-proteobacteria) APG3 | [CP021106.3](https://crisprcas.i2bc.paris-saclay.fr/MainDb/StrainList/CP021106.3) | 27/28 | 3.00E-08 |
| Type I-F | | | | | | | | |
| 19 | SCELSE-8IIC | Cas1 clade I |  | GTTCACTGCCGCACAGGCAGCTCAGAAA | Desulfurivibrio alkaliphilus AHT 2 (d-proteobacteria) | [CP001940.1](https://crisprcas.i2bc.paris-saclay.fr/MainDb/StrainList/CP001940.1) | 28/28 | 1.00E-10 |
|  |  |  |  |  | Halomonas sp. Y2R2 (g-proteobacteria) | [CP038437.2](https://crisprcas.i2bc.paris-saclay.fr/MainDb/StrainList/CP038437.2) | 28/28 | 1.00E-10 |
|  |  |  |  |  | Pseudoalteromonas tunicata (g-proteobacteria) D2 | [CP011032.1](https://crisprcas.i2bc.paris-saclay.fr/MainDb/StrainList/CP011032.1) | 28/28 | 1.00E-10 |
|  |  |  |  |  | Pseudoalteromonas tunicata (g-proteobacteria) D2 89e3fb8 | [CP031961.1](https://crisprcas.i2bc.paris-saclay.fr/MainDb/StrainList/CP031961.1) | 28/28 | 1.00E-10 |
|  |  |  |  |  | Shewanella inventionis (g-proteobacteria) D1489 | [CP082926.1](https://crisprcas.i2bc.paris-saclay.fr/MainDb/StrainList/CP082926.1) | 28/28 | 1.00E-10 |
| 20 | SCUT-2 | SCUT-2 Type I-F | Cas1 clade I | GTTCACTGCCGCACAGGCAGCTCAGAAA | Desulfurivibrio alkaliphilus AHT 2 (d-proteobacteria) | [CP001940.1](https://crisprcas.i2bc.paris-saclay.fr/MainDb/StrainList/CP001940.1) | 28/28 | 1.00E-10 |
|  |  |  |  |  | Halomonas sp. Y2R2 (g-proteobacteria) | [CP038437.2](https://crisprcas.i2bc.paris-saclay.fr/MainDb/StrainList/CP038437.2) | 28/28 | 1.00E-10 |
|  |  |  |  |  | Pseudoalteromonas tunicata (g-proteobacteria) D2 | [CP011032.1](https://crisprcas.i2bc.paris-saclay.fr/MainDb/StrainList/CP011032.1) | 28/28 | 1.00E-10 |
|  |  |  |  |  | Pseudoalteromonas tunicata (g-proteobacteria) D2 89e3fb8 | [CP031961.1](https://crisprcas.i2bc.paris-saclay.fr/MainDb/StrainList/CP031961.1) | 28/28 | 1.00E-10 |
|  |  |  |  |  | Shewanella inventionis (g-proteobacteria) D1489 | [CP082926.1](https://crisprcas.i2bc.paris-saclay.fr/MainDb/StrainList/CP082926.1) | 28/28 | 1.00E-10 |
| Type III-A | | | | | | | | |
| 21 | MAXAC027 | MAXAC027 Type III-A | Cas10 clade I | GTAGAAAACCAGCCCTGATTTCTAAGGGATTAAGAC | Rhodoferax antarcticus (b-proteobacteria) DSM 24876 | [CP019240.1](https://crisprcas.i2bc.paris-saclay.fr/MainDb/StrainList/CP019240.1) | 27/37 | 3.00E-06 |
| 22 | SCELSE-10IIF | SCELSE-10IIF Type III-A | Cas10 clade I | GTAGCAACCCAGCCCTGATTACGAAGGGATTAAGAC | Methylomonas sp. DH-1 (g-proteobacteria) | [CP014360.1](https://crisprcas.i2bc.paris-saclay.fr/MainDb/StrainList/CP014360.1) | 25/36 | 4.00E-05 |
| 23 | BATAC285 | BATAC285 Type III-A | Cas10 clade II | GTCAGTAACCAGCCCTGTAATCAAAGGGATTGAGAC | Tepidimonas taiwanensis (b-proteobacteria) LMG 22826 | [CP083911.1](https://crisprcas.i2bc.paris-saclay.fr/MainDb/StrainList/CP083911.1) | 25/36 | 7.00E-07 |
| 24 | AALB | AALB Type III-A | Cas10 clade II | GTCAGTAACCAGCCCTGAAATCAAAGGGATTGAGAC | Tepidimonas taiwanensis (b-proteobacteria) LMG 22826 | [CP083911.1](https://crisprcas.i2bc.paris-saclay.fr/MainDb/StrainList/CP083911.1) | 26/36 | 3.00E-09 |
| 25 | UWLDOIC | UWLDOIC Type III-A | Cas10 clade II | GTCAGTAACCAGCCCTGAAATCAAAGGGATTGAGAC | Tepidimonas taiwanensis (b-proteobacteria) LMG 22826 | [CP083911.1](https://crisprcas.i2bc.paris-saclay.fr/MainDb/StrainList/CP083911.1) | 25/36 | 7.00E-07 |
| 26 | ACC012 | ACC012 Type III-A | Cas10 clade II | GTCAGTAACCAGCCCTGAAATCAAAGGGATTGAGAC | Tepidimonas taiwanensis (b-proteobacteria) LMG 22826 | [CP083911.1](https://crisprcas.i2bc.paris-saclay.fr/MainDb/StrainList/CP083911.1) | 26/36 | 3.00E-09 |
| 27 | UW7 | UW7 Type III-A | Cas10 clade III | GTCAGTATGCAGCCCTGAAAACAAAGGGATTGAGAC | Tepidimonas taiwanensis (b-proteobacteria) LMG 22826 | [CP083911.1](https://crisprcas.i2bc.paris-saclay.fr/MainDb/StrainList/CP083911.1) | 25/36 | 7.00E-07 |
| 28 | UW13 | UW13 Type III-A | Cas10 clade III | GTCAGTATGCAGCCCTGAAAACAAAGGGATTGAGAC | Tepidimonas taiwanensis (b-proteobacteria) LMG 22826 | [CP083911.1](https://crisprcas.i2bc.paris-saclay.fr/MainDb/StrainList/CP083911.1) | 25/36 | 7.00E-07 |
| 29 | SCELSE-4IIF | SCELSE-4IIF Type III-A | Cas10 clade III | GTCAGTAGCCAGCCCTGAAATCAAAGGGATTAAGAC | Rhodoferax antarcticus (b-proteobacteria) DSM 24876 | [CP019240.1](https://crisprcas.i2bc.paris-saclay.fr/MainDb/StrainList/CP019240.1) | 27/37 | 5.00E-08 |
| 30 | SCELSE-3IIF | SCELSE-3IIF Type III-A | Cas10 clade III | GTCAGTAGCCAGCCCTGAAATCAAAGGGATTAAGAC | Rhodoferax antarcticus (b-proteobacteria) DSM 24876 | [CP019240.1](https://crisprcas.i2bc.paris-saclay.fr/MainDb/StrainList/CP019240.1) | 27/37 | 5.00E-08 |
| 31 | SCELSE-6IIF | SCELSE-6IIF Type III-A | Cas10 clade III | GTCAGTAGCCAGCCCTGAAATCAAAGGGATTAAGAC | Rhodoferax antarcticus (b-proteobacteria) DSM 24876 | [CP019240.1](https://crisprcas.i2bc.paris-saclay.fr/MainDb/StrainList/CP019240.1) | 27/37 | 5.00E-08 |
| Type III-B | | | | | | | | |
| 32 | UW7 | UW7 Type III-B | Cas10 clade IV | GTTCTCTCTCCCCGAATTCCTGAAGGGGATTAAGAC | Ferrovum myxofaciens (b-proteobacteria) | [CP053675.2](https://crisprcas.i2bc.paris-saclay.fr/MainDb/StrainList/CP053675.2) | 26/36 | 2.00E-07 |
| 33 | SCELSE-6IIF | SCELSE-6IIF Type III-B | Cas10 clade IV | GTTCTCTCTCCCCGAATTCCTGAAGGGGATTAAGAC | Ferrovum myxofaciens (b-proteobacteria) | [CP053675.2](https://crisprcas.i2bc.paris-saclay.fr/MainDb/StrainList/CP053675.2) | 26/36 | 2.00E-07 |
| 34 | SCELSE-3IIF | SCELSE-3IIF Type IIIB | Cas10 clade IV | GTTCTCTCTCCCCGAATTCCTGAAGGGGATTAAGAC | Ferrovum myxofaciens (b-proteobacteria) | [CP053675.2](https://crisprcas.i2bc.paris-saclay.fr/MainDb/StrainList/CP053675.2) | 26/36 | 2.00E-07 |
| 35 | SCELSE-4IIF | SCELSE-4IIF Type IIIB | Cas10 clade IV | GTTCTCTCTCCCCGAATTCCTGAAGGGGATTAAGAC | Ferrovum myxofaciens (b-proteobacteria) | [CP053675.2](https://crisprcas.i2bc.paris-saclay.fr/MainDb/StrainList/CP053675.2) | 26/36 | 2.00E-07 |
| 36 | ACC003 | ACC003 Type IIIB | Cas10 clade V | GTCTCAATCCCTTTGATTTCAGGGCTGGTTACTGAC | Tepidimonas taiwanensis (b-proteobacteria) LMG 22826Bacteria | CP083911.1 | 26/36 | 3.00E-09 |
| 37 | UW12 | UW12 Type IIIB | Cas10 clade V | GTCAGTAACCAGCCCTGAAAACAAAGGGATTGAGAC | Tepidimonas taiwanensis (b-proteobacteria) LMG 22826 | [CP083911.1](https://crisprcas.i2bc.paris-saclay.fr/MainDb/StrainList/CP083911.1) | 25/36 | 7.00E-07 |
| 38 | UW1 | UW1 Type IIIB | Cas10 clade V | GTCAGTAACCAGCCCTGAAATCAAAGGGATTGAGAC | Tepidimonas taiwanensis (b-proteobacteria) LMG 22826 | [CP083911.1](https://crisprcas.i2bc.paris-saclay.fr/MainDb/StrainList/CP083911.1) | 26/36 | 3.00E-09 |
| 39 | UW5 | UW5 Type IIIB | Cas10 clade V | GTCAGTAACCAGCCCTGAAATCAAAGGGATTGAGAC | Tepidimonas taiwanensis (b-proteobacteria) LMG 22826 | [CP083911.1](https://crisprcas.i2bc.paris-saclay.fr/MainDb/StrainList/CP083911.1) | 26/36 | 3.00E-09 |
| 40 | UW9 | UW9 Type IIIB | Cas10 clade V | GTCAGTAACCAGCCCTGAAATCAAAGGGATTGAGAC | Tepidimonas taiwanensis (b-proteobacteria) LMG 22826 | [CP083911.1](https://crisprcas.i2bc.paris-saclay.fr/MainDb/StrainList/CP083911.1) | 26/36 | 3.00E-09 |

Table S5. DRs in each CRISPR-Cas locus compared to the CRISPRCasdb database

| **CRISPR loci** | **Spacers** | **Target phage genomes** |
| --- | --- | --- |
| SCELSE-3IIF Type III-A | CCGCCAAGTACGACGGCCTCAACCCCGGCCAGAAG | phage2_SCUT-2 |
|  | CGCGAACTGCAACCGGAGTGGAACAGGGCTTTTGG | phage8_NTU30-0125 |
|  | CCGAGGATGGCTCCCCGTGCATCGTGTCGGCGATG | phage22_NTU30-0823 |
|  | CCGAGGATGGCTCCCCGTGCATCGTGTCGGCGATG | phage19_NTU30-0802 |
|  | CTGGTGGCAGCGTGAGCGACGGCGTTCTGACCCG | phage22_NTU30-0823 |
|  | CTGGTGGCAGCGTGAGCGACGGCGTTCTGACCCG | phage19_NTU30-0802 |
|  | CTGGTGGCAGCGTGAGCGACGGCGTTCTGACCCG | phage17_NTU30-0515 |
|  | CTGGTGGCAGCGTGAGCGACGGCGTTCTGACCCG | phage14_NTU30-0125 |
|  | GTCACCGTGCTGGTGTACGCGCTCATCGACACCGT | phage24_NTU35-0823 |
|  | GTCACCGTGCTGGTGTACGCGCTCATCGACACCGT | phage12_NTU30-0125 |
| SCELSE-3IIF Type III-B | GAAGATGGATTTTCGTTCGTCGCCGTCGTTCC | phage15_NTU30-0823 |
|  | GAAGATGGATTTTCGTTCGTCGCCGTCGTTCC | phage15_NTU30-0802 |
|  | GAAGATGGATTTTCGTTCGTCGCCGTCGTTCC | phage15_NTU30-0515 |
|  | GAAGATGGATTTTCGTTCGTCGCCGTCGTTCC | phage10_NTU30-0125 |
|  | CTCTCCGCGTTTGCTTTTGTCTGCGCATCATCAA | phage21_NTU30-0802 |
|  | CTCTCCGCGTTTGCTTTTGTCTGCGCATCATCAA | phage7_NTU30-0125 |
| SCELSE-10IIF Type III-A | None |  |
| SCELSE-4IIF Type III-A | ACGGTGTCGATGAGCGCGTACACCAGCACGGTGAC | phage24_NTU35-0823 |
|  | ACGGTGTCGATGAGCGCGTACACCAGCACGGTGAC | phage12_NTU30-0125 |
|  | CGGGTCAGAACGCCGTCGCTCACGCTGCCACCAG | phage22_NTU30-0823 |
|  | CGGGTCAGAACGCCGTCGCTCACGCTGCCACCAG | phage19_NTU30-0802 |
|  | CGGGTCAGAACGCCGTCGCTCACGCTGCCACCAG | phage17_NTU30-0515 |
|  | CGGGTCAGAACGCCGTCGCTCACGCTGCCACCAG | phage14_NTU30-0125 |
|  | CATCGCCGACACGATGCACGGGGAGCCATCCTCGG | phage22_NTU30-0823 |
|  | CATCGCCGACACGATGCACGGGGAGCCATCCTCGG | phage19_NTU30-0802 |
|  | CCAAAAGCCCTGTTCCACTCCGGTTGCAGTTCGCG | phage8_NTU30-0125 |
|  | CTTCTGGCCGGGGTTGAGGCCGTCGTACTTGGCGG | phage2_SCUT-2 |
| SCELSE-4IIF Type III-B | GAAGATGGATTTTCGTTCGTCGCCGTCGTTCC | phage15_NTU30-0823 |
|  | GAAGATGGATTTTCGTTCGTCGCCGTCGTTCC | phage15_NTU30-0802 |
|  | GAAGATGGATTTTCGTTCGTCGCCGTCGTTCC | phage15_NTU30-0515 |
|  | GAAGATGGATTTTCGTTCGTCGCCGTCGTTCC | phage10_NTU30-0125 |
|  | CTCTCCGCGTTTGCTTTTGTCTGCGCATCATCAA | phage21_NTU30-0802 |
|  | CTCTCCGCGTTTGCTTTTGTCTGCGCATCATCAA | phage7_NTU30-0125 |
| SCELSE-8IIC Type I-F | ATGCAGTCTATCGGGTGATGAGTGGTGCTAGA | phage16_NTU30-0515 |
|  | ATGCAGTCTATCGGGTGATGAGTGGTGCTAGA | phage11_NTU30-0125 |
|  | ATTCCCGGCCGATCCGAACTGCTCCGGGAATT | phage3_SCUT-2 |
|  | ACGCAAAATATCCTGGCCGAACCTCGCGGAGT | phage3_SCUT-2 |
|  | AGGGGAAAGAGATCCTGACGACGGGCAAATCC | phage1_SCUT-2 |
|  | TCTGCAGCAACGACGAAACCCCCTGCGCCCTC | phage4_SCUT-2 |
| SCELSE-6IIF Type III-A | ACGGTGTCGATGAGCGCGTACACCAGCACGGTGAC | phage24_NTU35-0823 |
|  | ACGGTGTCGATGAGCGCGTACACCAGCACGGTGAC | phage12_NTU30-0125 |
|  | CGGGTCAGAACGCCGTCGCTCACGCTGCCACCAG | phage22_NTU30-0823 |
|  | CGGGTCAGAACGCCGTCGCTCACGCTGCCACCAG | phage19_NTU30-0802 |
|  | CGGGTCAGAACGCCGTCGCTCACGCTGCCACCAG | phage17_NTU30-0515 |
|  | CGGGTCAGAACGCCGTCGCTCACGCTGCCACCAG | phage14_NTU30-0125 |
|  | CATCGCCGACACGATGCACGGGGAGCCATCCTCGG | phage22_NTU30-0823 |
|  | CATCGCCGACACGATGCACGGGGAGCCATCCTCGG | phage19_NTU30-0802 |
|  | CCAAAAGCCCTGTTCCACTCCGGTTGCAGTTCGCG | phage8_NTU30-0125 |
|  | CTTCTGGCCGGGGTTGAGGCCGTCGTACTTGGCGG | phage2_SCUT-2 |
| SCELSE-6IIF Type III-B | TTGATGATGCGCAGACAAAAGCAAACGCGGAGAG | phage21_NTU30-0802 |
|  | TTGATGATGCGCAGACAAAAGCAAACGCGGAGAG | phage7_NTU30-0125 |
|  | GGAACGACGGCGACGAACGAAAATCCATCTTC | phage15_NTU30-0823 |
|  | GGAACGACGGCGACGAACGAAAATCCATCTTC | phage15_NTU30-0802 |
|  | GGAACGACGGCGACGAACGAAAATCCATCTTC | phage15_NTU30-0515 |
|  | GGAACGACGGCGACGAACGAAAATCCATCTTC | phage10_NTU30-0125 |
| SCELSE-2IIC Type I-A | TGGTAGTCGATCACCCAGGTCTTGCCGCCTTTGCCG | phage3_SCUT-2 |
| SCUT-2 Type I-F | TCGTCCTGATTGGCGCGTTCGGCTATTCGGTC | phage4_SCUT-2 |
|  | TCGCTGAGGTGGTCCATGATTTCGCGCAGATA | phage3_SCUT-2 |
|  | TTGAAGCTGCGCAACTGGGCGCGTGCGTGGAT | phage23_NTU30-0823 |
|  | TTGAAGCTGCGCAACTGGGCGCGTGCGTGGAT | phage20_NTU30-0802 |
|  | TAGAGCACGACGAAGCCTTACGGCTACCAAGT | phage23_NTU30-0823 |
|  | TAGAGCACGACGAAGCCTTACGGCTACCAAGT | phage20_NTU30-0802 |
|  | GCGCTTCGTCGATCACGATCAGATCCACGTCC | phage1_SCUT-2 |
|  | CGATCACCTCGAACAGCATGACCTCGGCGACT | phage9_NTU30-0125 |
|  | ATCTCGCGCACCCGTGCCCGCTGTTCTTCAAT | phage18_NTU30-0515 |
|  | CCAACGCTGCGCGAAGAGGACTCGGAGTTGTG | phage1_SCUT-2 |
|  | CTCGAACGCAACATAGACAGCATCAAGTCCAC | phage1_SCUT-2 |
|  | TCCTTGTCCCGCAGTCGGTCCCGGTGCAGTTT | phage1_SCUT-2 |
|  | TTCGAGGCCGGTGAGAAATTTTGGCCGAGTGC | phage5_SCUT-2 |
|  | TTCGAGGCCGGTGAGAAATTTTGGCCGAGTGC | phage13_NTU30-0125 |
|  | TCTTTCTTGATTTCGGCATGGACGGTATGCGC | phage4_SCUT-2 |
|  | AGCAGGTATTCCCGCCAGACCGCGGAACTGCA | phage6_SCUT-2 |
|  | AGCAGGTATTCCCGCCAGACCGCGGAACTGCA | phage13_NTU30-0125 |
|  | GGCATCCGTCCTGCGGTCGGCTTGTGCGTGAT | phage13_NTU30-0125 |

Table S6. Phage genomes retrieved from the SCUT and NTU SBRs compared to the IMG/VR database

| **Phage code** | **Query sequence name** | **UViGs** | **Lineage** | **Identities** | **E-value** |
| --- | --- | --- | --- | --- | --- |
| phage1_SCUT-2 | SCUT-2_817_l_44919_cov_13.698266full | [IMGVR_UViG_3300013800_000258](https://img.jgi.doe.gov/cgi-bin/vr/main.cgi?section=ViralBrowse&page=uviginfo&uvig_id=IMGVR_UViG_3300013800_000258) | Duplodnaviria; Heunggongvirae; Uroviricota; Caudoviricetes; Caudovirales; ; ; ; | 5099/5313 96% | 0 |
| phage2_SCUT-2 | SCUT-2_950_l_40164_cov_44.658655full | [IMGVR_UViG_3300029936_000085](https://img.jgi.doe.gov/cgi-bin/vr/main.cgi?section=ViralBrowse&page=uviginfo&uvig_id=IMGVR_UViG_3300029936_000085) | Duplodnaviria; Heunggongvirae; Uroviricota; Caudoviricetes; Caudovirales; Podoviridae; ; ; | 11792/12241 96% | 0 |
| phage3_SCUT-2 | SCUT-2_1793_l_24214_cov_135.145122full | [IMGVR_UViG_3300028804_000035](https://img.jgi.doe.gov/cgi-bin/vr/main.cgi?section=ViralBrowse&page=uviginfo&uvig_id=IMGVR_UViG_3300028804_000035) | Duplodnaviria; Heunggongvirae; Uroviricota; Caudoviricetes; Caudovirales; ; ; ; | 7835/8539 92% | 0 |
| phage4_SCUT-2 | SCUT-2_3483_l_12823_cov_141.257597full | [IMGVR_UViG_3300028804_000035](https://img.jgi.doe.gov/cgi-bin/vr/main.cgi?section=ViralBrowse&page=uviginfo&uvig_id=IMGVR_UViG_3300028804_000035) | Duplodnaviria; Heunggongvirae; Uroviricota; Caudoviricetes; Caudovirales; ; ; ; | 2858/2953 97% | 0 |
| phage5_SCUT-2 | SCUT-2_4730_l_9642_cov_27.970168full | [IMGVR_UViG_3300022549_000013](https://img.jgi.doe.gov/cgi-bin/vr/main.cgi?section=ViralBrowse&page=uviginfo&uvig_id=IMGVR_UViG_3300022549_000013) | Duplodnaviria; Heunggongvirae; Uroviricota; Caudoviricetes; Caudovirales; ; ; ; | 1975/2580 77% | 0 |
| phage6_SCUT-2 | SCUT-2_17027_l_3247_cov_28.407268full | [IMGVR_UViG_3300013771_000016](https://img.jgi.doe.gov/cgi-bin/vr/main.cgi?section=ViralBrowse&page=uviginfo&uvig_id=IMGVR_UViG_3300013771_000016) | Duplodnaviria; Heunggongvirae; Uroviricota; Caudoviricetes; Caudovirales; Siphoviridae; ; ; | 2593/3252 80% | 0 |
| phage7_NTU30-0125 | NTU30-0125_966_l_44019_cov_198.457238full | [IMGVR_UViG_3300005682_000004](https://img.jgi.doe.gov/cgi-bin/vr/main.cgi?section=ViralBrowse&page=uviginfo&uvig_id=IMGVR_UViG_3300005682_000004) |  | 7101/7986 89% | 0 |
| phage8_NTU30-0125 | NTU30-0125_1064_l_42016_cov_2055.839970full | [IMGVR_UViG_3300013771_000004](https://img.jgi.doe.gov/cgi-bin/vr/main.cgi?section=ViralBrowse&page=uviginfo&uvig_id=IMGVR_UViG_3300013771_000004) | Duplodnaviria; Heunggongvirae; Uroviricota; Caudoviricetes; Caudovirales; ; ; ; | 3953/4193 94% | 0 |
| phage9_NTU30-0125 | NTU30-0125_1524_l_33728_cov_420.111009full | [IMGVR_UViG_3300009673_000100](https://img.jgi.doe.gov/cgi-bin/vr/main.cgi?section=ViralBrowse&page=uviginfo&uvig_id=IMGVR_UViG_3300009673_000100) |  | 8008/8097 99% | 0 |
| phage10_NTU30-0125 | NTU30-0125_10878_l_7801_cov_5.023496full | [IMGVR_UViG_3300005670_000031](https://img.jgi.doe.gov/cgi-bin/vr/main.cgi?section=ViralBrowse&page=uviginfo&uvig_id=IMGVR_UViG_3300005670_000031) | Duplodnaviria; Heunggongvirae; Uroviricota; Caudoviricetes; Caudovirales; ; ; ; | 4926/5007 98% | 0 |
| phage11_NTU30-0125 |  |  |  |  |  |
| phage12_NTU30-0125 | NTU30-0125_21134_l_4636_cov_2081.934949full | [IMGVR_UViG_3300013502_000003](https://img.jgi.doe.gov/cgi-bin/vr/main.cgi?section=ViralBrowse&page=uviginfo&uvig_id=IMGVR_UViG_3300013502_000003) | Duplodnaviria; Heunggongvirae; Uroviricota; Caudoviricetes; Caudovirales; ; ; ; | 2240/2331 96% | 0 |
| phage13_NTU30-0125 | NTU30-0125_4700_l_15250_cov_63.231787full | [IMGVR_UViG_3300005674_000088](https://img.jgi.doe.gov/cgi-bin/vr/main.cgi?section=ViralBrowse&page=uviginfo&uvig_id=IMGVR_UViG_3300005674_000088) | Duplodnaviria; Heunggongvirae; Uroviricota; Caudoviricetes; Caudovirales; Siphoviridae; ; ; | 3751/4735 79% | 0 |
| phage14_NTU30-0125 |  |  |  |  |  |
| phage15_NTU30-0515 | NTU30-0515_1769_l_28276_cov_53.226179full | [IMGVR_UViG_3300029931_000055](https://img.jgi.doe.gov/cgi-bin/vr/main.cgi?section=ViralBrowse&page=uviginfo&uvig_id=IMGVR_UViG_3300029931_000055) | Duplodnaviria; Heunggongvirae; Uroviricota; Caudoviricetes; Caudovirales; ; ; ; | 10135/11526 88% | 0 |
| phage16_NTU30-0515 | NTU30-0515_39562_l_2749_cov_2.687454full | [IMGVR_UViG_3300009873_000236](https://img.jgi.doe.gov/cgi-bin/vr/main.cgi?section=ViralBrowse&page=uviginfo&uvig_id=IMGVR_UViG_3300009873_000236) | Duplodnaviria; Heunggongvirae; Uroviricota; Caudoviricetes; Caudovirales; ; ; ; | 368/494 74% | 2.00E-45 |
| phage17_NTU30-0515 | NTU30-0515_8965_l_7317_cov_19.443542full | [IMGVR_UViG_3300005684_000011](https://img.jgi.doe.gov/cgi-bin/vr/main.cgi?section=ViralBrowse&page=uviginfo&uvig_id=IMGVR_UViG_3300005684_000011) | 1 | 855/1049 82% | 0 |
| phage18_NTU30-0515 | NTU30-0515_78245_l_1814_cov_5.552587full | [IMGVR_UViG_3300013800_000258](https://img.jgi.doe.gov/cgi-bin/vr/main.cgi?section=ViralBrowse&page=uviginfo&uvig_id=IMGVR_UViG_3300013800_000258) | Duplodnaviria; Heunggongvirae; Uroviricota; Caudoviricetes; Caudovirales; ; ; ; | 1766/1802 98% | 0 |
| phage19_NTU30-0802 | NTU30-0802_2033_l_32420_cov_76.289665full | [IMGVR_UViG_3300003281_000001](https://img.jgi.doe.gov/cgi-bin/vr/main.cgi?section=ViralBrowse&page=uviginfo&uvig_id=IMGVR_UViG_3300003281_000001) | Duplodnaviria; Heunggongvirae; Uroviricota; Caudoviricetes; Caudovirales; ; ; ; | 711/871 82% | 0 |
| phage15_NTU30-0802 | NTU30-0802_7229_l_11915_cov_9.216020full | [IMGVR_UViG_3300029931_000055](https://img.jgi.doe.gov/cgi-bin/vr/main.cgi?section=ViralBrowse&page=uviginfo&uvig_id=IMGVR_UViG_3300029931_000055) | Duplodnaviria; Heunggongvirae; Uroviricota; Caudoviricetes; Caudovirales; ; ; ; | 10135/11526 88% | 0 |
| phage20_NTU30-0802 |  |  |  |  |  |
| phage21_NTU30-0802 | NTU30-0802_34625_l_3627_cov_3.502800full | [IMGVR_UViG_3300013333_000005](https://img.jgi.doe.gov/cgi-bin/vr/main.cgi?section=ViralBrowse&page=uviginfo&uvig_id=IMGVR_UViG_3300013333_000005) |  | 1883/2145 88% | 0 |
| phage22_NTU30-0823 | NTU30-0823_2291_l_28150_cov_120.898096full | [IMGVR_UViG_3300003281_000001](https://img.jgi.doe.gov/cgi-bin/vr/main.cgi?section=ViralBrowse&page=uviginfo&uvig_id=IMGVR_UViG_3300003281_000001) | Duplodnaviria; Heunggongvirae; Uroviricota; Caudoviricetes; Caudovirales; ; ; ; | 711/871 82% | 0 |
| phage15_NTU30-0823 | NTU30-0823_5731_l_11915_cov_11.486594full | [IMGVR_UViG_3300029931_000055](https://img.jgi.doe.gov/cgi-bin/vr/main.cgi?section=ViralBrowse&page=uviginfo&uvig_id=IMGVR_UViG_3300029931_000055) | Duplodnaviria; Heunggongvirae; Uroviricota; Caudoviricetes; Caudovirales; ; ; ; | 10135/11526 88% | 0 |
| phage23_NTU30-0823 |  |  |  |  |  |
| phage24_NTU35-0823 | NTU35-0823_7386_l_9832_cov_8.902526full | [IMGVR_UViG_3300005664_000310](https://img.jgi.doe.gov/cgi-bin/vr/main.cgi?section=ViralBrowse&page=uviginfo&uvig_id=IMGVR_UViG_3300005664_000310) | Duplodnaviria; Heunggongvirae; Uroviricota; Caudoviricetes; Caudovirales; ; ; ; | 1585/1934 82% | 0 |

Table S7. Prophages identified from the *Ca.* Accumulibacter genomes

| **Number** | **Genomes** | **Name** | **Size** | **Total proteins** | **Score** | **Completeness** | **GC content** |
| --- | --- | --- | --- | --- | --- | --- | --- |
| 1 | 66-26 | 66-26_ppg1 | 9.5Kb | 10 | 20 | incomplete | 62.89% |
| 2 | 66-26 | 66-26_ppg2 | 14Kb | 11 | 20 | incomplete | 68.68% |
| 3 | 66-26 | 66-26_ppg3 | 28.5Kb | 10 | 20 | incomplete | 65.21% |
| 4 | 66-26 | 66-26_ppg4 | 8Kb | 8 | 10 | incomplete | 62.37% |
| 5 | 66-26 | 66-26_ppg5 | 8.2Kb | 7 | 30 | incomplete | 56.32% |
| 6 | 66-26 | 66-26_ppg6 | 9Kb | 8 | 10 | incomplete | 63.13% |
| 7 | AALB | AALB_ppg1 | 6.8Kb | 8 | 10 | incomplete | 61.51% |
| 8 | AALB | AALB_ppg2 | 10.3Kb | 11 | 30 | incomplete | 59.26% |
| 9 | BA-93 | BA-93_ppg1 | 12.6Kb | 13 | 40 | incomplete | 60.85% |
| 10 | BA-93 | BA-93_ppg2 | 40.9Kb | 36 | 80 | questionable | 62.17% |
| 11 | BAT3C415 | BAT3C415_ppg1 | 9.4Kb | 10 | 40 | incomplete | 62.30% |
| 12 | BAT3C415 | BAT3C415_ppg2 | 27.1Kb | 16 | 50 | incomplete | 61.30% |
| 13 | BATAC285 | BATAC285_ppg1 | 10.1Kb | 13 | 20 | incomplete | 64.22% |
| 14 | BATAC285 | BATAC285_ppg2 | 13.6Kb | 12 | 40 | incomplete | 63.92% |
| 15 | Bin19 | Bin19_ppg1 | 10.9Kb | 7 | 20 | incomplete | 62.12% |
| 16 | Bin19 | Bin19_ppg2 | 14.3Kb | 17 | 40 | incomplete | 61.44% |
| 17 | DS2011 | DS2011_ppg1 | 36.7Kb | 45 | 80 | questionable | 65.01% |
| 18 | MAG-196 | MAG-196_ppg1 | 4.8Kb | 7 | 60 | incomplete | 60.97% |
| 19 | MAG-203 | MAG-203_ppg1 | 8.6Kb | 12 | 30 | incomplete | 63.86% |
| 20 | MAXAC027 | MAXAC027_ppg1 | 13.7Kb | 25 | 50 | incomplete | 58.36% |
| 21 | MAXAC027 | MAXAC027_ppg2 | 12.3Kb | 17 | 30 | incomplete | 61.25% |
| 22 | MAXAC027 | MAXAC027_ppg3 | 8.2Kb | 21 | 30 | incomplete | 57.35% |
| 23 | MAXAC027 | MAXAC027_ppg4 | 20Kb | 24 | 20 | incomplete | 45.73% |
| 24 | MAXAC027 | MAXAC027_ppg5 | 9.8Kb | 13 | 30 | incomplete | 60.65% |
| 25 | MAXAC027 | MAXAC027_ppg6 | 15.7Kb | 17 | 50 | incomplete | 62.45% |
| 26 | MAXAC027 | MAXAC027_ppg7 | 17.1Kb | 25 | 120 | intact | 59.09% |
| 27 | MAXAC027 | MAXAC027_ppg8 | 9Kb | 16 | 20 | incomplete | 61.05% |
| 28 | MAXAC027 | MAXAC027_ppg9 | 20.3Kb | 28 | 20 | incomplete | 30.61% |
| 29 | MAXAC027 | MAXAC027_ppg10 | 7.3Kb | 16 | 40 | incomplete | 59.51% |
| 30 | SBRL | SBRL_ppg1 | 18.3Kb | 17 | 90 | questionable | 63.79% |
| 31 | SBRL | SBRL_ppg2 | 7.8Kb | 10 | 40 | incomplete | 59.44% |
| 32 | SBRL | SBRL_ppg3 | 27.9Kb | 34 | 50 | incomplete | 64.63% |
| 33 | SBRL | SBRL_ppg4 | 20.2Kb | 9 | 10 | incomplete | 61.74% |
| 34 | SBRS | SBRS_ppg1 | 8.3Kb | 8 | 30 | incomplete | 57.56% |
| 35 | SBRS | SBRS_ppg2 | 49.2Kb | 48 | 120 | intact | 62.15% |
| 36 | SBRS | SBRS_ppg3 | 26.4Kb | 17 | 30 | incomplete | 62.77% |
| 37 | SBRS | SBRS_ppg4 | 11.4Kb | 11 | 50 | incomplete | 60.34% |
| 38 | SCELSE-1 | SCELSE-1_ppg1 | 8.4Kb | 8 | 50 | incomplete | 63.15% |
| 39 | SCELSE-1 | SCELSE-1_ppg2 | 6.8Kb | 7 | 10 | incomplete | 61.90% |
| 40 | SCN18 | SCN18_ppg1 | 8Kb | 8 | 10 | incomplete | 62.37% |
| 41 | SCN18 | SCN18_ppg2 | 28.5Kb | 10 | 20 | incomplete | 65.21% |
| 42 | SCN18 | SCN18_ppg3 | 8.8Kb | 8 | 10 | incomplete | 63.21% |
| 43 | SCN18 | SCN18_ppg4 | 9.5Kb | 10 | 20 | incomplete | 62.89% |
| 44 | SCN18 | SCN18_ppg5 | 5.9Kb | 7 | 10 | incomplete | 60.23% |
| 45 | SK-02 | SK-02_ppg1 | 7.9Kb | 13 | 30 | incomplete | 61.45% |
| 46 | SK-02 | SK-02_ppg2 | 17.1Kb | 16 | 100 | intact | 63.71% |
| 47 | SSA1 | SSA1_ppg1 | 20Kb | 19 | 90 | questionable | 64.66% |
| 48 | SSA1 | SSA1_ppg2 | 45.7Kb | 55 | 110 | intact | 65.95% |
| 49 | SSA1 | SSA1_ppg3 | 8.8Kb | 11 | 50 | incomplete | 59.21% |
| 50 | SSA1 | SSA1_ppg4 | 14.3Kb | 17 | 40 | incomplete | 61.45% |
| 51 | SSA1 | SSA1_ppg5 | 8.8Kb | 11 | 60 | incomplete | 59.12% |
| 52 | SSB1 | SSB1_ppg1 | 25.9Kb | 29 | 120 | intact | 66.28% |
| 53 | SSB1 | SSB1_ppg2 | 24.4Kb | 26 | 150 | intact | 67.18% |
| 54 | SSB1 | SSB1_ppg3 | 13.1Kb | 12 | 40 | incomplete | 66.15% |
| 55 | SSB1 | SSB1_ppg4 | 9.4Kb | 8 | 20 | incomplete | 66.14% |
| 56 | SSB1 | SSB1_ppg5 | 25Kb | 10 | 10 | incomplete | 67.04% |
| 57 | UW1 | UW1_ppg1 | 11.1Kb | 10 | 60 | incomplete | 63.50% |
| 58 | UW1 | UW1_ppg2 | 12.3Kb | 15 | 20 | incomplete | 60.58% |
| 59 | UW5 | UW5_ppg1 | 10.1Kb | 8 | 20 | incomplete | 63.03% |
| 60 | UW5 | UW5_ppg2 | 7.8Kb | 7 | 10 | incomplete | 65.66% |
| 61 | UW5 | UW5_ppg3 | 7.3Kb | 8 | 30 | incomplete | 64.00% |
| 62 | UW6 | UW6_ppg1 | 24.2Kb | 15 | 30 | incomplete | 61.48% |
| 63 | UW6 | UW6_ppg2 | 25Kb | 22 | 30 | incomplete | 63.77% |
| 64 | UW6 | UW6_ppg3 | 27.2Kb | 15 | 20 | incomplete | 63.44% |
| 65 | UW7 | UW7_ppg1 | 10.1Kb | 9 | 10 | incomplete | 62.19% |
| 66 | UW7 | UW7_ppg2 | 8.4Kb | 8 | 50 | incomplete | 63.29% |
| 67 | UW7 | UW7_ppg3 | 7.4Kb | 9 | 10 | incomplete | 66.39% |
| 68 | UW8 | UW8_ppg1 | 8.6Kb | 11 | 30 | incomplete | 63.87% |
| 69 | UW8 | UW8_ppg2 | 23.3Kb | 11 | 50 | incomplete | 60.61% |
| 70 | UW9 | UW9_ppg1 | 7.3Kb | 8 | 30 | incomplete | 64.00% |
| 71 | UW9 | UW9_ppg2 | 10.1Kb | 8 | 20 | incomplete | 63.03% |
| 72 | UW9 | UW9_ppg3 | 7.8Kb | 7 | 10 | incomplete | 65.65% |
| 73 | UW12 | UW12_ppg1 | 8.8Kb | 8 | 10 | incomplete | 57.05% |
| 74 | UW13 | UW13_ppg1 | 41.8Kb | 53 | 150 | intact | 65.24% |
| 75 | UWLDOIC | UWLDOIC_ppg1 | 16Kb | 17 | 20 | incomplete | 62.70% |
| 76 | SCELSE-7IIH | SCELSE-7IIH_ppg1 | 38.1Kb | 54 | 90 | questionable | 64.94% |
| 77 | SCELSE-6IIF | SCELSE-6IIF_ppg1 | 8.4Kb | 8 | 50 | incomplete | 63.67% |
| 78 | SCELSE-6IIF | SCELSE-6IIF_ppg2 | 9.9Kb | 9 | 10 | incomplete | 62.93% |
| 79 | SCELSE-6IIF | SCELSE-6IIF_ppg3 | 8.1Kb | 6 | 10 | incomplete | 67.11% |
| 80 | SCELSE-8IIC | SCELSE-8IIC_ppg1 | 7.8Kb | 9 | 10 | incomplete | 59.13% |
| 81 | SCELSE-8IIC | SCELSE-8IIC_ppg2 | 11.3Kb | 10 | 50 | incomplete | 57.40% |
| 82 | SCELSE-8IIC | SCELSE-8IIC_ppg3 | 10.8Kb | 14 | 10 | incomplete | 59.54% |
| 83 | SCELSE-8IIC | SCELSE-8IIC_ppg4 | 9.2Kb | 8 | 30 | incomplete | 61.56% |
| 84 | SCELSE-8IIC | SCELSE-8IIC_ppg5 | 7.5Kb | 12 | 60 | incomplete | 63.81% |
| 85 | SCELSE-5IIH | SCELSE-5IIH_ppg1 | 38.1Kb | 54 | 90 | questionable | 64.94% |
| 86 | SCELSE-4IIF | SCELSE-4IIF _ppg1 | 8.4Kb | 8 | 50 | incomplete | 63.67% |
| 87 | SCELSE-4IIF | SCELSE-4IIF _ppg2 | 9.9Kb | 9 | 10 | incomplete | 62.93% |
| 88 | SCELSE-4IIF | SCELSE-4IIF _ppg3 | 8.1Kb | 6 | 10 | incomplete | 67.11% |
| 89 | SCELSE-3IIF | SCELSE-3IIF _ppg1 | 8.4Kb | 8 | 50 | incomplete | 63.67% |
| 90 | SCELSE-3IIF | SCELSE-3IIF _ppg2 | 9.9Kb | 9 | 10 | incomplete | 62.93% |
| 91 | SCELSE-3IIF | SCELSE-3IIF _ppg3 | 8.1Kb | 6 | 10 | incomplete | 67.11% |
| 92 | SCELSE-10IIF | SCELSE-10IIF _ppg1 | 35.3Kb | 29 | 100 | intact | 65.50% |
| 93 | SCELSE-10IIF | SCELSE-10IIF _ppg2 | 17.6Kb | 17 | 30 | incomplete | 62.86% |
| 94 | SCELSE-10IIF | SCELSE-10IIF _ppg3 | 9.6Kb | 9 | 40 | incomplete | 69.15% |
| 95 | SCELSE-2IIC | SCELSE-2IIC_ppg1 | 10.2Kb | 8 | 10 | incomplete | 59.77% |
| 96 | SCELSE-2IIC | SCELSE-2IIC_ppg2 | 11.3Kb | 10 | 50 | incomplete | 57.40% |
| 97 | SCELSE-2IIC | SCELSE-2IIC_ppg3 | 10.1Kb | 14 | 80 | questionable | 60.09% |
| 98 | SCELSE-9IIF | SCELSE-9IIF_ppg1 | 18.3Kb | 24 | 90 | questionable | 66.58% |
| 99 | SCELSE-9IIF | SCELSE-9IIF_ppg2 | 29.1Kb | 15 | 30 | incomplete | 64.17% |
| 100 | SCELSE-9IIF | SCELSE-9IIF_ppg3 | 9.6Kb | 9 | 40 | incomplete | 69.15% |
| 101 | ACC003 | ACC003_ppg1 | 36.2Kb | 27 | 120 | intact | 62.35% |
| 102 | ACC003 | ACC003_ppg2 | 11.3Kb | 12 | 80 | questionable | 62.28% |
| 103 | ACC003 | ACC003_ppg3 | 15.5Kb | 19 | 20 | incomplete | 63.83% |
| 104 | ACC005 | ACC005_ppg1 | 36.7Kb | 36 | 80 | questionable | 61.98% |
| 105 | ACC005 | ACC005_ppg2 | 9.4Kb | 10 | 80 | questionable | 61.23% |
| 106 | ACC005 | ACC005_ppg3 | 19.1Kb | 17 | 50 | incomplete | 61.71% |
| 107 | ACC005 | ACC005_ppg4 | 6.8Kb | 7 | 70 | questionable | 61.15% |
| 108 | ACC005 | ACC005_ppg5 | 31Kb | 10 | 50 | incomplete | 58.64% |
| 109 | ACC005 | ACC005_ppg6 | 23Kb | 25 | 60 | incomplete | 59.57% |
| 110 | ACC005 | ACC005_ppg7 | 11.7Kb | 11 | 50 | incomplete | 61.52% |
| 111 | ACC007 | ACC007_ppg1 | 17.8Kb | 10 | 10 | incomplete | 63.70% |
| 112 | ACC007 | ACC007_ppg2 | 11.8Kb | 13 | 90 | questionable | 64.82% |
| 113 | ACC007 | ACC007_ppg3 | 8.1Kb | 10 | 30 | incomplete | 61.44% |
| 114 | ACC007 | ACC007_ppg4 | 20Kb | 18 | 70 | questionable | 59.04% |
| 115 | ACC007 | ACC007_ppg5 | 8.5Kb | 8 | 70 | questionable | 62.76% |
| 116 | ACC007 | ACC007_ppg6 | 35.3Kb | 26 | 60 | incomplete | 62.25% |
| 117 | ACC007 | ACC007_ppg7 | 21.6Kb | 25 | 70 | questionable | 64.25% |
| 118 | ACC012 | ACC012_ppg1 | 12.7Kb | 12 | 30 | incomplete | 60.03% |
| 119 | ACC012 | ACC012_ppg2 | 9.9Kb | 12 | 40 | incomplete | 61.50% |
| 120 | ACC012 | ACC012_ppg3 | 28.2Kb | 15 | 20 | incomplete | 63.02% |
| 121 | ACC012 | ACC012_ppg4 | 19.9Kb | 23 | 60 | incomplete | 62.98% |
| 122 | ACC012 | ACC012_ppg5 | 9.4Kb | 7 | 60 | incomplete | 61.23% |
| 123 | ACC012 | ACC012_ppg6 | 18.9Kb | 17 | 60 | incomplete | 59.87% |
| 124 | ACC012 | ACC012_ppg7 | 34.5Kb | 28 | 90 | questionable | 60.91% |
| 125 | SCUT-1 | SCUT-1_ppg1 | 7.6Kb | 7 | 10 | incomplete | 64.91% |
| 126 | SCUT-1 | SCUT-1_ppg2 | 11.3Kb | 10 | 50 | incomplete | 57.45% |
| 127 | UBA6585 | UBA6585_ppg1 | 13.2Kb | 19 | 50 | incomplete | 67.26% |
| 128 | UBA6585 | UBA6585_ppg2 | 14.4Kb | 15 | 30 | incomplete | 64.08% |
| 129 | UBA6658 | UBA6658_ppg1 | 36.7Kb | 31 | 80 | questionable | 61.98% |
| 130 | SCUT-2 | SCUT-2_ppg1 | 7.6Kb | 6 | 10 | incomplete | 64.96% |
| 131 | SCUT-2 | SCUT-2_ppg2 | 10.2Kb | 8 | 10 | incomplete | 59.76% |
| 132 | SCUT-2 | SCUT-2_ppg3 | 13.5Kb | 12 | 60 | incomplete | 64.38% |
| 133 | SCUT-2 | SCUT-2_ppg4 | 11.3Kb | 10 | 50 | incomplete | 57.43% |

Table S8. Prophage identification results by using the IMG/VR database

| **Prophage region** | **UViGs** | **Lineage** | **Genome Name** | **E-value** | **Identities** |
| --- | --- | --- | --- | --- | --- |
| BA-93_ppg2 | [IMGVR_UViG_3300026283_000026](https://img.jgi.doe.gov/cgi-bin/vr/main.cgi?section=ViralBrowse&page=uviginfo&uvig_id=IMGVR_UViG_3300026283_000026) | Duplodnaviria; Heunggongvirae; Uroviricota; Caudoviricetes; Caudovirales; Siphoviridae; ; ; | Wastewater treatment Type I Accumulibacter community from EBPR Bioreactor in Madison, WI, USA - Reactor 1_4/24/2008_ DNA (SPAdes) | 0 | 8680/8680 100% |
| DS2011_ppg1 | [IMGVR_UViG_3300035670_002386](https://img.jgi.doe.gov/cgi-bin/vr/main.cgi?section=ViralBrowse&page=uviginfo&uvig_id=IMGVR_UViG_3300035670_002386) | Duplodnaviria; Heunggongvirae; Uroviricota; Caudoviricetes; Caudovirales; ; ; ; | Fracking water microbial communities from deep shales in Oklahoma, United States - MC-6-LG | 0 | 4005/4204 95% |
| MAXAC027_ppg7 | [IMGVR_UViG_3300009767_000039](https://img.jgi.doe.gov/cgi-bin/vr/main.cgi?section=ViralBrowse&page=uviginfo&uvig_id=IMGVR_UViG_3300009767_000039) | Duplodnaviria; Heunggongvirae; Uroviricota; Caudoviricetes; Caudovirales; ; ; ; | Active sludge microbial communities of municipal wastewater-treating anaerobic digesters from Japan - AD_JPNAS3_MetaG | 0 | 2394/3016 79% |
| SBRL_ppg1 | [IMGVR_UViG_3300026299_000013](https://img.jgi.doe.gov/cgi-bin/vr/main.cgi?section=ViralBrowse&page=uviginfo&uvig_id=IMGVR_UViG_3300026299_000013) |  | Wastewater treatment Type I Accumulibacter community from EBPR Bioreactor in Madison, WI, USA - Reactor 1_9/17/2007_ DNA (SPAdes) | 0 | 6785/8428 81% |
| SBRS_ppg2 | [IMGVR_UViG_2022004001_000001](https://img.jgi.doe.gov/cgi-bin/vr/main.cgi?section=ViralBrowse&page=uviginfo&uvig_id=IMGVR_UViG_2022004001_000001) | Duplodnaviria; Heunggongvirae; Uroviricota; Caudoviricetes; Caudovirales; ; ; ; | Wastewater treatment Type I Accumulibacter community from EBPR Bioreactor in Madison, WI - Type I | 0 | 13517/13855 98% |
| SK-02_ppg2 | [IMGVR_UViG_3300026299_000013](https://img.jgi.doe.gov/cgi-bin/vr/main.cgi?section=ViralBrowse&page=uviginfo&uvig_id=IMGVR_UViG_3300026299_000013) |  | Wastewater treatment Type I Accumulibacter community from EBPR Bioreactor in Madison, WI, USA - Reactor 1_9/17/2007_ DNA (SPAdes) | 0 | 6776/8447 80% |
| SSA1_ppg1 | [IMGVR_UViG_3300026299_000013](https://img.jgi.doe.gov/cgi-bin/vr/main.cgi?section=ViralBrowse&page=uviginfo&uvig_id=IMGVR_UViG_3300026299_000013) |  | Wastewater treatment Type I Accumulibacter community from EBPR Bioreactor in Madison, WI, USA - Reactor 1_9/17/2007_ DNA (SPAdes) | 0 | 6775/8447 80% |
| SSA1_ppg2 | [IMGVR_UViG_3300005684_000053](https://img.jgi.doe.gov/cgi-bin/vr/main.cgi?section=ViralBrowse&page=uviginfo&uvig_id=IMGVR_UViG_3300005684_000053) | Duplodnaviria; Heunggongvirae; Uroviricota; Caudoviricetes; Caudovirales; ; ; ; | Enhanced biological phosphorus removal bioreactor viral communities from the University of Queensland, Australia - SBR4-V91805 Phage Sequencing | 0 | 1100/1278 86% |
| SSB1_ppg1 | [IMGVR_UViG_3300013502_000014](https://img.jgi.doe.gov/cgi-bin/vr/main.cgi?section=ViralBrowse&page=uviginfo&uvig_id=IMGVR_UViG_3300013502_000014) | Duplodnaviria; Heunggongvirae; Uroviricota; Caudoviricetes; Caudovirales; Siphoviridae; ; ; | Activated sludge bacterial and viral communities from EBPR bioreactors in Brisbane, Australia - M81612 | 0 | 15174/16263 93% |
| SSB1_ppg2 | [IMGVR_UViG_3300013502_000014](https://img.jgi.doe.gov/cgi-bin/vr/main.cgi?section=ViralBrowse&page=uviginfo&uvig_id=IMGVR_UViG_3300013502_000014) | Duplodnaviria; Heunggongvirae; Uroviricota; Caudoviricetes; Caudovirales; Siphoviridae; ; ; | Activated sludge bacterial and viral communities from EBPR bioreactors in Brisbane, Australia - M81612 | 0 | 14951/16155 93% |
| UW13_ppg1 | [IMGVR_UViG_3300005684_000046](https://img.jgi.doe.gov/cgi-bin/vr/main.cgi?section=ViralBrowse&page=uviginfo&uvig_id=IMGVR_UViG_3300005684_000046) | Duplodnaviria; Heunggongvirae; Uroviricota; Caudoviricetes; Caudovirales; ; ; ; | Enhanced biological phosphorus removal bioreactor viral communities from the University of Queensland, Australia - SBR4-V91805 Phage Sequencing | 0 | 17188/17365 99% |
| SCELSE-7IIH_ppg1 | [IMGVR_UViG_3300009653_000060](https://img.jgi.doe.gov/cgi-bin/vr/main.cgi?section=ViralBrowse&page=uviginfo&uvig_id=IMGVR_UViG_3300009653_000060) | Duplodnaviria; Heunggongvirae; Uroviricota; Caudoviricetes; Caudovirales; ; ; ; | Active sludge microbial communities of municipal wastewater-treating anaerobic digesters from Canada - AD_UKC130_MetaG | 0 | 1390/1821 76% |
| SCELSE-5IIH_ppg1 | [IMGVR_UViG_3300009653_000060](https://img.jgi.doe.gov/cgi-bin/m/main.cgi?section=ViralBrowse&page=uviginfo&uvig_id=IMGVR_UViG_3300009653_000060) | Duplodnaviria; Heunggongvirae; Uroviricota; Caudoviricetes; Caudovirales; ; ; ; | Active sludge microbial communities of municipal wastewater-treating anaerobic digesters from Canada - AD_UKC130_MetaG | 0 | 1390/1821 76% |
| SCELSE-10IIF _ppg1 | [IMGVR_UViG_3300005102_000012](https://img.jgi.doe.gov/cgi-bin/m/main.cgi?section=ViralBrowse&page=uviginfo&uvig_id=IMGVR_UViG_3300005102_000012) | Duplodnaviria; Heunggongvirae; Uroviricota; Caudoviricetes; Caudovirales; ; ; ; | Wastewater treatment Type I Accumulibacter community from EBPR Bioreactor in Madison, WI, USA - Reactor 1_1/10/2011_ DNA | 0 | 1864/2257 83% |
| SCELSE-2IIC_ppg3 | [IMGVR_UViG_3300013771_000016](https://img.jgi.doe.gov/cgi-bin/m/main.cgi?section=ViralBrowse&page=uviginfo&uvig_id=IMGVR_UViG_3300013771_000016) | Duplodnaviria; Heunggongvirae; Uroviricota; Caudoviricetes; Caudovirales; Siphoviridae; ; ; | Activated sludge bacterial and viral communities from EBPR bioreactors in Brisbane, Australia - M92206 | 0 | 7114/7442 96% |
| SCELSE-9IIF_ppg1 | [IMGVR_UViG_3300001712_000001](https://img.jgi.doe.gov/cgi-bin/m/main.cgi?section=ViralBrowse&page=uviginfo&uvig_id=IMGVR_UViG_3300001712_000001) | Duplodnaviria; Heunggongvirae; Uroviricota; Caudoviricetes; Caudovirales; ; ; ; | Marine viral communities from the Pacific Ocean - LP-31 | 0 | 1232/1517 81% |
| ACC003_1 | [IMGVR_UViG_3300025292_000004](https://img.jgi.doe.gov/cgi-bin/m/main.cgi?section=ViralBrowse&page=uviginfo&uvig_id=IMGVR_UViG_3300025292_000004) | Duplodnaviria; Heunggongvirae; Uroviricota; Caudoviricetes; Caudovirales; ; ; ; | Arabidopsis root microbial communities from North Carolina, USA - plate scrape MF_Cvi_mLB_r2 (SPAdes) | 0 | 1982/2713 73% |
| ACC003_2 | [IMGVR_UViG_3300005683_000242](https://img.jgi.doe.gov/cgi-bin/m/main.cgi?section=ViralBrowse&page=uviginfo&uvig_id=IMGVR_UViG_3300005683_000242) |  | Enhanced biological phosphorus removal bioreactor viral communities from the University of Queensland, Australia - SBR4-V91802 Phage Sequencing | 3.00E-164 | 649/809 80% |
| ACC005_ppg1 | [IMGVR_UViG_3300003767_000006](https://img.jgi.doe.gov/cgi-bin/m/main.cgi?section=ViralBrowse&page=uviginfo&uvig_id=IMGVR_UViG_3300003767_000006) | Duplodnaviria; Heunggongvirae; Uroviricota; Caudoviricetes; Caudovirales; ; ; ; | Wastewater treatment Type I Accumulibacter community from EBPR Bioreactor in Madison, WI, USA - Reactor 1_10/4/2010_ DNA | 0 | 5972/6783 88% |
| ACC005_ppg2 |  |  |  |  |  |
| ACC005_ppg4 | [IMGVR_UViG_3300031417_000051](https://img.jgi.doe.gov/cgi-bin/m/main.cgi?section=ViralBrowse&page=uviginfo&uvig_id=IMGVR_UViG_3300031417_000051) | Duplodnaviria; Heunggongvirae; Uroviricota; Caudoviricetes; Caudovirales; ; ; ; | Phyllosphere microbial communities from UC Gill Tract Community Farm, Albany, California, United States - DLSLSB.R2 | 6.00E-55 | 1132/1614 70% |
| ACC007_ppg2 | [IMGVR_UViG_3300005988_000029](https://img.jgi.doe.gov/cgi-bin/m/main.cgi?section=ViralBrowse&page=uviginfo&uvig_id=IMGVR_UViG_3300005988_000029) |  | Wastewater effluent complex algal communities from Wisconsin, to seasonally profile nutrient transformation and Carbon sequestration - JI 9/18/14 C2 DNA | 0 | 1021/1055 97% |
| ACC007_ppg4 | [IMGVR_UViG_3300003778_000016](https://img.jgi.doe.gov/cgi-bin/m/main.cgi?section=ViralBrowse&page=uviginfo&uvig_id=IMGVR_UViG_3300003778_000016) | Duplodnaviria; Heunggongvirae; Uroviricota; Caudoviricetes; Caudovirales; Siphoviridae; ; ; | Wastewater treatment Type I Accumulibacter community from EBPR Bioreactor in Madison, WI, USA - Reactor 1_9/17/2007_ DNA | 0 | 5831/7135 82% |
| ACC007_ppg5 |  |  |  |  |  |
| ACC007_ppg7 | [IMGVR_UViG_3300003767_000006](https://img.jgi.doe.gov/cgi-bin/m/main.cgi?section=ViralBrowse&page=uviginfo&uvig_id=IMGVR_UViG_3300003767_000006) | Duplodnaviria; Heunggongvirae; Uroviricota; Caudoviricetes; Caudovirales; ; ; ; | Wastewater treatment Type I Accumulibacter community from EBPR Bioreactor in Madison, WI, USA - Reactor 1_10/4/2010_ DNA | 0 | 10927/11753 93% |
| ACC012_ppg7 | [IMGVR_UViG_3300000053_000013](https://img.jgi.doe.gov/cgi-bin/m/main.cgi?section=ViralBrowse&page=uviginfo&uvig_id=IMGVR_UViG_3300000053_000013) |  | Coal bed methane well microbial communities from Alberta, Canada | 0 | 907/1073 85% |
| UBA6658_ppg1 | [IMGVR_UViG_3300003767_000006](https://img.jgi.doe.gov/cgi-bin/m/main.cgi?section=ViralBrowse&page=uviginfo&uvig_id=IMGVR_UViG_3300003767_000006) | Duplodnaviria; Heunggongvirae; Uroviricota; Caudoviricetes; Caudovirales; ; ; ; | Wastewater treatment Type I Accumulibacter community from EBPR Bioreactor in Madison, WI, USA - Reactor 1_10/4/2010_ DNA | 0 | 5972/6783 88% |
| ACC005_ppg4 | [IMGVR_UViG_3300031417_000051](https://img.jgi.doe.gov/cgi-bin/m/main.cgi?section=ViralBrowse&page=uviginfo&uvig_id=IMGVR_UViG_3300031417_000051) | Duplodnaviria; Heunggongvirae; Uroviricota; Caudoviricetes; Caudovirales; ; ; ; | Phyllosphere microbial communities from UC Gill Tract Community Farm, Albany, California, United States - DLSLSB.R2 | 6.00E-55 | 1132/1614 70% |
| ACC007_ppg2 | [IMGVR_UViG_3300005988_000029](https://img.jgi.doe.gov/cgi-bin/m/main.cgi?section=ViralBrowse&page=uviginfo&uvig_id=IMGVR_UViG_3300005988_000029) |  | Wastewater effluent complex algal communities from Wisconsin, to seasonally profile nutrient transformation and Carbon sequestration - JI 9/18/14 C2 DNA | 0 | 1021/1055 97% |
| ACC007_ppg4 | [IMGVR_UViG_3300003778_000016](https://img.jgi.doe.gov/cgi-bin/m/main.cgi?section=ViralBrowse&page=uviginfo&uvig_id=IMGVR_UViG_3300003778_000016) | Duplodnaviria; Heunggongvirae; Uroviricota; Caudoviricetes; Caudovirales; Siphoviridae; ; ; | Wastewater treatment Type I Accumulibacter community from EBPR Bioreactor in Madison, WI, USA - Reactor 1_9/17/2007_ DNA | 0 | 5831/7135 82% |
| ACC007_ppg5 |  |  |  |  |  |
| ACC007_ppg7 | [IMGVR_UViG_3300003767_000006](https://img.jgi.doe.gov/cgi-bin/m/main.cgi?section=ViralBrowse&page=uviginfo&uvig_id=IMGVR_UViG_3300003767_000006) | Duplodnaviria; Heunggongvirae; Uroviricota; Caudoviricetes; Caudovirales; ; ; ; | Wastewater treatment Type I Accumulibacter community from EBPR Bioreactor in Madison, WI, USA - Reactor 1_10/4/2010_ DNA | 0 | 10927/11753 93% |
| ACC012_ppg7 | [IMGVR_UViG_3300000053_000013](https://img.jgi.doe.gov/cgi-bin/m/main.cgi?section=ViralBrowse&page=uviginfo&uvig_id=IMGVR_UViG_3300000053_000013) |  | Coal bed methane well microbial communities from Alberta, Canada | 0 | 907/1073 85% |
| UBA6658_ppg1 | [IMGVR_UViG_3300003767_000006](https://img.jgi.doe.gov/cgi-bin/m/main.cgi?section=ViralBrowse&page=uviginfo&uvig_id=IMGVR_UViG_3300003767_000006) | Duplodnaviria; Heunggongvirae; Uroviricota; Caudoviricetes; Caudovirales; ; ; ; | Wastewater treatment Type I Accumulibacter community from EBPR Bioreactor in Madison, WI, USA - Reactor 1_10/4/2010_ DNA | 0 | 5972/6783 88% |

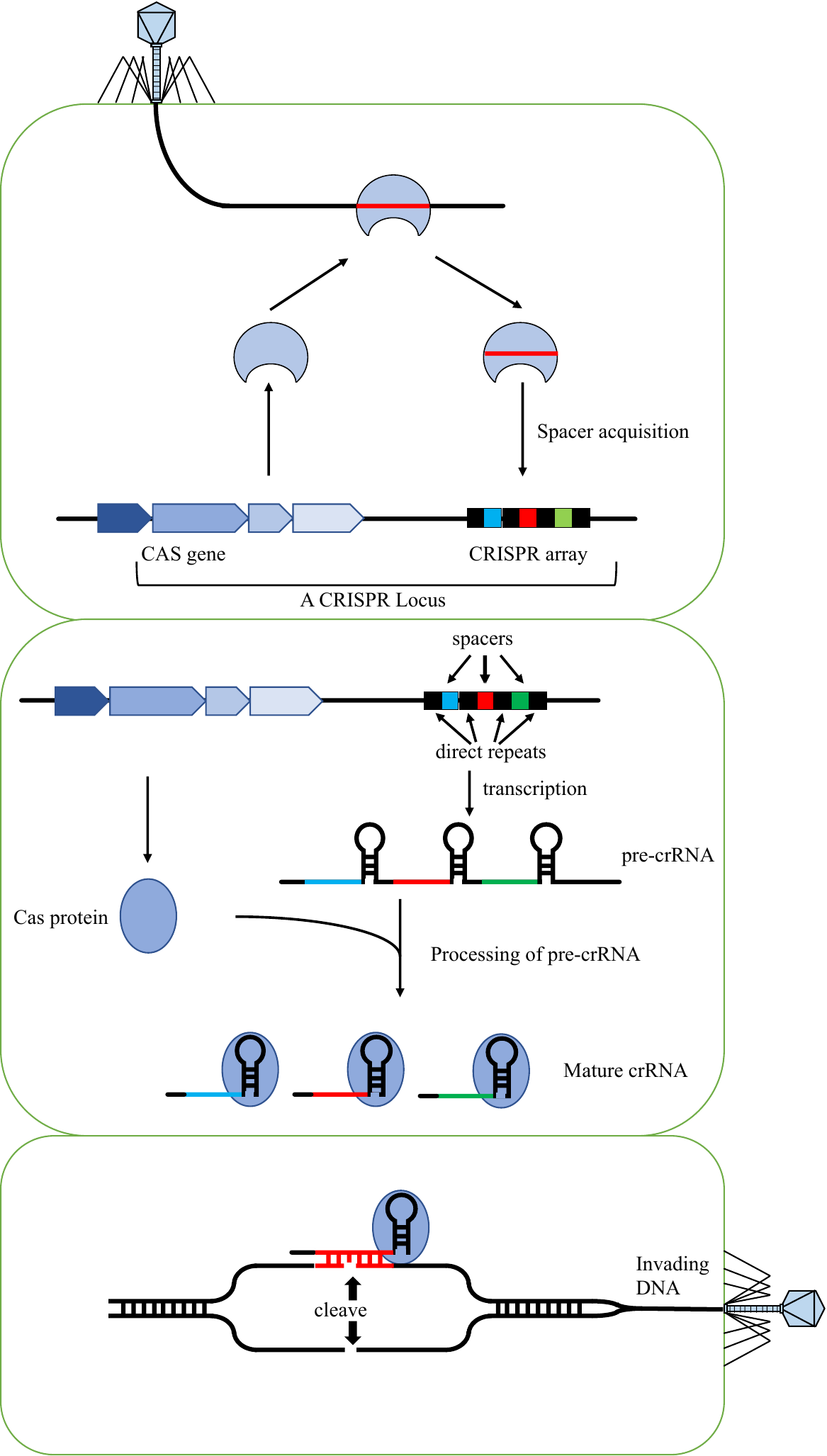

**Figure S1.** The anti-phage mechanism of the CRISPR-Cas system. Three squares represent three stages of the CRISPR-Cas system as adaption, CRISPR RNA (crRNA) biogenesis, and interference. At the adaption stage, the CRISPR-Cas system captures foreign nucleic acid (such as phage genetic material and plasmid) and integrates a fragment into the CRISPR array as spacers to provide infection memory. At the crRNA biogenesis stage, CRISPR repeat-spacer arrays are transcribed and processed into crRNAs. each crRNA contains a conserved direct repeat (DR) and a variable spacer. In the infection stage, the interference machinery is guided by crRNA to identify and cleave foreign nucleic acids.

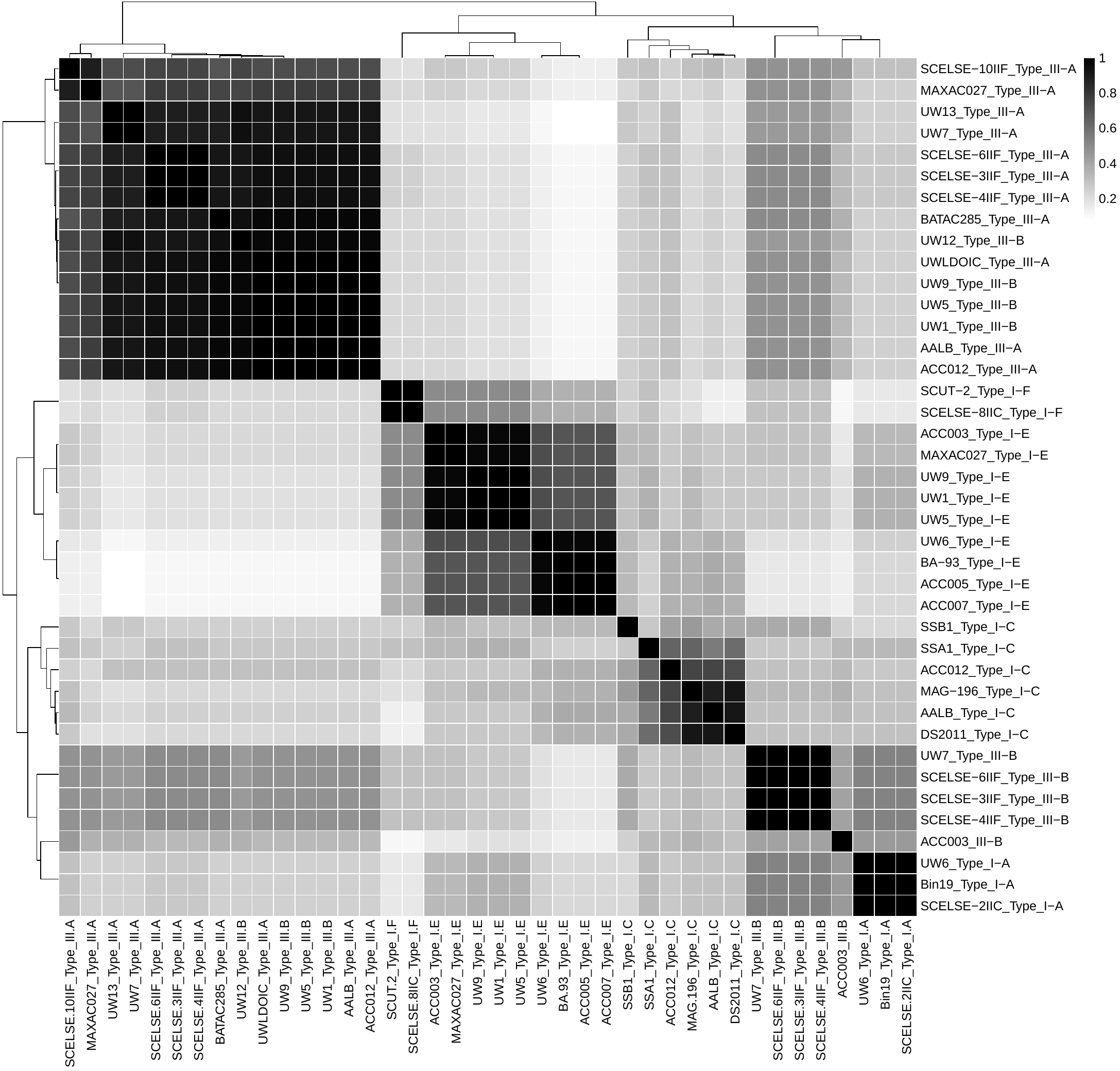

**Figure S2**. Sequence percent identity between each two DRs in all CRISPR loci.

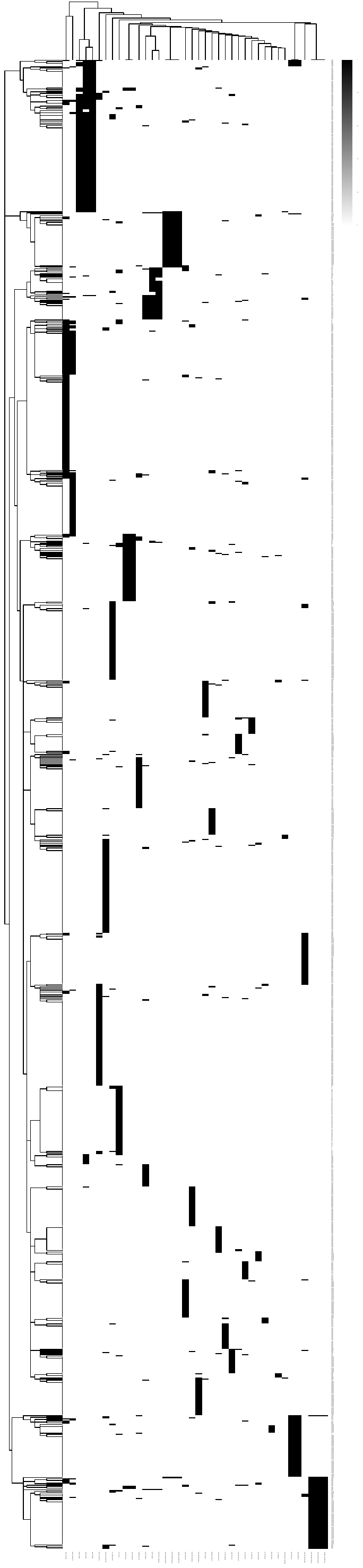

**Figure S3.** Relationships among spacers from all CRISPR-Cas loci in *Ca.* Accumulibacter.

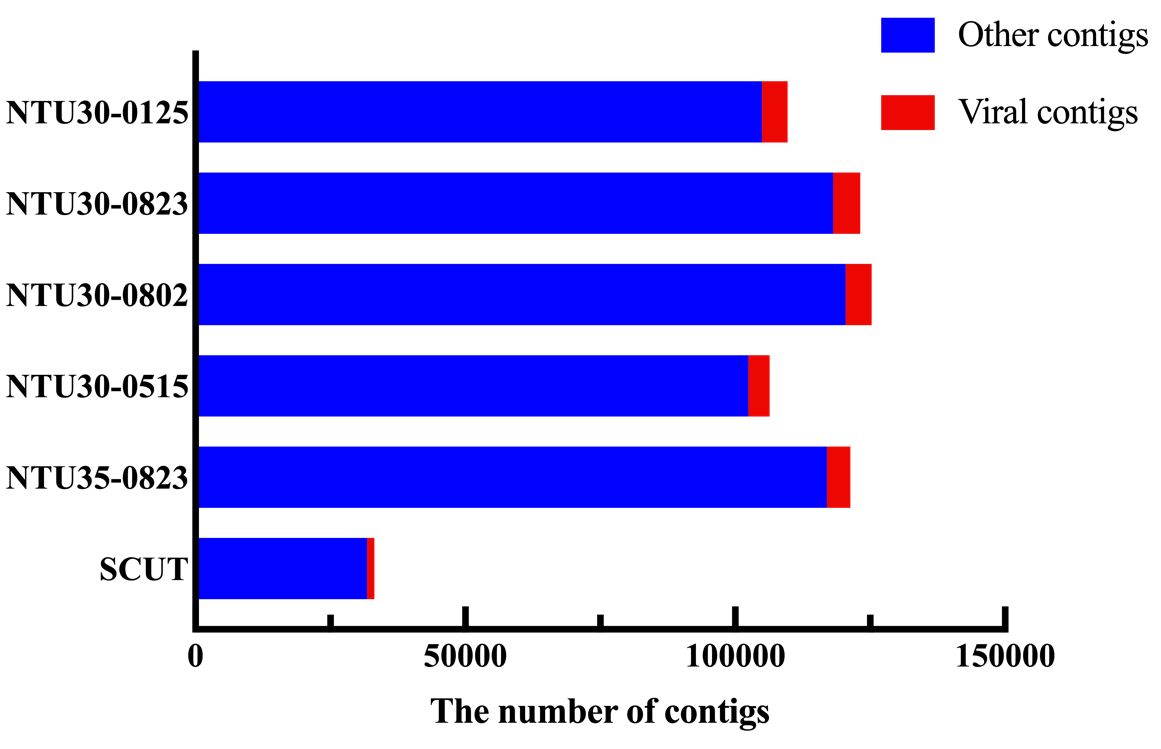

**Figure S4.** VirSorter (Guo et al., 2021) identified results of the 6 metagenomic datasets from our lab-scale reactors. Red bar indicates the viral contigs; blue bar indicates other contigs.

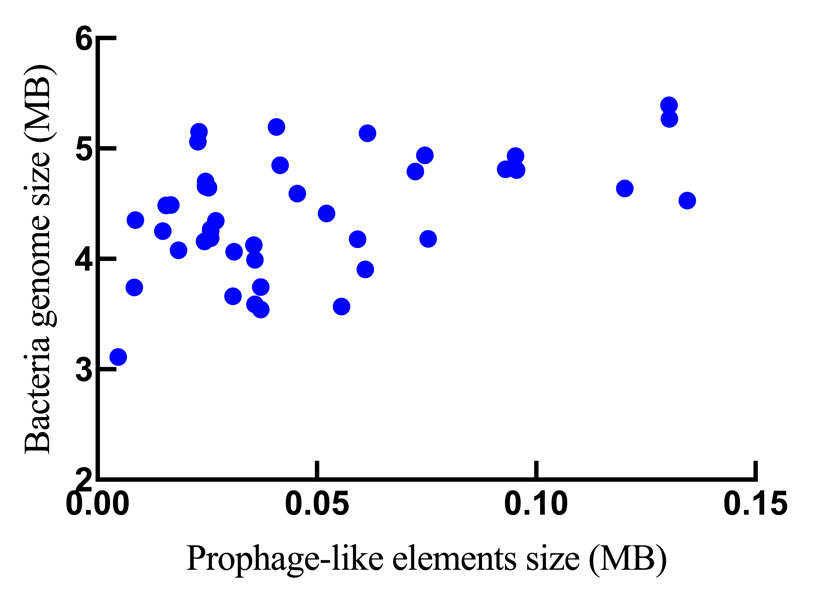

**Figure S5.** The relationship between the size of *Ca.* Accumulibacter genomes and that of the prophage-like elements.
